# High-resolution spatial mapping of cell state and lineage dynamics *in vivo* with PEtracer

**DOI:** 10.1101/2025.06.15.659774

**Authors:** Luke W. Koblan, Kathryn E. Yost, Pu Zheng, William N. Colgan, Matthew G. Jones, Dian Yang, Arhan Kumar, Jaspreet Sandhu, Alexandra Schnell, Dawei Sun, Can Ergen, Reuben A. Saunders, Xiaowei Zhuang, William E. Allen, Nir Yosef, Jonathan S. Weissman

## Abstract

Charting the spatiotemporal dynamics of cell fate determination in development and disease is a long-standing objective in biology. Here we present the design, development, and extensive validation of PEtracer, a prime editing-based, evolving lineage tracing technology compatible with both single-cell sequencing and multimodal imaging methodologies to jointly profile cell state and lineage in dissociated cells or while preserving cellular context in tissues with high spatial resolution. Using PEtracer coupled with MERFISH spatial transcriptomic profiling in a syngeneic mouse model of tumor metastasis, we reconstruct the growth of individually-seeded tumors *in vivo* and uncover distinct modules of cell-intrinsic and cell-extrinsic factors that coordinate tumor growth. More generally, PEtracer enables systematic characterization of cell state and lineage relationships in intact tissues over biologically-relevant temporal and spatial scales.

## Main

Defining the history of cell division, differentiation, movement, and death as well as the coordination of these processes in both time and space provides fundamental insights into how complex structures emerge in multicellular life (*1–3*). The comprehensive fate mapping of *C. elegans* produced critical insights into the dynamic cell state, lineage, and spatial factors that coordinate the deterministic development of this organism (*4*). Creating analogous maps for mammalian systems has remained out of reach, yet joint cell state and lineage maps in higher organisms would help elucidate many biological processes such as the development of a single fertilized oocyte into a complete organism (*5–8*), the strategies that stem cell populations employ to maintain tissues over the lifespan of an animal (*9–12*), and how disruptions in these processes contribute to disease, including how tumors grow, evolve, and metastasize (*13–20*).

A wealth of approaches over the last century and a half have been developed to track cell lineages (*21*). Just as the mutational clock can be used to assemble phylogenies of species, somatic mutations that accrue in cells can reveal how dynamic biological processes unfold in a single organism. While these approaches enable lineage tracing for human samples (*11*, *22–32*), their temporal resolution is limited by natural mutation frequencies (which may not occur on the timescale of a biological process of interest) and the ease of sampling mutations with high sensitivity across the nuclear and mitochondrial genome. Further, it is currently challenging to assay these mutations with high spatial resolution. By contrast, initial prospective lineage tracing efforts and modern equivalents track static marks in cells–including dyes, fluorescent reporter proteins, or recombined DNA reporter sequences–that, in the context of imaging experiments, can provide a view of the clonal architecture of a tissue (*21*, *33–38*). While these methods can preserve spatial context, they typically necessitate prior knowledge of labeling times or lineages, and can only resolve progenitor outputs at specific time points, thereby limiting their capacity to capture dynamic cellular processes over time.

To address these limitations and thereby resolve cellular phylogenies with high spatial and temporal resolution, there has recently been considerable progress in developing engineered molecular recording technologies that pair cell lineage tracing with rich cell phenotyping (*39–52*). Typically, these approaches employ Cas9-based genome editing to stochastically install heritable lineage tracing marks (LMs) at neutral, pre-determined sites which can then be read out along with cell state using single-cell readouts. Systems with continuous accrual of heritable marks and sufficient diversity can enable reconstruction of vast phylogenetic trees, ideally recording the complete lineage history of all cells in a tissue. While current approaches have predominantly focused on capturing phylogenetic data using sequencing-based readouts in dissociated cells, thereby losing spatial context, there has recently been interest in developing spatially resolved lineage readouts (*34*, *40*, *42*, *53*, *54*). Imaging-based cell profiling methods are uniquely suited for multimodal cell phenotyping, providing true single-cell spatial resolution at tissue scale, while quantifying hundreds to thousands of RNA molecules, protein localization, subcellular morphology, and even epigenetic signatures (*55–70*). However, integrating the depth and timescale of evolving lineage tracing with high-dimensional, spatially resolved cell state mapping *in vivo* remains an important challenge.

Here, we describe a prime editor-based lineage tracer (PEtracer) that integrates recent advances in gene editing, massively multiplexed and multimodal imaging, and lineage tracing methodologies to marry the depth and timescale of evolving lineage tracing with high-dimensional, spatially resolved cell state mapping. We selected prime editing as the backbone of our lineage tracing system based on principled modeling of the properties needed for optimal recording and accurate tree reconstruction (*71*, *72*). Prime editing is a highly flexible genome editing technology that deterministically rewrites stretches of DNA (*73*, *74*), enabling precise DNA modification without reliance on the formation of double stranded DNA breaks (*75–80*).

We use prime editing to install immutable, predefined five nucleotide (5nt) lineage tracing marks (LMs) that can be robustly assayed by both single-cell sequencing (*39*, *53*, *81–84*) (e.g. droplet-based scRNA-seq) and *in situ* hybridization-based imaging (compatible with multiplexed error-robust fluorescence *in situ* hybridization, or MERFISH transcriptomic profiling) (*54*, *56*). Compatibility with both imaging and sequencing single-cell technologies for lineage and cell state measurements allows us to leverage their complementary strengths, enabling rich, unbiased transcriptional profiling integrated with scalable, cost-effective single-cell measurements in tissue sections with subcellular resolution. We optimized this system to stochastically install LMs at predefined edit sites with high efficiency to enable robust lineage tracing over short timescales and tuned tracing kinetics for longer experiments. We performed rigorous *in vitro* benchmarking to demonstrate that PEtracer can reconstruct phylogenies with near-perfect agreement between reconstructed clades and ground-truth static barcodes in a manner that is robust to biological and technical variability.

Further, we used the PEtracer system to study a transplantable model of murine metastatic breast cancer and reconstruct the three-dimensional growth of individually-seeded tumors *in vivo* to dissect how cell-intrinsic and cell-extrinsic factors coordinate cellular behaviors during metastatic outgrowth. Using high-resolution spatial measurements of cell state and lineage information from tens of thousands of cells, we uncover distinct modules linked to malignant cell growth that integrate spatial location, cellular neighborhood, and cell-intrinsic gene expression. These experiments highlight the scalable, robust, and high-resolution spatial view of cell state and lineage that can be achieved with the PEtracer and elucidate how these factors are coordinated to sculpt tumor evolution. More generally, the PEtracer system represents an important step towards creating comprehensive cell fate maps during development and disease in mammalian biology.

### *In silico* modeling of lineage tracing systems

Compatibility with imaging readouts defined key parameters in the design and engineering of an evolving, high-resolution lineage tracing system. In particular, assaying lineage information using fluorescence *in situ* hybridization (FISH)-based imaging requires that each edit outcome used for tree reconstruction (i.e. lineage marks; LMs) can be readily discriminated with a predefined set of probes. Imaging readouts of current Cas9-based lineage tracers would therefore be challenging, as these systems install hundreds of LMs with skewed and stochastic edit distributions (*39*, *80–82*, *85*). We therefore used *in silico* simulations to assess whether a system that uses fewer LMs could generate high-quality phylogenies and how experimentally-relevant parameters might influence tree reconstruction performance with such a system (*86*).

Accordingly, we generated ground-truth phylogenies, simulated the stochastic accumulation of LMs over these trees, and then reconstructed trees using these LMs. This framework allowed us to systematically evaluate how key experimental parameters observed with previous Cas9-based lineage tracing systems affect tree reconstruction accuracy, including: the number of cells being traced, the number of possible editing outcomes for each engineered edit site (number of LMs), the relative installation efficiencies for different LMs (i.e. editing entropy), the number of distinct edit sites used for tree building, the fraction of edit sites where genome editing has installed a LM, and LM detection efficiency. Performance was quantified by calculating both the Robinson-Foulds distance (*87*) and mean number of triplets (*88*) shared between the reconstructed and ground-truth phylogenies (**Methods**).

Our simulations showed that reconstruction accuracy improves with both more balanced LM distributions (i.e. higher normalized entropy; H_norm_) and higher numbers of LMs (**Fig. 1A** and **table S1**) (*89*). However, the incremental benefit of additional LMs diminishes as their number increases (**Fig. 1A**). We decided to use eight well-balanced (H_norm_ >0.9) LMs as this number provides a good compromise between reconstruction accuracy and the feasibility of readout with FISH-based imaging. We found that a system with eight or more balanced LMs can accurately reconstruct trees across a range of final edit fractions–unlike systems with fewer LMs–and performs as well as existing Cas9-based methods across a range of experimental parameters when trees are reconstructed using either UPGMA or neighbor joining (NJ, **fig. S1A-C** and **tables S2,3**) (*90–92*). As expected, low detection efficiency can degrade reconstruction performance, but this effect can be counteracted by increasing the number of edit sites (**fig. S1B**).

**Fig. 1.**
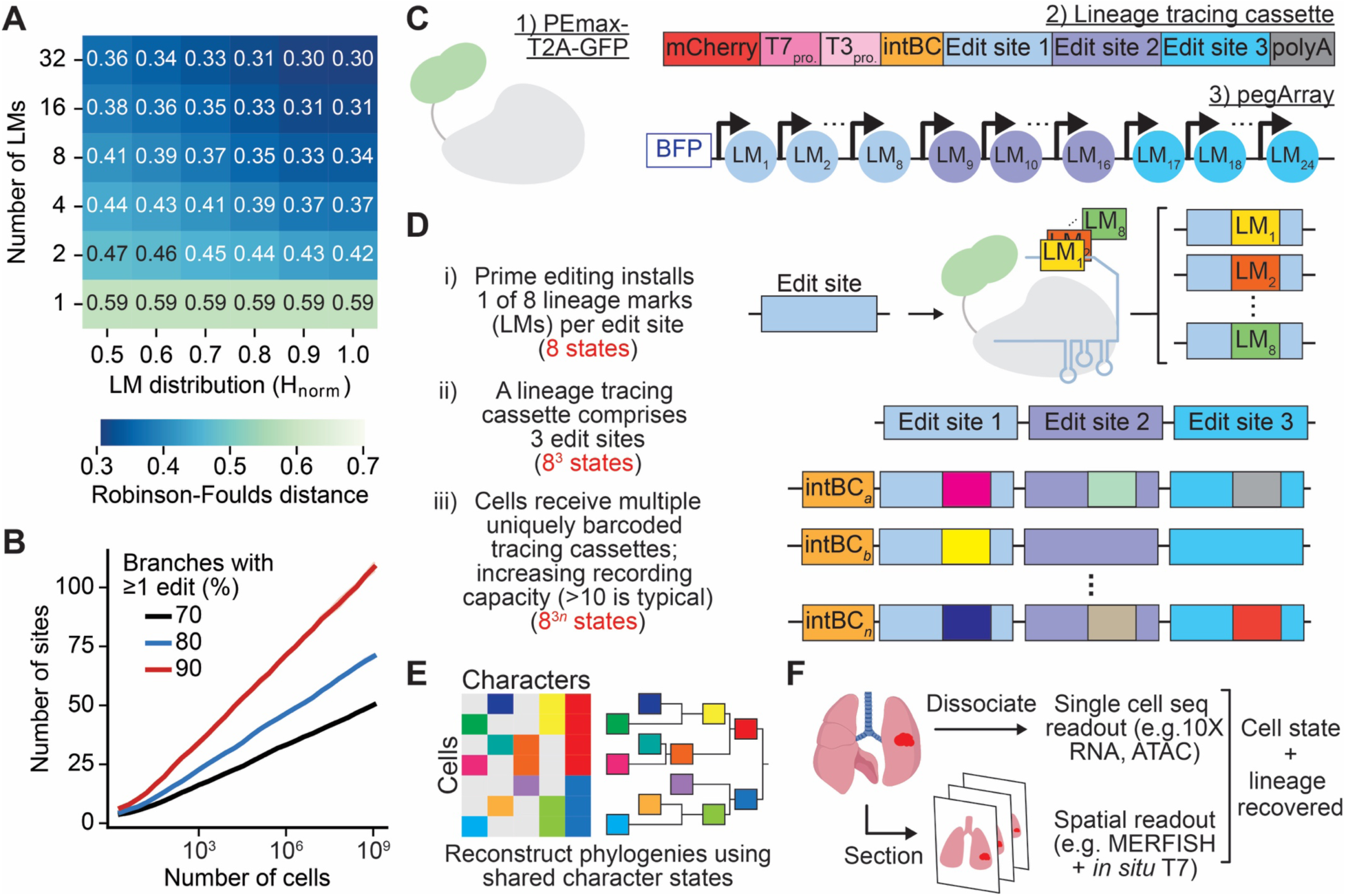
Building the PEtracer system. (**A**) Mean Robinson-Foulds reconstruction error for simulated phylogenies (n = 10) while varying the number of lineage marks (LMs) and normalized entropy (H_norm_) of their relative installation efficiencies. (**B**) Mean fraction of branches in simulated phylogenies (n = 10) marked by any edit for different numbers of cells and edit sites given optimal editing kinetics. (**C**) Components of the PEtracer system: the PEmax-T2A-GFP genome editing agent, lineage recording cassettes in the 3’ UTR of an mCherry gene, where each cassette contains a unique integration barcode (intBC) and three distinct edit sites that can be assayed either by droplet-based single-cell sequencing or by *in situ* transcription via T7 or T3 polymerization, and pegArrays, which encode all 24 LMs used for lineage tracing (eight LMs for each of the three edit sites). (**D**) Schematic of evolving lineage tracing using PEtracer. PEmax installs LMs at pre-determined edit sites, lineage tracing cassettes contain three edit sites, and intBCs enumerate distinct integrated cassettes. (**E**) Character-by-cell matrices detail the continuous accumulation of LMs at edit sites in individual cells throughout a biological process, enabling phylogenetic reconstruction. (**F**) PEtracer cell state and lineage readout compatibility with both dissociative single cell readouts and those that preserve the architecture of the original tissue context. Seq. = sequencing. Lung image in (F) from BioRender.

Evolving lineage tracers enable construction of accurate phylogenies with LMs that are continuously installed throughout a biological process of interest, thereby marking the dynamics of cell fate decisions across time. However, there is a tradeoff between the rate of LM installation and ability to mark later cell divisions due to plateauing as the number of unmodified sites decays over time (**fig. S1D**). Our simulations and theoretical work (*89*) indicate that achieving a final edit saturation of 60-80% provides an optimal balance between rapid LM accumulation and minimal plateauing, thereby maximizing the accuracy of tree reconstructions (**fig. S1B**). Thus, to study biological processes across diverse time scales, the kinetics of a lineage tracing system must be precisely tuned (**fig. S1B,E**). With well-tuned editing kinetics, only 75 edit sites are theoretically required to mark 90% of cell divisions in a phylogeny with 10^6^ cells, which can be extended to 10^9^ cells with 100 edit sites (**Fig. 1B** and **table S4**). Our simulations indicate that large-scale, accurate phylogenetic reconstructions can be performed when eight LMs are installed at edit sites with similar efficiencies (H_norm_ > 0.9). Notably, this design should be well-suited for *in vivo* studies given the robustness to imperfect editing and detection rates (**fig. S1B**).

#### Design of the PEtracer system

We identified prime editing (PE) as a gene editing technology uniquely suited to satisfy the requirements that emerged from our simulation work for high-resolution lineage tracing compatible with both sequencing and imaging readouts. Specifically, compatibility with hybridization-based imaging modalities requires the installation of a predefined set of LMs. Prime editing enables highly precise insertion of diverse, predefined sequences in genomic DNA with minimal undesired byproducts or editing-induced toxicity (*71*, *75–79*). A predefined set of LMs can be encoded by unique pegRNAs targeting the same edit site, and high H_norm_ values can be achieved by selecting pegRNAs that evenly compete during LM installation. Further, PE-installed LMs can be designed to precisely alter the protospacer adjacent motif (PAM) or seed region of the protospacer to prevent re-editing of the targeted sequence, thus installing predefined and immutable marks, unlike Cas9 nucleases or base editors. Finally, the LMs can be designed to facilitate accurate LM discrimination using FISH-based imaging readouts (**fig. S1F**).

Our prime editing-based lineage tracing system (PEtracer) has three primary components: 1) the PEmax prime editor (*72*) whose expression can be monitored using co-expressed GFP, 2) lineage tracing cassettes in the 3’ UTR of an mCherry gene containing a unique DNA integration barcode (intBC) and three distinct edit sites where LMs can be continuously and stochastically installed, and 3) arrays of pegRNAs (pegArrays) encoding 24 LMs (8 LMs per edit site) linked to a BFP gene where each pegRNA encodes a distinct LM (**Fig. 1C-F**). intBCs and LMs can be read out by sequencing of polyadenylated transcripts or imaging of *in situ* transcribed cassettes detected using pre-designed hybridization probes (**Fig. 1F**). Importantly, because each cassette is randomly integrated at a different genomic locus, they are spatially distinct and thus can be resolved by imaging following T7 transcription *in situ* (i.e. the Zombie protocol (*54*)) and distinguished by MERFISH imaging of the intBCs. We designed PEtracer to integrate with MERFISH experiments so we could assay both lineage and endogenous RNA expression in tissue sections. Further, we selected a PE2 editing strategy, as opposed to PE3-type editing which involves opposite strand nicking, to avoid DSB formation and stochastic insertions and deletions (indels) installed at edit sites (*71*).

We engineered the core components of the PEtracer system to meet the benchmarks established with our simulations, which involved three core technical optimizations: identification of edit site sequences and optimized pegRNAs orthogonal to the genomes of various model organisms that enable rapid LM installation while preventing re-editing, selection of a set of LMs that were installed at these loci with similar efficiencies and were maximally discriminable using hybridization-based approaches, and tuning the editing kinetics of LM installation to use the PEtracer system across a broad range of biological processes from the timescale of individual cell divisions to many weeks.

#### PEtracer edit site and LM installation modality optimizations

We first determined the edit site sequences, DNA editing strategy, and LM lengths that allow for highly efficient editing and accurate detection by both sequencing and imaging readouts. We initially screened edit sites and pegRNAs using editing efficiency at an early time point (72 hours post-transfection) as a proxy for rapid editing that would enable labeling of each cell division in lineage tracing experiments over short timescales. Given that prime editing efficiencies can be limited by endogenous cellular mismatch repair (MMR) pathways in a cell type-specific manner (*72*), we aimed to identify DNA edits that would be robust to this constraint. We reasoned that five nucleotide (5nt) marks were short enough to still be installed with high efficiency by PE2 (*93*), should evade efficient MMR detection (*72*), and provide highly diverse sequences that can be robustly discriminated by hybridization probes (**fig. S1F** and **table S5**).

We tested four potential strategies for installing 5nt LMs at edit sites: 1) inserting a 5nt mark between the protospacer adjacent motif (PAM) and the adjacent genomic target sequence recognized by the protospacer of our pegRNAs, 2) deleting the PAM and 3nt of the genomic target seed region (positions 18-20 of the protospacer where the PAM NGG are positions 21-23) and replacing these deleted bases with a 5nt mark, 3) performing the same edit as variant 1 but recoding the 3nt seed and PAM, and 4) replacing the 3nt seed sequence with a 5nt mark (**Fig. 2A**, right). Each of these editing strategies prevents re-editing, which ensures immutable, heritable marks and boosts PE efficiency (*71*). We tested the installation of two representative 5nt sequences using each of these four editing strategies across six genomic loci (*AAVS1*, *EMX1*, *FANCF*, HEK293T site 3 [HEK3], *RNF2*, and *RUNX1*) in HEK293T cells using pegRNAs with fixed primer binding site (PBS) lengths and a range of reverse transcription template (RTT) lengths previously used for these loci (*94*). We selected three loci with consistent and efficient editing: edit type 1 at the *RNF2* and HEK3 loci and edit type 2 at the *EMX1* locus (**Fig. 2A**). Edits at the *RUNX1* and *AAVS1* loci also performed well and may serve as useful starting points for future expansions to the PEtracer technology. We performed further RTT optimization using epegRNAs, which contain a structured 3’ end (*94*), to increase PE editing performance (*RNF2* = 56%, 1.5-fold improvement; HEK3 = 60%, 3.6-fold improvement; *EMX1* = 66%, 3.6-fold improvement; **fig. S2A**). The improved editing efficiencies observed with these optimized epegRNAs enable efficient labeling of these edit sites during a single cell division even in rapidly dividing cells.

**Fig. 2.**
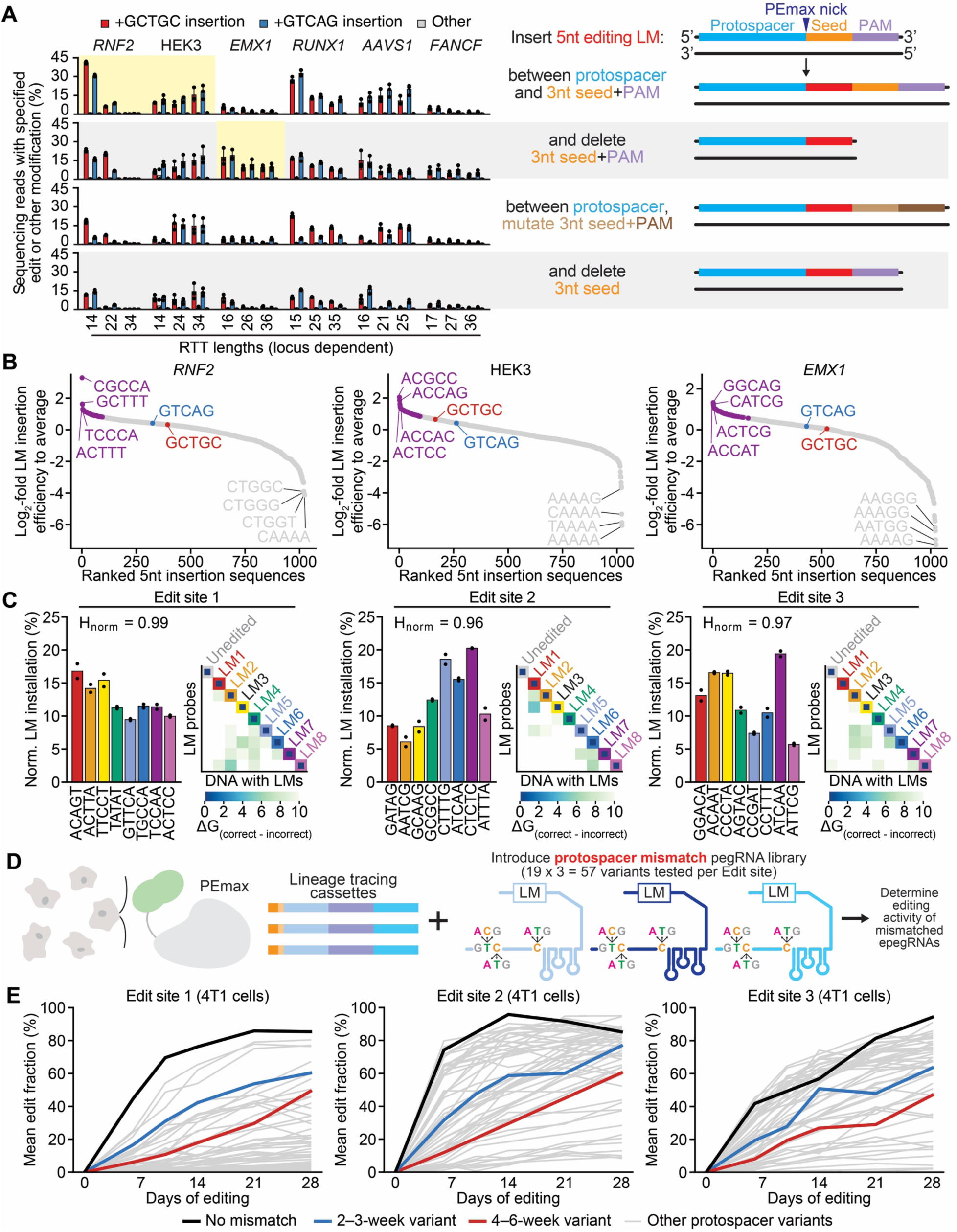
Engineering of the PEtracer system. (**A**) Optimization of lineage mark (LM) insertion strategies (diagrams on right) across six endogenous genomic loci (*RNF2*, HEK3, *EMX1*, *RUNX1*, *AAVS1*, *FANCF*) using two representative five nucleotide (5nt) LMs in HEK293T cells. Editing rates reflect the percentage of sequencing reads that contain the intended edit out of all aligned reads. Mean of three biological replicates ± standard deviation depicted. Sites and strategies used for subsequent experiments highlighted in yellow. Reverse transcription template (RTT) lengths are specific to each edit site. (**B**) Comprehensive LM screen comparing insertion efficiency of all possible 1,024 5nt insertions at three test loci in HEK293T cells. Log_2_-fold insertion efficiencies were calculated relative to input plasmid library and averaged across three biological replicates. Representative 5nt LMs used in (A) shown in red and blue, and 96 best-performing LMs highlighted in purple. (**C**) Bar plot of the normalized (Norm.) LM installation efficiencies for the eight LMs in the final 24-mer pegArray (without protospacer mismatches) at each of the three finalized edit sites in B16-F10 cells (on left for each edit site) averaged across two biological replicates; H_norm_ values provided for each edit site. Predicted probe hybridization specificity (Gibbs free energy = ΔG) for discriminating each of the eight LMs from one another (on right for each site). (**D**) Schematic of epegRNA protospacer mismatch screening for tuning lineage tracing kinetics. (**E**) Editing kinetics for protospacer mismatch variants at edit sites 1, 2, and 3 in 4T1 cells. Grey lines show all protospacer variants tested; no mismatch protospacers shown in black; protospacer mismatches selected for 2–3-week timescales shown in blue; 4–6-week protospacer mismatches in red.

Our initial optimizations were performed at human genomic loci; therefore, cognate epegRNAs would modify endogenous sequences in addition to our edit sites in human cells. We therefore evaluated two orthogonalization methods–either complementing (Ortho v1) or reverse complementing (Ortho v2) the protospacer seed sequence. We generated synthetic lineage tracing cassettes that harbored three target sequences in tandem (see **Fig. 1C**) comprising the endogenous human sequences or the Ortho v1 or Ortho v2 variants. While epegRNAs targeting the genomic sequences showed high LM insertion efficiencies at both the lineage tracing cassettes and at the endogenous target loci, we observed equivalent editing efficiencies at our orthogonal edit sites with virtually no modification of endogenous sequences with orthogonal epegRNAs (**figs. S2B,C** and **Methods**). We moved forward with Ortho v1 edit sites and cognate epegRNAs. Importantly, *in silico* analysis revealed no potential CRISPR/Cas9-dependent off-target loci with three or fewer mismatches in humans or common model organisms as likely to support prime editing (**fig. S2D** and **Methods**) (*95*).

As done previously (*39*), we constructed lineage tracing cassettes by inserting our edit site sequences in the 3’ UTR of an mCherry reporter so that we could enrich for cells with high numbers of tracing cassette integrations by FACS sorting (**Fig. 1C**). The orthogonalized variants of the *RNF2*, HEK3, and *EMX1* loci became edit sites 1, 2, and 3, respectively, with all editing occurring on the same DNA strand to minimize the generation of tandem duplications or indels (*72*, *96*). Lineage tracing cassettes are expressed at a high level for efficient capture by single-cell droplet-based sequencing technologies. To facilitate detection by imaging, we introduced T7 and T3 promoters in the lineage tracing cassette to enable *in situ* transcription and thus amplification of these integrated cassettes (*54*). Each cassette is marked by an integration barcode (intBC) containing two barcode sequences, a 30nt barcode for sequencing readout and a 183nt barcode designed to have optimal GC content, Tm, and sequence orthogonality for hybridization-based imaging readout. We generated a library of 2,171 intBC-tagged lineage tracing cassettes to enable polyclonal cell line engineering where individual clones can be distinguished based on the set of intBCs present (**table S6**).

#### Selection of balanced LM sequences compatible with imaging-based readout

With tracing cassettes and preliminary epegRNAs designed, we next sought to screen through the set of 1,024 possible 5nt LMs to identify sequences that were both efficiently installed and maximally discriminable by hybridization-based readouts. While our initial optimizations were performed using two representative 5nt insertions, we reasoned that different sequences were likely to be installed with higher efficiencies, thus enabling tracing on shorter timescales. To identify high-efficiency LMs, we transfected HEK293T cells with epegRNA pools representing all possible 1,024 LM sequences for each target locus. We observed unique LM installation preferences for each site, with the exception of a common bias against polyA sequences that act as terminators for the U6 promoter by creating a polyT stretch in the epegRNA (**Fig. 2B** and **table S7**). We identified many LMs that were installed with markedly higher efficiency than the test sequences, and further validated the installation efficiencies for the top scoring LMs for each site (*EMX1* = 96, HEK3 = 94, *RNF2* = 85) in an arrayed manner (**fig. S3A**).

From these sets of efficiently-installed LMs, we sought to identify those that could be robustly distinguished by imaging readouts at orthogonal edit sites. We selected LMs with installation efficiencies within 10% of one another and computed their predicted discriminability from one another and the unedited state by hybridization probes for each site (**figs. S3A,B** and **table S8**). Unsurprisingly, a number of these LMs are predicted to be challenging to distinguish due to sequence similarity (*e.g.* GCTTT and ACTTT at Edit Site 1). After excluding LMs with high pairwise cross-hybridization, we computationally nominated a set of 20 potential 5nt LM sequences (**fig. S3A,B**). Retesting at each orthogonalized edit site confirmed consistent and balanced insertion activity (**fig. S3C**). The best-performing 18 of these LMs were then used for subsequent engineering efforts to ensure similar installation efficiencies among competing epegRNAs for each edit site.

Finally, to identify sets of eight epegRNAs per edit site where each LM is installed with a similar efficiency when all epegRNAs are expressed in the same cell, we created arrays comprising eight epegRNAs each (8-mer arrays, **table S9)**. Initial 8-mer epegRNA arrays showed reasonably balanced performance, with H_norm_ ranging from 0.90 to 0.98 (**fig. S4A**); however, through an additional round of rational engineering, we achieved more balanced installation efficiencies (H_norm_ for edit site 1 = 0.98, 2 = 0.99, 3 = 0.96**; fig. S4B**). We noted that the first two positions in the 8-mer arrays typically showed lower editing activity, so we strategically ordered the epegRNAs to preserve LM balance when we constructed a 24-mer array consisting of an 8-mer for each edit site, resulting in balanced and efficient installation for each lineage mark at its corresponding edit site (**Fig. 2C**). Moreover, the sets of eight LMs for each edit site are predicted to be highly discriminable from one another and the unedited state based on calculated Gibbs free energy differences, suggesting that we can generate hybridization probes that specifically detect each LM (**Fig. 2C** and **table S8**).

#### Kinetic tuning of epegRNAs to use PEtracer across diverse timescales

To identify epegRNAs that would enable continuous editing over timescales ranging from several days to months, we tested a library of mismatches in the protospacers of epegRNAs targeting each edit site. We exhaustively altered all protospacer positions except the first initiating G for a total of 171 epegRNA variants (3 bases x 19 positions = 57 variants per protospacer x 3 protospacers) using a single representative LM for each edit site (**Fig. 2D**). We generated a lentiviral library using a CROPseq-style design where epegRNA variants were engineered into the 3’ LTR of a lentiviral genome and the UTR of a polyadenylated mRNA to enable capture in single-cell sequencing experiments (*97*). With this design, we can determine both epegRNA protospacer identity and the fraction of sites edited for each cell by scRNA-seq. We collected time points across a 28-day experiment in two MMR-competent mouse cancer cell lines which have been widely used to model melanoma and breast cancer metastasis (B16-F10 and 4T1, respectively) (*98–100*) to assay editing performance of all 171 protospacer variant epegRNAs in pooled experiments. These data create a suite of epegRNA protospacer mismatches that slow LM installation for use in lineage tracing experiments across diverse timescales (**fig. S1E**).

We identified numerous protospacer mismatches with slower rates of LM installation than unmodified protospacer sequences (**Fig. 2E** and **fig. S4C**; black lines). To estimate the rate at which edit sites are modified (in LM edits per day), we fit curves modeling the exponential decay of available, unmodified edit sites over time (**table S10**). When accounting for a small fraction of cells that remain unedited, we observe high concordance between these modeling results and real data, suggesting a uniform editing rate throughout the experiment. Across both cell lines, consistent patterns of protospacer mismatches were observed to decrease the editing rate, with PAM-proximal mismatches having the largest decreases in editing rate, as expected (**fig. S4D**). Notably, the relative decrease in editing efficiency for each variant was consistent between both cell lines; however, the median edits per day was 1.74-fold faster in B16-F10 cells compared to 4T1, highlighting cell type-specific differences in PE performance that may alter absolute editing rates (**fig. S4E,F**). We selected protospacer variants which tuned the desired editing rate to match the timescales for subsequent *in vitro* and *in vivo* experiments over 2-3 weeks (0.05 - 0.1 edits/[edit site * day]) and 4-6 weeks (0.02 - 0.04 edits/[edit site * day]), respectively (**Fig. 2E**, blue and red lines, respectively). For example, with 60 edit sites and a rate of 0.02 edits/[edit site * day], on average one edit would be introduced per day over 4-6 weeks. Notably, other protospacer variants screened here would enable tracing over even longer timescales.

### *In vitro* validation of PEtracer readout using fully-edited cells

To measure PEtracer integration barcode (intBC) and LM detection efficiency and decoding accuracy using both sequencing and imaging readouts, we generated cells with defined linkages between integration barcodes and lineage marks (**Fig. 3A**). We selected clones of 4T1 cells from a polyclonal population with high numbers of integrated tracing cassettes edited to saturation. These clones have distinct sets of integrations, typically ∼10 copies and thus ∼30 edit sites per clone. We pooled four clones to cover all 24 LMs as well as the unedited state for all three edit sites (**Fig. 3B,C**). Using single-cell sequencing we created a whitelist linking intBCs with consensus LMs for each clone (**table S11)**, which we then used to evaluate the detection efficiency and accuracy of lineage readout across 6,883 sequenced single cells. We observed a 98.5% true positive rate for intBC detection with only a 1.3% false positive rate and we correctly called LMs for all detected integrations with an accuracy of 99.9% (**Fig. 3D,E** and **fig. S5A**).

**Fig. 3.**
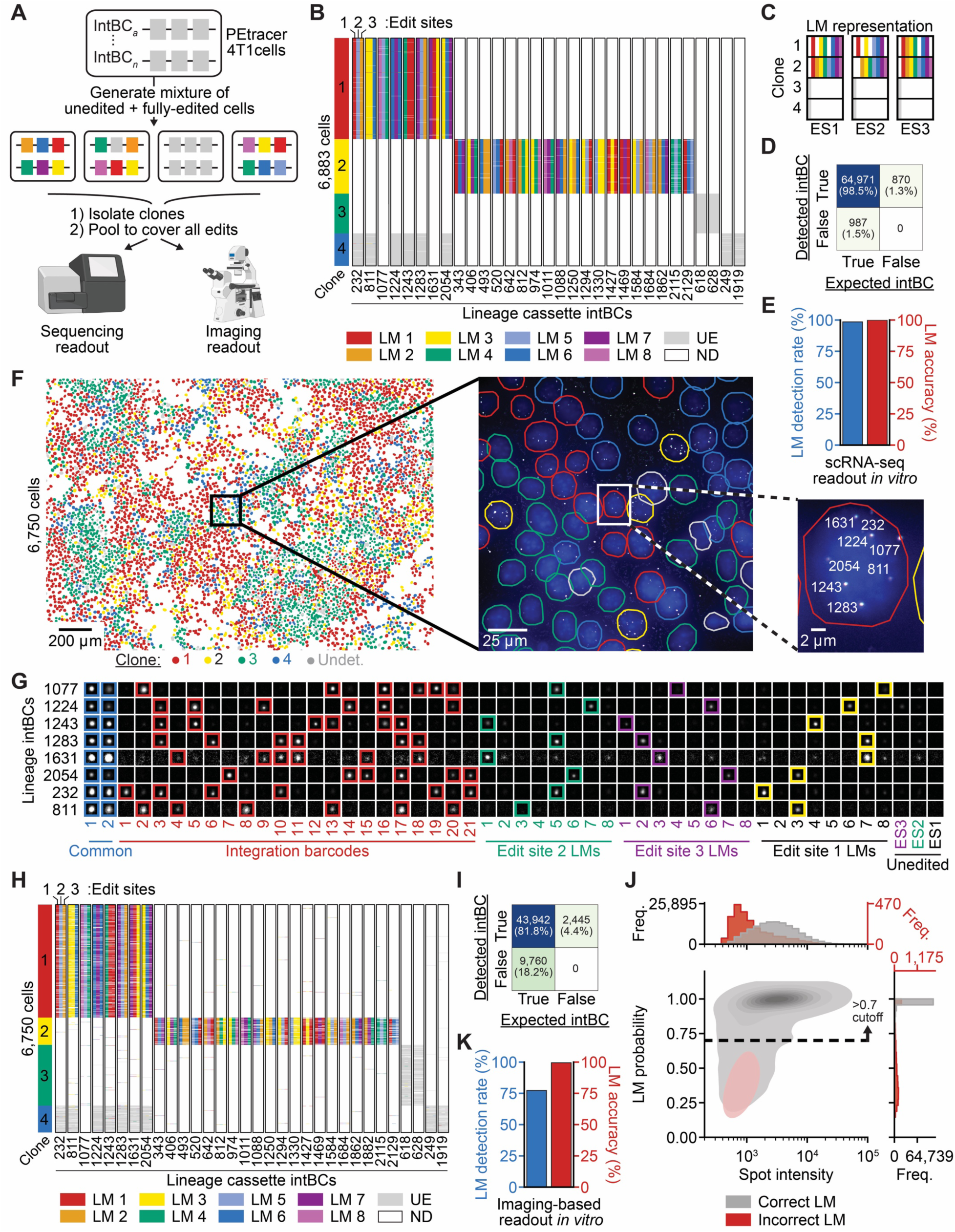
Evaluation of PEtracer intBC and LM detection efficiency and decoding accuracy. (**A**) Experimental schematic for generating fully-edited PEtracer cells with defined linkages between integration barcodes (intBCs) and lineage marks (LMs) to enable ground-truth evaluation of intBC and LM detection efficiency and decoding accuracy. Representative sorting gates shown in fig. S14. Sequencer and microscope images from BioRender. (**B**) Droplet-based single-cell RNA sequencing (scRNA-seq) of 4 clones (6,883 cells). Color blocks on the left indicate clone identity, where clone 1 = red, clone 2 = yellow, clone 3 = green, and clone 4 = blue. Lineage tracing cassettes consisting of three edit sites are annotated with their respective intBCs at the bottom and show distinct LM combinations across the different clones. Unedited state (UE) is shown in grey; when an intBC was not detected (ND) in a cell, it is shown as white. (**C**) Summarized LM representation across the four combined clones. All 24 LMs are observed in the fully-edited clones 1 and 2; the unedited state is represented in clones 3 and 4. ES = edit site. (**D**) Confusion matrix depicting true positive, false negative, and false positive rates for detected and expected intBCs by droplet-based scRNA-seq. (**E**) LM detection rate (blue, left) and accuracy (red, right) for scRNA-seq data based on expected intBC-LM pairing. (**F**) Wide field view (left) of fully-edited and unedited clones plated for imaging-based lineage readout experiments pseudo-colored to indicate clonal identity. Color scheme matches (B), with the addition of undetermined cells (Undet.) that could not be confidently assigned to a clone shown in grey. Higher magnification of a field of view (middle) and a representative single cell (right) with DAPI nuclear staining (blue) and the two common bits (C1 = green; C2 = magenta). Integration amplicons appear as colocalized common bit puncta (white) and decoded intBCs are numbered to match our codebook (see table S6). (**G**) Example readouts for intBC and LM detection and decoding following *in situ* T7 transcription of integrated lineage tracing cassettes. Seventeen imaging rounds in three colors (50 total bits, columns) for distinct integration amplicons (rows) include two common bits to identify candidate spots followed by 21 integration barcode bits (Hamming distance 4, Hamming weight 6 code), 24 LM bits, and three bits for the unedited state. (**H**) Equivalent panel to (B) for imaging-based readout of 6,750 fully-edited cells using *in situ* T7 transcription of lineage tracing cassettes. (**I**) Confusion matrix as in (D) for imaging-based readout. (**J**) Validation and performance of the logistic regression classifier for LM assignment from imaging-based PEtracer data. Freq. = Frequency. Assignment probability cutoff of p >0.7 for decoded LMs is noted as a dashed line. (**K**) LM detection rate and accuracy after filtering and decoding with the LM classifier in (J).

To evaluate intBC and LM detection rates and accuracy by hybridization-based imaging readout, we first used single-cell sequencing to create a mapping of 30nt sequencing barcodes to 183nt imaging barcodes for each integration (**Methods**, **fig. S5B,** and **table S11**). With this link established, we plated fully-edited cells at high density onto coverslips and used T7 RNA polymerase to generate localized RNA amplicons *in situ* from genomically integrated lineage tracing cassettes using an in-gel transcription protocol adapted from the Zombie protocol (*54*). Building on previous insights that tissue clearing procedures enhance the detection efficiency and detection limit for both native and T7 polymerase *in situ*-produced RNA species (*57*, *101*), we extended standard MERFISH sample preparation protocols to include extensive digestion, denaturation, and tissue clearing of polyacrylamide hydrogel embedded samples (**Methods** and **tables S12-16**). Indeed, this substantial clearing increased the total fraction of decodable intBCs generated by *in situ* T7 amplification by 19.8% compared to the standard Zombie protocol, primarily by improving decoding performance for dimmer amplicon spots (**fig. S5C-E**). Across 5,614 plated cells (**Fig. 3F**), we conducted 17 rounds of 3-color MERFISH imaging to read out lineage tracing information by identifying candidate amplicons (with two colocalized bits for common sequences within lineage tracing cassettes), decoding intBCs (using a combinatorial detection scheme with a modified Hamming distance 4, Hamming weight 6 binary 21-bit code), and detecting LMs and the unedited state for the three edit sites (using 27 separate bits) (**Fig. 3G,H**). With this improved protocol, the intBC detection rate was 81.8% without an appreciable increase in the false positive rate relative to the standard Zombie protocol (**Fig. 3I** and **fig. S5D**).

We next optimized our decoding of LMs across detected spots. First, using the maximum spot intensity across hybridization rounds to assign LMs, we achieved 97.7% accuracy (**fig. S5F**). However, this approach does not account for differences in probe affinity and cross hybridization that might affect LM spot brightness, thus we trained a logistic regression classifier to identify the most likely LM for each site given a vector of spot brightness across the rounds of imaging (**fig. S5G**). With 5-fold cross validation, we achieved a decoding accuracy of 98.2%, which was further improved by applying a probability threshold (p>0.7) for LM assignment (**Fig. 3J** and **fig. S5F**). After excluding low-confidence LMs (which tended to be dimmer on average) our final LM detection rate was 78.6% while the accuracy increased to 99.4%–with each LM being individually decoded with >98.3% accuracy (**Fig. 3K** and **fig. S5H**). Based on our simulations, these LM detection and accuracy rates enable accurate tree reconstruction. Having confirmed efficient and accurate intBC and LM measurement in fully-edited cells, we next set out to benchmark lineage tree reconstruction performance using the PEtracer system.

### Benchmarking of *in vitro* tree reconstruction

Next, we rigorously evaluated our ability to reconstruct phylogenies from cells undergoing evolving lineage tracing using droplet-based single-cell sequencing. We used two successive rounds of static barcoding to introduce independent, ground-truth marks of shared ancestry at sequential time points to empirically assess the accuracy of phylogenies reconstructed using PEtracer (**Fig. 4A**). In 4T1 cells with ∼10-15 copies of the tracing cassettes and a pegArray tuned to edit over a 2–3-week timescale, we initiated evolving lineage tracing using a lentiviral construct expressing the PEmax editor. Lentiviral barcode libraries encoded in the 3’ UTR of either a puromycin or blasticidin resistance cassette were introduced at days five and seven respectively after tracing was initiated. On day 16, we harvested cells and performed single-cell RNA sequencing, capturing evolving LM data, transcriptional profiles, and puromycin- and blasticidin-linked static barcodes for each cell.

**Fig. 4.**
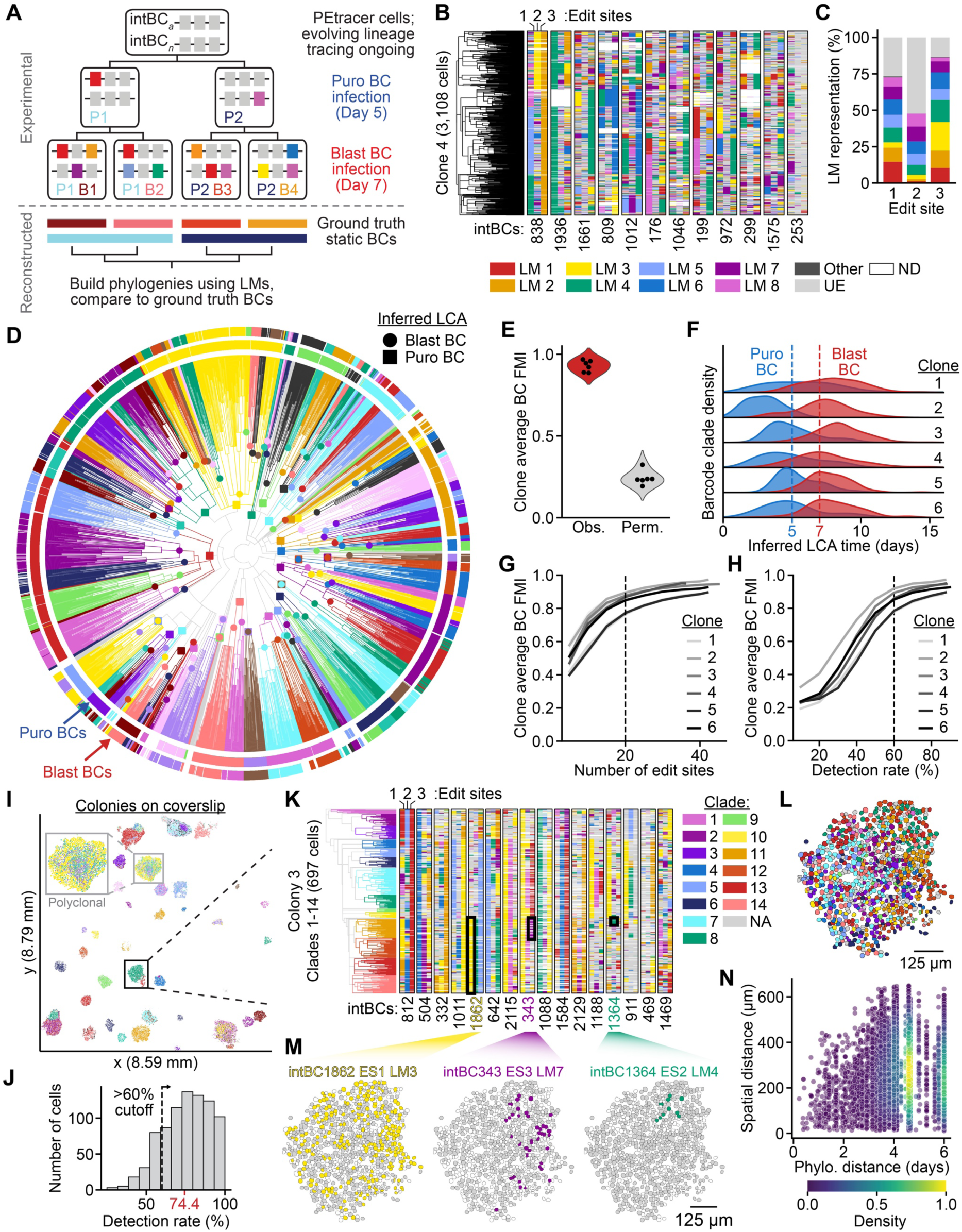
Reconstruction of high-resolution, accurate phylogenies with PEtracer. (**A**) Schematic of *in vitro* benchmarking experiment where phylogenies reconstructed based on the continuous accrual of lineage marks (LMs) can be evaluated for their ability to correctly group cells that share static barcodes introduced at two timepoints. Top shows the experimental design, where cells engineered with PEtracer components begin evolving lineage tracing on day zero, followed by two rounds of lentiviral infection with static barcode (BC) libraries associated with antibiotic selection markers puromycin (Puro; blue lettering) and blasticidin (Blast; red lettering). Bottom depicts how ground-truth static BC groups can be compared to reconstructed phylogenies. (**B**) Character matrix captured by single-cell RNA-seq (scRNA-seq) readout paired with phylogenetic tree reconstructed from LMs using the neighbor joining algorithm for the 3,108 cells in clone 4. Each integration barcode (intBC) is listed below its set of three corresponding edit sites that are colored by LM identity; UE = unedited, ND = not detected. (**C**) Averaged edit site modification frequencies for six clones assayed in this experiment colored by LM. (**D**) Phylogeny from (B) depicted as a circle with rings colored to show puromycin (inner ring) and blasticidin (outer ring) static BC groups for each cell (white indicates missing data). Phylogeny branches colored to match BC assignments. The inferred lowest common ancestor (LCA) is marked and colored to match each BC group (Puro = square; Blast = circle). (**E**) Averaged Fowlkes-Mallows Index (FMI) scores for puromycin and blasticidin BC groups measuring the agreement between LCA clades and observed (Obs.) static barcode labels versus randomly permuted (Perm.) barcode labels across the 6 phylogenies captured in this experiment. (**F**) Inferred LCA timing for puromycin and blasticidin static BC groups across 6 phylogenies. Dashed Day 5 and Day 7 lines indicate the true timing of when these BCs were introduced into cells. Averaged puromycin and blasticidin barcode FMI scores when down-sampling the number of edit sites (**G**) or the detection rate (**H**) used for tree reconstruction with six experimental phylogenies assayed using scRNA-seq. (**I**) Imaged coverslip of 4T1 cells growing as intermixed clones with cells pseudo-colored to indicate clone identity. x and y dimensions provided in millimeters (mm). Grey inset box highlights a polyclonal colony comprising clones 2, 14, 15, and 29. Black inset box shows clone 3 of 64, which we analyze in (J-N). (**J**) Histogram of intBC detection rate for cells in clone 3. The average detection rate is shown in red; a 60% cutoff for cells included in phylogenetic reconstructions is noted with a dashed line. (**K**) Character matrix captured by imaging readout of the 697 cells with robust intBC and LM detection in clone 3. LM annotations identical to (B) are shown across the intBCs enumerated on the x-axis. Clades 1 through 14 are colored on the phylogeny to the left. NA = not assigned a clade. (**L**) Nuclear masks for cells in clone 3 colored by their phylogenetic clade assignment. (**M**) Spatial positions of cells in clone 3 marked with specified edits that define spatial neighborhoods of cells. (**N**) Pairwise phylogenetic (Phylo.) versus spatial distance for cells in clone 3.

We then reconstructed phylogenies for six clonal groups of cells using only the evolving LM information (**Fig. 4B** and **fig. S6**). These phylogenies show efficient LM labeling of each edit site (edit site 1 = 73.4%, edit site 2 = 47.7%, edit site 3 = 86.5%), with representation from all LMs (**Fig. 4C**). Notably, unintended edits (“other” in **Fig. 4B,C**) occur at extremely low rates (edit site 1 = 0.54%, edit site 2 = 0.07%, edit site 3 = 0.36%) consistent with the high editing fidelity observed with PE2, ensuring that these phylogenies can be readily assayed using both sequencing and imaging readouts. To assess the quality of reconstructed trees, we quantified the agreement between groups of cells with a shared static barcode and phylogenetic clades defined by the inferred point in the tree when each barcoding event likely occurred (the lowest common ancestor; LCA). Strikingly, we reproducibly observed near-perfect agreement between LCA clades and static barcoding groups (**Fig. 4D**, **fig. S6**, and **fig. S7A)**. We quantified this agreement using the Fowlkes-Mallows Index (FMI) (*102*) across six independent phylogenies (FMI scores ranging from 0.85 to 0.97; maximum value of 1) (**Fig. 4E**, **fig. S7A**, and **table S17**). Notably, imperfect calling of barcode groups was partially responsible for FMI scores <1 (**fig. S6**). Both puromycin and blasticidin static barcode groups have consistently high FMI scores (mean = 0.92, minimum = 0.85, maximum = 1.00) when trees are reconstructed with either NJ or UPGMA algorithms (**fig. S7B**). The high FMI scores for both puromycin and blasticidin groups demonstrate high accuracy at multiple depths within the reconstructed phylogenies. Given the consistent rate of editing demonstrated by our kinetics experiments (**Fig. 2E**), we could estimate the lifespan of each ancestral point in the tree and use these branch lengths to infer when static barcoding events occurred in the experiment. While these branch length estimates tend to be noisy, encouragingly, the inferred mean time for puromycin (5 days) and blasticidin LCAs (7 days) aligned with the timeline of the experiment for most clones (**Fig. 4D,F, fig. S7A**, and **table S17**).

As a tree-independent evaluation of tracing accuracy, we calculated the Hamming distance (0 = all LMs the same at all edit sites, 2 = all LMs different) between pairs of cells that either share or do not share a static barcode group. When evaluating these phylogenies (*e.g.* Clone 4; see **Fig. 4D**), pairs of cells with matched puromycin barcodes shared significantly more LMs than unmatched pairs (KS = 0.80), with even greater LM overlap for blasticidin barcodes (KS = 0.89), as cells received these barcodes later in the experiment and therefore should group together more closely related cells (**fig. S7C**). Phylogenetic distances (days since LCA) using our estimated branch lengths are strongly correlated with Hamming distances, but more clearly distinguish matched versus unmatched puromycin (KS = 0.96) and blasticidin (KS = 0.98) static barcode pairs (**fig. S7C**). We observe the same trend across all sampled clones where phylogenetic distance outperforms Hamming distance, particularly for puromycin barcode pairs which tend to be less well separated (**fig. S7D**). This improvement arises from using a cumulative metric over all observed recording sites for tree reconstruction with distance-based algorithms (such as NJ or UPGMA), which mitigates the effect of homoplasy (the same LM independently occurring at the same site in multiple cells).

We performed down-sampling analyses to determine reconstruction accuracy as experimentally-relevant conditions vary. Encouragingly, across the six experimentally-generated phylogenies benchmarked by static barcoding, we found that reconstruction accuracy assessed by FMI score remained high with >20 edit sites and with a detection rate of >60%–cutoffs that agree with our simulation work and which we use for subsequent reconstructions (**Fig. 4G,H** and **tables S18,19**). These experiments and down-sampling analyses demonstrate that we can build accurate phylogenies using the PEtracer system across a range of experimental conditions.

#### Imaging-based spatial lineage analysis of motile cells

To evaluate the quality and scalability of phylogenetic reconstructions generated using high-resolution, imaging-based PEtracer data, we profiled the growth dynamics of highly-motile 4T1 cells *in vitro* (*103*). We sparsely seeded cells with active lineage tracing components onto glass coverslips and allowed these cells to grow into colonies over six days (**Fig. 4I** and **fig. S8A**). For one of the imaged coverslips, we captured lineage information for 18,675 cells across 64 evolving clones as large as 1,167 cells (**Fig. 4I**, **fig. S8A**, and **table S20**). These clones had an average of 40±10 edit sites with a mean edit fraction of 76.4±27.2% and an intBC detection rate of 75.8±2.5%, values that our simulations and benchmarked barcoding data suggest would enable reconstruction of high-accuracy trees (**Fig. 4J** and **fig. S8B-D**). Colonies can be seeded by individual cells or by multiple cells that grow into one another. Indeed, in these colonies, we observed considerable intermixing between seeding clones within polyclonal colonies, highlighting substantial cell movement during colony expansion (**Fig. 4I**, inset). Similarly, we observed marked intermixing between distinct phylogenetic clades within clones (**Fig. 4K,L**). When we look at the growth dynamics within a clone, we observe that LMs that occur early are widely distributed, likely due to movement during the initial stages of colony growth, whereas LMs that occur later are restricted to progressively smaller spatial regions, reflecting the limited time for dispersion since these cells shared a more recent common ancestor and confirming the accuracy of these phylogenies (**Fig. 4M**). Indeed, phylogenetic and spatial distances are correlated, with mean spatial distances between pairs of cells that share a common ancestor within the last three days being substantially lower than randomly permuted pairs (**Fig. 4N**, and **fig. S8E**). This trend is consistent irrespective of detection rate, indicating that the thresholds determined by our simulation efforts enable accurate phylogenetic reconstructions (**fig. S8E**). These conclusions are further supported by the correlation between phylogenetic and LM distances (**fig. S8F**), and similar patterns are observed in other sampled clones, where close relatives are near one another, but cell movement quickly degrades the coupling of phylogenetic and spatial distances (**Fig. 4N** and **fig. S8G**). Direct observation of these colonies over time supports the intermixing of clones and rapid cell movement during their outgrowth (**Supp. video 1**). These results indicate that PEtracer imaging data can be used to reconstruct large, highly-accurate phylogenies with precise, true single-cell spatial resolution.

### Spatially resolved transcriptomic profiling of metastatic tumors *in vivo*

We next applied PEtracer *in vivo* to jointly profile cellular state, microenvironment, and phylogeny using the syngeneic 4T1 breast carcinoma cell line to model cancer cell extravasation, metastatic colonization, and growth in immunocompetent mice (*98*, *99*, *104*). We engineered 4T1 cells as in the colony growth experiment, but with a pegArray tuned to edit over the 4-6 weeks of *in vivo* tumor growth, and initiated clonal lung metastases by tail vein injection (**Fig. 5A**).

**Fig. 5.**
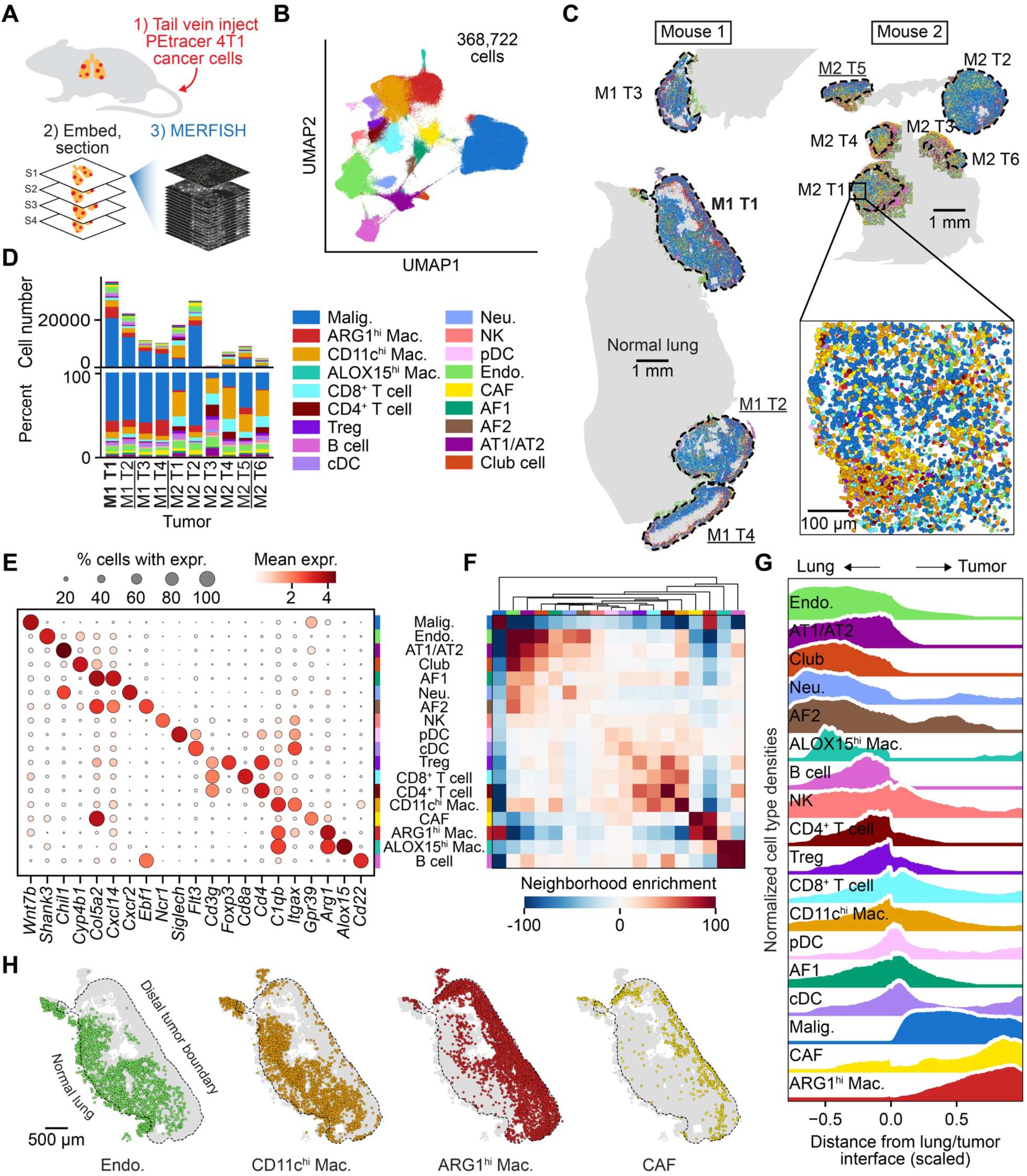
Spatial transcriptomics of metastatic 4T1 lung tumors. (**A**) Experimental schematic for generating MERFISH data from lung tumors following tail vein injection of 4T1 cells engineered with PEtracer components into immunocompetent mice. PEtracer 4T1 cells undergo evolving lineage tracing throughout tumor growth; tumor-bearing lungs are embedded for sectioning and imaged. (**B**) UMAP embedding of 368,722 cells sampled from 10 tumors in two animals, where each cell is colored by its assignment to the 18 distinct cell types resolved by this library. Cell type color annotations are listed below the UMAP. (**C**) Spatial organization of cell types in 4T1 lesions in the lungs of two mice, with a zoom-in showing nuclear segmentation masks. Cell types are colored using the annotations in (B). Mouse 1 (M1; left) has four distinct tumors (T1-T4); Mouse 2 (M2; right) has six (T1-T6). M1 T1 is bolded to indicate that this tumor is analyzed more deeply in main text Fig. 6 and fig. S12. M1 T2, M1 T4, and M2 T5 are underlined as they are analyzed further in fig. S11. Adjacent normal lung is shown in grey; tissue-level scale bars = 1 mm; inset scale bar = 100 µm. (**D**) Percentage and cell number counts for each cell type grouped by tumor, colors consistent with those in (B). (**E**) Marker gene expression (expr.) across annotated cell types. Dot color indicates the mean gene expression for a marker gene within a cell type, while dot size indicates the percentage of cells within a cell type that express that gene. (**F**) Cellular neighborhood enrichment (k = 6 closest neighbors) for cells in sampled tissues across M1 and M2. (**G**) Normalized cell type densities as a function of their scaled distance to the lung/tumor interface (boundary between adjacent lung and tumor; −1 corresponds to cells within the lung furthest from the lung/tumor interface and +1 corresponds to cells within the tumor furthest from the lung/tumor interface). Cell type colors consistent with legend in (B). (F, G) depict data from 9 tumors (M2 T3 excluded from analysis due to low cell number). (**H**) Spatial distribution of Endothelial cells, CD11c^hi^ macs., ARG1^hi^ macs., and CAFs across M1 T1 section 2. Cell type abbreviations are as follows: Malig. = malignant, Mac. = macrophage, Treg = regulatory T cell, cDC = conventional Dendritic Cell, Neu. = Neutrophil, NK = Natural Killer cell, pDC = plasmacytoid Dendritic Cell, Endo. = Endothelial cell, CAF = Cancer Associated Fibroblast, AF1/AF2 = Alveolar Fibroblast Type 1/2, AT1/AT2 = alveolar epithelial type 1/2.

After 4-5 weeks of tumor growth, we collected 20 μm sections from the lungs of tumor-bearing mice. We selected 124 genes based on scRNA-seq data (*105–107*) (**Methods** and **table S21**) to enable annotation of major cell types including cancer, immune, and stromal cells. With this panel, we first assessed the cell type composition in 4T1 lung lesions with MERFISH imaging of endogenous cellular RNAs *in situ* (**Fig. 5A**). We assayed RNA expression using a custom 16-bit MERFISH library in five tissue sections encompassing 10 tumors from two mice–in total, we profiled 368,722 total cells with an average of 178 transcripts detected per cell (**Fig. 5B-D, fig. S9A,** and **table S22**). Extensive experimental optimizations to prevent RNA degradation and analytical improvements to accurately segment cells were required to generate high-quality MERFISH data in lung tumors (**Methods**) (*108–113*). The resulting data showed high correlation between replicates and with bulk RNA-seq of 4T1 lung metastases as well as pseudo-bulk scRNA-seq of matched cell types (**fig. S9B-D**) (*114*). Transcriptional clusters were annotated based on the expression of previously-described marker genes (*105–107*) to distinguish 18 cell types (**Fig. 5D,E** and **fig. S9E,F**). Overall, we noted consistency in the set of cell types that are present in each tumor but variability in cell type frequency related to tumor size, with relatively less immune cell infiltration for larger tumors (**Fig. 5D**). Larger tumors often had a necrotic core containing mostly cellular debris with few intact nuclei or RNA molecules detected by MERFISH (**Fig. 5C** and **fig. S9A**).

We next examined the neighborhood enrichment and spatial distribution of cell types within 4T1 tumors and in relation to the lung/tumor interface (**Fig. 5F,G**). As expected, we observed that AT1/AT2 cells, club cells, and endothelial cells were enriched in the non-tumor, normal lung tissue (**Fig. 5F-H**). The majority of immune cell types, including CD11c^hi^ (Integrin alpha X) macrophages, were enriched near the lung/tumor interface (**Fig. 5G,H**). Immune cell types with distinct spatial distribution patterns included neutrophils which were enriched in the non-tumor lung and ARG1^hi^ (Arginase 1) macrophages which were enriched at the outer tumor boundary furthest from the adjacent lung (distal tumor boundary; **Fig. 5G,H**), as described previously (*115*). Spatial distribution patterns also varied between fibroblast subsets. Alveolar fibroblasts 1 and 2 (AF1 and AF2) were enriched at the lung/tumor interface and non-tumor lung, respectively (**Fig. 5G**). Cancer-associated fibroblasts were enriched at the distal tumor boundary, where we also observed ARG1^hi^ macrophage enrichment and reduced endothelial cell density, consistent with prior descriptions of M2-like macrophages enriched in hypoxic tumor regions (*116–119*) **(Fig. 5G,H**). These results reveal the intricate spatial organization of diverse cell types within the tumor microenvironment in this syngeneic model of cancer metastases.

#### Lineage maps reveal spatial patterns of metastatic tumor growth

To dissect how cell lineage informs tissue structure, we further processed the same tumor sections profiled by MERFISH and applied our optimized in-gel T7 amplification protocol to read out lineage information (**Fig. 6A**). We first validated the consistency in decoding intBC and LM *in vivo* by imaging a primary 4T1 tumor comprising fully-edited cells, where the optimizations used for *in vitro* samples enabled robust LM readout in tumor sections with 99.6% accuracy (**fig. S10**). For tumor sections, we noted a relationship between intBC detection efficiency, nuclei volume, and the z position of the nuclei in the 3D image (**Fig. 6B**). The decreased volume of nuclei at the top and bottom of the tissue slice indicates that these nuclei were bisected during sectioning, which limits intBC detection efficiency because integrations are distributed throughout the 3D volume of the nucleus. Therefore, we only included nuclei with z positions 4 μm (corresponding to the average nuclei diameter) from the edge of the tissue section for which the average intBC detection rate per cell was 55.2±9.7% (**fig. S11A** and **table S23**). Across 11 clones (from 10 tumors profiled; M2 T2 contained two founding clones) the number of edit sites ranged from 36 to 63 (**fig. S11B**). Four phylogenetic clones with mean edit fractions >50% were selected for further analysis (**fig. S11C**). As with the *in vitro* 4T1 colonies, we reconstructed phylogenies with neighbor joining using all cells with >60% intBCs detected, on average 45.2% of whole malignant cell nuclei (range 35.8%-57.5%; **Fig. 6C** and **fig. S11D**). After this stringent quality filtering to ensure accurate tree reconstruction, we obtained four phylogenies across up to four sections from these tumors. *In vivo* trees range in size from 1,404 (Mouse 2 Tumor 5; M2 T5 section 1) to 19,029 cells (Mouse 1 Tumor 1; M1 T1 sections 1-4) with near optimal editing kinetics and LM diversity consistent with editing rates observed in these cells *in vitro* (**Fig. 6D,E** and **fig. S11E,F**).

**Fig. 6.**
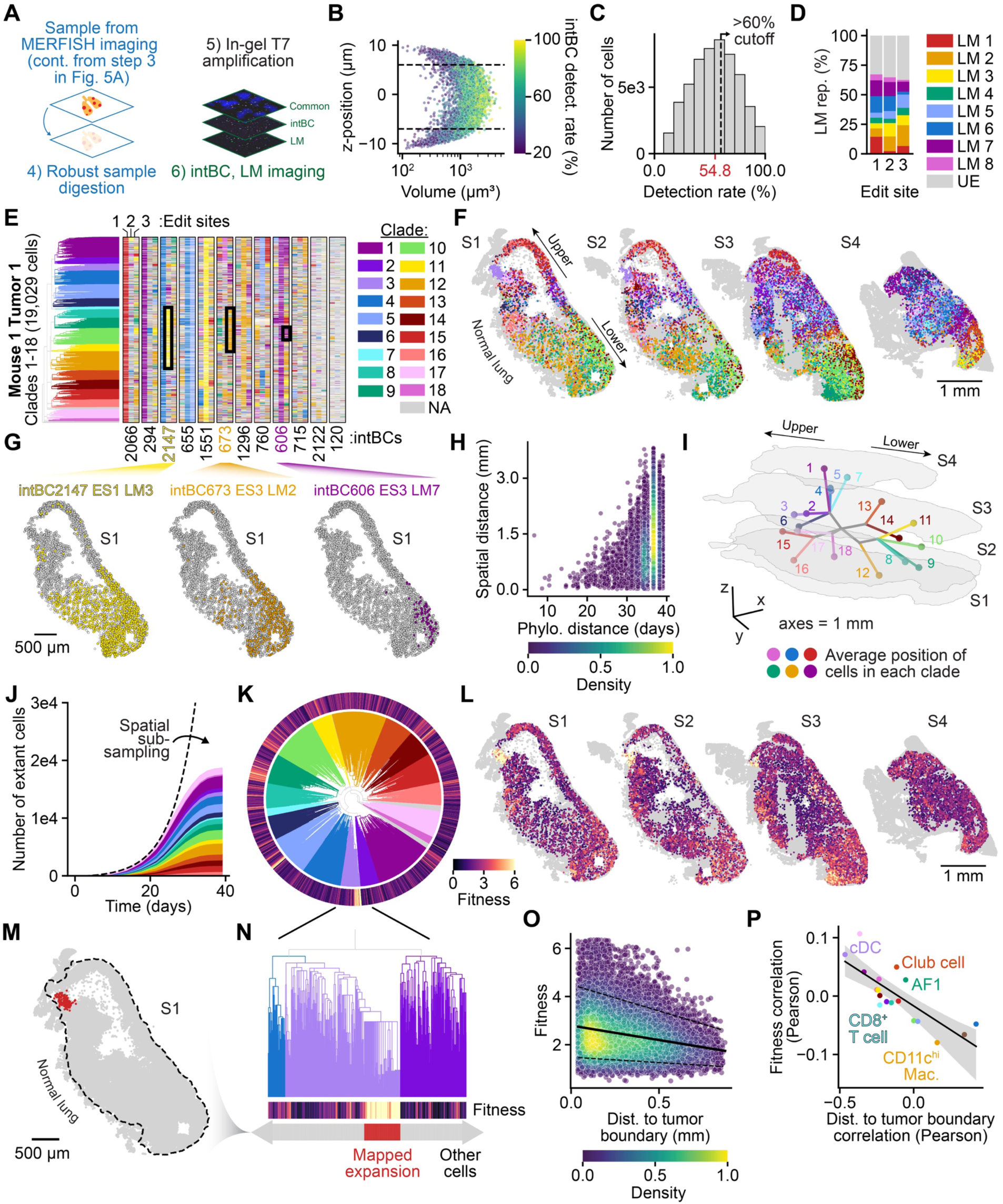
*In vivo* phylogenetic reconstructions at tissue scale. (**A**) Experimental schematic for imaging-based readout of PEtracer lineage data from samples after completion of MERFISH RNA readout. Samples undergo extensive digestion, in-gel T7 *in situ* amplification, and imaging of PEtracer integration barcodes (intBCs) and lineage marks (LMs). (**B**) intBC detection (detect.) rate as a function of cell volume and its z-position. (**C**) Histogram of cell count by *in vivo* intBC detection rate for M1 T1. The average detection rate is shown in red; a 60% cutoff for cells included for phylogenetic reconstructions is noted as a dashed line. (**D**) Edit site LM modification representation across all cells included in the phylogeny for M1 T1. UE = unedited. (**E**) Character matrix and phylogeny for the 19,029 cells with >60% intBC detection efficiency across four sections (S1 to S4) for M1 T1. intBCs are listed below the character matrix which is colored by LM identity for each edit site. LM coloring same as in (D), where additionally white = not detected. Clades 1 through 18 are colored in the phylogeny. NA = not assigned. Clade color scheme shared with (F). (**F**) Cells from four sections (S1-S4) colored by clade assignment. The location of the normal lung and the upper and lower portions of the lesion are annotated. Scale bar = 1 mm. (**G**) Spatial organization of edits in M1 T1 S1 that define phylogenetically related spatial domains. Grey cells denote other LM or unedited, white cells denote not detected. ES = edit site. (**H**) Pairwise phylogenetic (Phylo.) versus spatial distance for M1 T1 phylogeny. (**I**) Three-dimensional reconstruction of the colored clades in M1 T1. Balls indicate the average spatial position of cells in a clade; branches indicate the average position of contributing ancestors to each node. Sections S1 to S4 depicted as light outlines. Axes lengths = 1 mm. (**J**) Estimated number of extant cells as a function of time. Each clade is colored as in (E); dotted line shows an exponential fit curve of tumor growth. (**K**) Phylogeny from (E) for M1 T1 cells across all four sections depicted as a circle with phylogeny branches colored by clade assignment. Outer ring depicts inferred fitness for each cell. (**L**) Spatial organization of cell fitness across four sections for M1 T1. A spatially-confined (**M**), high-fitness expansion (**N**) is observed at the lung/tumor boundary. Cells in this expansion in S1 are shown in red, other cells shown in grey. (**O**) Malignant cell distance (dist.) to the tumor boundary versus fitness. Black line shows regression, dotted line mark top and bottom deciles. (**P**) Pearson correlation between indicated cell type density with fitness versus correlation of cell type density with distance to tumor boundary. Black line depicts regression with ribbon indicating 95% confidence interval. Mac. = macrophage, AF1 = Alveolar Fibroblast 1, cDC = conventional Dendritic Cell. See table S24 for relevant statistics.

With these tissue-scale phylogenies reconstructed using the continual accrual of LMs, we could study the coordinated effects of cell state, lineage, and spatial organization on tumor evolution *in vivo*. Phylogenetic clades occupy distinct spatial regions with consistent patterns observed across multiple sections and tumors (**Fig. 6E,F** and **fig. S11G**). In tumor M1 T1, for example, intBC2147 edit site 1 LM3 (ES1 LM3) marked an early split in the tree, with cells bearing this LM predominantly located in the lower half of the tumor, suggesting an early axis of tumor growth parallel to the lung/tumor interface (**Fig. 6G**). Similar to the *in vitro* colonies, LMs defining later branches in the tree–representing cells with more recent common ancestors–are confined to progressively smaller spatial regions (**Fig. 6G**). Notably, the correlation between pairwise phylogenetic distance and maximum spatial distance was higher than *in vitro* colonies, possibly due to the increased cell density and size of the tumor reducing the extent and impact of local cell movement (**Fig. 6H**). However, spatial intermixing of distinct phylogenetic clades indicates that 4T1 cells remain motile *in vivo* (**Fig. 6F,H** and **fig. S11G**). Cells with more similar sets of LMs (lower Hamming distances) also tended to be spatially close, though this trend was weaker than that observed for phylogenetic distance (**fig. S12A,B**).

To reconstruct the trajectory of tumor growth in three dimensions, we integrated lineage information at multiple levels of the reconstructed tree with the spatial distributions of profiled cells to calculate the average spatial location of cells within each phylogenetic clade, as well as the average spatial locations of ancestral nodes in the tree (**Fig. 6I**). This global analysis nominated an early phylogenetic and spatial bifurcation of tumor growth, with clades 1-7 located in the upper half of the tumor and clades 8-12 in the lower half. Clades 17 and 18 represent a phylogenetic outgroup with rich lineage sub-structure that is spatially enriched at the lung/tumor interface in the upper region of the tumor, a region with high local phylogenetic diversity (**Fig. 6F,I** and **fig. S12C-E**). The high clade diversity and the presence of this subclonal outgroup nominate this region near the lung/tumor interface as a possible origin of tumor growth. To understand the growth dynamics of phylogenetic clades over time, we estimated the number of extant cells for each clade by cutting the tree at various time points throughout tumor development (**Fig. 6J**). This analysis suggested exponential growth for all clades over the course of tumor development, consistent with mostly neutral evolution at the clade level. We expect that the divergence from exponential expansion at later time points is caused by subsampling due to collection of a limited number of tumor sections despite high cell recovery for individual sections, although subsampling could also mask changes in growth rates later in tumor development due to spatial and nutrient availability constraints. Together, these data provide a high-resolution view of tumor growth over time in three dimensions.

To understand the drivers of cell lineage dynamics *in vivo*, we used tree structure to assess phylogenetic fitness. Cells with heritable increases in fitness will undergo more cell divisions relative to other sub-clades over time, resulting in more extensive branching and increased sampling of descendants. We calculated fitness for each cell based on the branch lengths of its ancestors and relatives in the tree (**Methods**) and examined the spatial distribution of high-fitness cells (**Fig. 6K,L**) (*82*). We find that cells with high fitness are enriched along the periphery of the tumor adjacent to the normal lung, including a highly localized expansion at the lung/tumor interface (**Fig. 6M,N**). Overall, we observed a negative correlation between phylogenetic fitness and distance to the tumor boundary (**Fig. 6O**). We also confirmed that this trend could be reproduced using a tree-independent metric of fitness assessed by mean neighbor LM distance (**fig. S12F-H**). Given the diverse spatial distribution patterns of immune and stromal cells, we asked whether local cell type densities correlated with cancer cell fitness. Although the density of several cell types, including conventional dendritic cells (cDCs), were weakly correlated with fitness, we observed correlation between fitness-associated cell type densities and distance to the tumor boundary, complicating efforts to disentangle the effects of cell type density from spatial position (**Fig. 6P**, **fig. S12I-K,** and **table S24**). The wide variability in cancer cell phylogenetic fitness we observed at the tumor boundary suggests that additional cell-intrinsic and cell-extrinsic factors may cooperate to influence cancer cell dynamics and fitness.

#### Transcriptional and spatial determinants of cancer cell fitness

We designed a new MERFISH library to further assess gene programs associated with cancer progression including hypoxia, epithelial-mesenchymal transition (EMT), and cell cycle progression to better characterize transcriptional and spatial features that underlie differences in cancer cell fitness. Using this 175-gene library, we generated MERFISH data from an additional tumor sample assaying three sections with 104,219 total cells and an average of 261 transcripts per cell (Mouse 3 Tumor 1; M3 T1; **tables S22,25**). Our new library further resolved previously detected cell types, including macrophages and endothelial cells, into transcriptionally-distinct subtypes while still recovering all major annotations from our original library (**fig. S13A-D**). With this new library, we identified four spatially-varying malignant cell transcriptional modules with distinct cellular neighborhood composition using Hotspot (*120*): 1) “leading edge” cells at the distal tumor boundary, 2) “hypoxic” cells, 3) “tumor core” cells, and 4) “lung adjacent” cells (**Fig. 7A-C** and **fig. S13E,F**). Notably, grouping cancer cells into these modules also captured features of EMT with increased expression of mesenchymal-associated genes *Snai1* and *Sox9* in module 2/hypoxic cancer cells (**Fig. 7B**).

**Fig. 7.**
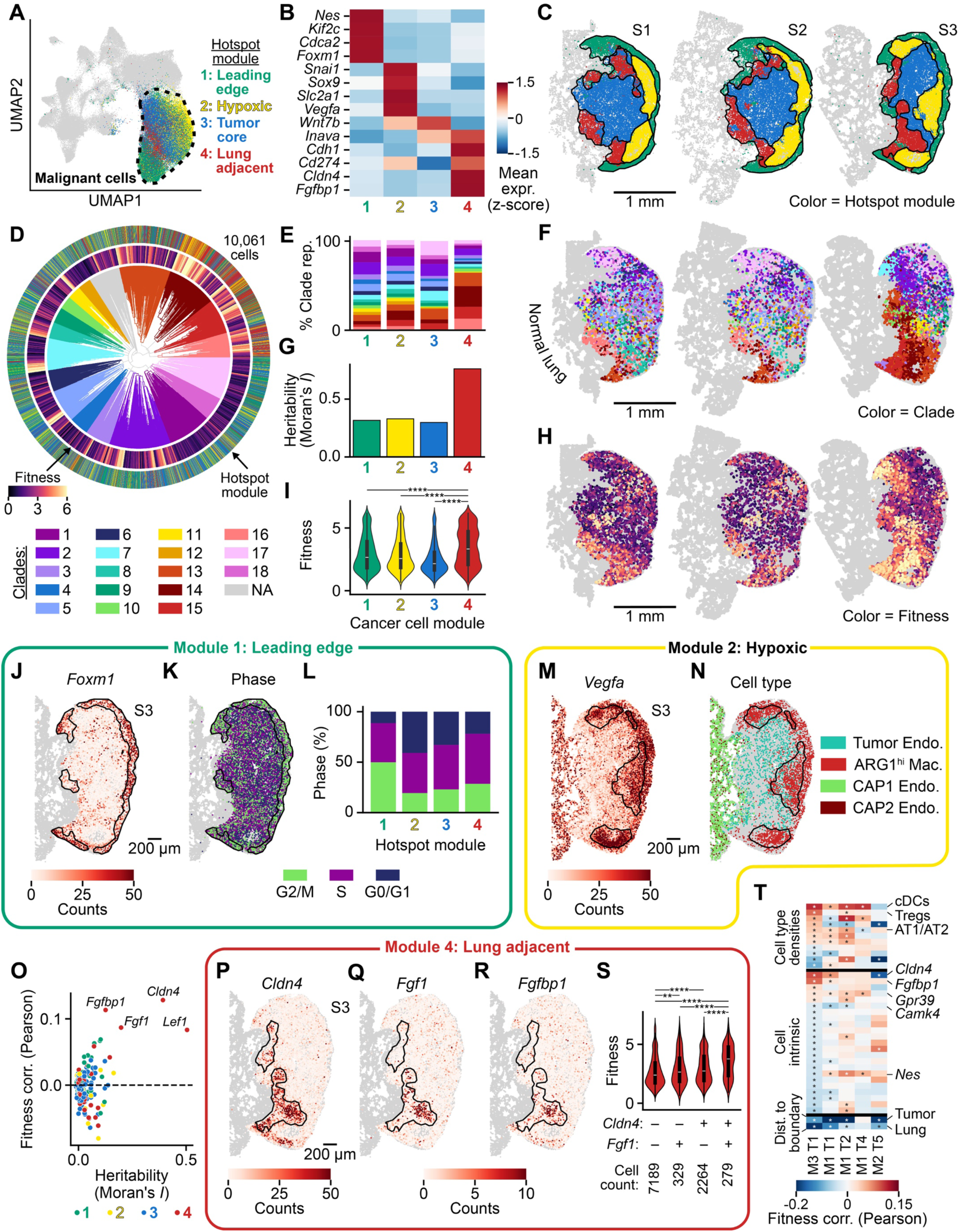
High resolution spatial mapping of cell state and lineage uncovers drivers of tumor growth. (**A**) UMAP embedding of 104,219 cells sampled across three sections (S1 to S3) of a tumor from Mouse 3 (M3 T1) where the malignant cells are outlined and colored by their assignment to four distinct Hotspot modules: Module 1/Leading edge, Module 2/Hypoxic, Module 3/Tumor core, and Module 4/Lung adjacent. Full UMAP embedding where each cell is colored by its annotated cell type can be found in fig. S13A. (**B**) Heatmap of marker gene expression (expr.) for each Hotspot module. (**C**) Spatial organization of modules 1-4 across M3 T1 S1-S3. (**D**) Phylogeny for all malignant cells with >60% integration barcode detection efficiency across M3 T1 S1-S3 depicted as a circle. Branches are colored by clade assignment (Clades 1-18) with a color key beneath. The inner ring denotes fitness while the outer ring denotes the module assignment for each of the cells in the tree. (**E**) Representation (rep.) of each of the 18 phylogenetic clades across the 4 Hotspot modules. (**F**) Spatial organization of phylogenetic clades 1-18 across M3 T1 S1-S3. (**G**) Quantification of module assignment heritability as measured by Moran’s *I* phylogenetic autocorrelation. (**H**) Spatial organization of cell fitness scores across M3 T1 S1-S3. (**I**) Violin plots of cell fitness scores by module with embedded boxplots showing quartiles. <u>Module 1/leading edge cell analyses = (J-L).</u> (**J**) MERFISH counts for *Foxm1* transcripts in M3 T1 S3 (malignant cells only). (**K**) Spatial organization cell cycle phase in M3 T1 S3 (malignant cells only). (**L**) Quantitation of cell cycle phase by Hotspot module. <u>Module 2/hypoxic cell analyses = (M-N).</u> (**M**) MERFISH counts for *Vegfa* transcripts in M3 T1 S3 (all cells). (**N**) Spatial distribution of Tumor endothelial (Endo.), ARG1^hi^ macrophages (Mac.), and capillary 1/2 (CAP1/CAP2) endo. cells. (**O**) Heritability versus Pearson fitness correlation (corr.) of individual genes colored by module assignment. <u>Module 4/lung adjacent cell analyses = (P-S).</u> MERFISH counts for (**P**) *Cldn4*, (**Q**) *Fgf1*, and (**R**) *Fgfbp1* transcripts in M3 T1 S3. (**S**) Violin plot for module 4 malignant cell fitness as a function of *Cldn4* and *Fgf1* expression. Abundance of cells with varying states listed below each entry. (**T**) Pearson correlation of malignant cell fitness with cell type densities, cell-intrinsic gene expression, or tumor features (distance (dist.) to the tumor boundary or lung-tumor interface) across the 5 tumors analyzed in this study. cDC = conventional Dendritic Cell, Treg = T regulatory cell, AT1/AT2 = alveolar epithelial type 1/2. Scale bars = 1 mm for (C), (F), and (H). Scale bars = 200 µm for (J-K), (M-N), (P-R). Significance in (I) and (S) denotes Mann-Whitney-Wilcoxon two-sided test where *p < 0.05, **p < 0.01, ***p < 0.001, ****p<0.0001. Significance in (T) denotes Student’s t-test of Pearson correlation coefficient. See tables S24, S26, and S27 for relevant statistics.

To understand how malignant cell modules emerged during tumor growth, we constructed a phylogeny for the 10,061 cells with >60% intBC detection in M3 T1 (**Fig. 7D** and **fig. S13G**). The spatial organization of clades within this tumor is consistent with other tumors in this study (**Fig. 7F** and **fig. S13H**). Broadly, we observed that every phylogenetic clade was represented in each Hotspot module with mostly uniform clade compositions with the exception of module 4 which was enriched for clades 13-16 (**Fig. 7E-F**). When we examine module assignments across our phylogeny, we observe clear intermixing of malignant cell modules within phylogenetic clades, suggesting that 4T1 cells exhibit transcriptional plasticity that allows them to quickly adapt to local microenvironmental conditions (**Fig. 7D** and **fig. S13I**). Despite this broad plasticity, we find striking differences in malignant cell module heritability. Specifically, module 4/lung adjacent cells showed a strong propensity to remain in that state (**Fig. 7G** and **table S26**). Module 4/lung adjacent cells express the highest level of epithelial markers such as *Cdh1*, encoding E-cadherin, which may underlie their tendency to remain in this state and location unlike leading edge cells which upregulate mesenchymal markers and therefore are more likely to move and exhibit transcriptional plasticity (**Fig. 7B**) (*121*). We next used the M3 T1 phylogenetic tree to parse how these varied factors were integrated to instruct cancer cell fitness. As in M1/M2 tumors, cells with high fitness were overrepresented in certain phylogenetic clades and exhibited clear enrichment at the tumor boundary, particularly within the module 4/lung adjacent module (**Fig. 7D,H,I**).

Together, these data suggest that malignant cell modules reflect the integrated effects of transcriptional programs and environmental cues that instruct tumor growth. Among module 1/leading edge cells at the distal boundary of the tumor, we observed higher expression of cell cycle-related genes including *Foxm1*, a cell cycle-associated transcription factor previously identified as a genetic dependency for 4T1 cell growth *in vivo* (**Fig. 7J-L**) (*122*). Cancer cells in module 1 occupy a cellular neighborhood enriched for CAFs, as we observed at the distal boundary in other tumors in this study (**Fig 5H** and **fig. S13F**). The spatial and transcriptional signature of the module 1/leading edge cells has low heritability indicating that the proliferative capacity of these cells is a transient state likely influenced by microenvironmental factors that do not instruct long-term, heritable behaviors (**Fig. 7E,G**). Underneath the leading edge of cycling cells, a hypoxic microenvironment defines module 2. Expression of hypoxia markers (e.g. *Vegfa*) are high in module 2 and correlate with cell type composition, notably a sharply demarcated boundary between tumor endothelial cells and ARG1^hi^ macrophages (**Fig. 7M,N**) (*116–119*). Malignant cells within the module 3/tumor core show no evident transcriptional or lineage signatures. By contrast, module 4/lung adjacent cells have the highest fitness, and this module is the most heritable in this tumor (**Fig. 7G,I**). Phylogenetic fitness reflects proliferative potential across several weeks and many cell divisions based on estimated branch lengths while cell cycle state can fluctuate in a matter of hours, which we suspect is why module 4/lung adjacent cells have higher fitness than module 1/leading edge cells despite having a slightly lower proportion of actively cycling cells (**Fig. 7L**).

We next sought to disentangle the varied effects of lineage, positional, cell-intrinsic, and cell-extrinsic factors that likely contribute to the characteristics of cells in module 4. Similar to our observations in M1/M2 tumors, we found that the densities of several cell types (i.e. cDCs, Tregs) were positively correlated with phylogenetic fitness and also enriched at the tumor boundary in the neighborhood of module 4/lung adjacent cells (**fig. S13F,J**). To identify cell-intrinsic drivers of fitness that coordinate with the microenvironment to promote growth, we looked for genes associated with fitness that also exhibited heritable expression patterns. This analysis nominated several heritable genes that may underlie cell-intrinsic fitness programs within module 4, including *Cldn4* (Claudin 4) and *Fgf1*/*Fgfbp1* (**Fig. 7O-R** and **tables S24,26**). *Cldn4*, which encodes an epithelial cell tight junction protein, is broadly expressed by module 4 malignant cells but lowly expressed elsewhere (**Fig. 7P**). In contrast, *Fgf1*/*Fgfbp1*, which encode Fibroblast Growth Factor 1 (FGF1) and its associated carrier protein Fibroblast Growth Factor Binding Protein 1, are expressed in a highly-specific spatial region of the tumor, with their expression being less heritable than that of *Cldn4* (**Fig. 7O,Q,R**). These genes illustrate how spatially- and phylogenetically-coherent programs interact to promote fitness within module 4, where cells co-expressing both *Cldn4* and *Fgf1* exhibit the highest phylogenetic fitness (**Fig. 7S**). We also found that *Fgf1* is expressed by non-malignant cells (**fig. S13C**), suggesting potential roles for both paracrine and autocrine signaling. Indeed, the high heritability and fitness of cells in module 4 is associated with a unique cellular neighborhood and thus signaling environment that may reinforce the transcriptional programs adopted by malignant cells to thrive in this niche.

Having nominated features that underlie cancer cell fitness in M3 T1, we revisited previous samples and were able to observe reproducible factors that govern tumor evolutionary programs (**Fig. 7T** and **table S24**). For nearly all tumors profiled, spatial location within the tumor–as assessed by distance to the tumor boundary or tumor/lung interface–is the best predictor of phylogenetic fitness. This metric of spatial organization co-varies with multiple features including space to grow, endogenous transcriptional programs, and cellular neighborhood (**fig. S13J**). We find that multiple cell-extrinsic and cell-intrinsic features are reproducibly correlated with fitness, including *Cldn4* and *Fgfbp1* expression (**Fig. 7T** and **fig. S13K**), suggesting that permissive microenvironmental effects coordinate with cell-intrinsic drivers of fitness to promote transition to a heritable, high fitness state. Together, these data provide a high-resolution view of evolutionary dynamics in both time and space, providing a robust toolkit to describe how cell-intrinsic, cellular neighborhood, and broader constraints–such as nutrient availability and space for cell division– inform the evolutionary dynamics of cells.

## Discussion

Here we describe PEtracer, an evolving lineage tracing technology that enables joint phylogenetic and cell state profiling using either sequencing or high-resolution imaging that preserves cellular context in tissues. Guided by computational modeling, we engineered the PEtracer system from first principles to enable robust, high-resolution phylogenetic reconstructions based on LMs installed across diverse biological timescales. We adapted prime editing for rapid lineage tracing capable of marking individual cell divisions for rapidly proliferating cells and tuned editing kinetics to enable lineage recording over many weeks or months. In contrast to approaches that install barcodes at discrete times, the nature of evolving LM installation with PEtracer allows us to capture key events that occur dynamically throughout a biological process of interest without *a priori* knowledge of when these events might occur. We performed rigorous benchmarking of our system to establish that we can assemble high-quality phylogenies that accurately reconstruct static barcode groups across multiple time points. Using sequencing and highly-multiplexed, multimodal imaging technologies with efficient molecular and cellular detection, we collected rich cell state and lineage data from intact tissue sections, which provided a detailed view of local tumor evolutionary dynamics. We reconstructed the growth patterns of individually-seeded tumors in the 4T1 transplantable model of murine breast cancer lung metastasis (*98*, *99*, *104*), demonstrating our ability to build phylogenies comprising tens of thousands of cells while retaining the rich cellular context within the tumor microenvironment.

Our tumor metastasis studies highlight how the integration of cell state, lineage, and spatial information at tissue scale *in vivo* can provide an unprecedented view of tumor evolution in time and space. With imaging-based spatial transcriptomics, we assign cancer cells to spatially- and transcriptionally-coherent modules defined by distinct cell-intrinsic and cell-extrinsic programs. At the outer tumor edge distal from the normal lung, we observe malignant cell division likely driven by the availability of room for further growth and a supportive cellular neighborhood rather than heritable, cell-intrinsic factors, as suggested by phylogenetic analysis, which shows low heritability of this state. Interestingly, phylogenetic fitness and cell cycle state seem to reflect proliferative capacity over different time scales in this model, as more proliferative cells at the leading edge do not have the highest fitness within the tumor. Instead, as leading edge cells grow outward, they leave behind a high density layer of cells where low oxygen availability induces a hypoxic state, upregulation of mesenchymal gene expression programs (*123*), and high *Vegfa* expression, which may ultimately promote angiogenesis to provide necessary nutrients. Regions where cancer cell proliferation outpaces the recruitment of blood vessels may lead to the areas of necrosis we observe in some tumors. Within the more vascularized tumor core, cellular fitness appears to be constrained by a lack of space for continued growth despite an available blood supply. In contrast, epithelial-like malignant cells adjacent to the normal lung grow in an nutrient- and immune-rich niche and have the highest phylogenetic fitness, indicating a propensity for tumor cells in this region to undergo a heritable change to a high-fitness state (*124*). We identified specific phylogenetic clades that thrive within this unique neighborhood, where some cells have co-opted distinct gene expression states (e.g. *Cldn4*, *Fgf1*/*Fgfbp1*) to drive high fitness. Notably, spatially-resolved phylogenies allow us to uncover complex relationships between transcriptional state, spatial location, and lineage that would be difficult to resolve with approaches relying on spatial or transcriptional neighbors to make lineage inferences due to cell migration and phenotypic plasticity. More broadly, our results elucidate transcriptional programs as well as spatial determinants that drive cancer cell fitness, and underscore the complex interplay between cell-intrinsic and environmental factors that ultimately shapes the trajectory of tumor growth and evolution.

These efforts motivate several new directions to further understand evolutionary relationships between cells in complex biological systems: i) While our work here describes a single model of cancer cell extravasation and metastatic outgrowth that has been preselected for metastatic ability, other tumor models may better capture cell state transitions and subclonal expansions during the early stages of tumor progression and may have distinct spatial patterns of growth. We envision using PEtracer to study many biological processes including cancer initiation, progression, therapeutic resistance, and the complete metastatic cascade as well as early development, regeneration, and homeostasis (*13–16*). ii) Assaying tissues with both imaging and sequencing readouts (*83*, *84*, *125*), would allow for joint lineage reconstruction, thereby facilitating full transcriptomic imputation based on phylogenetic neighbors (*58*, *126–128*). iii) Extending imaging efforts to profile other key aspects of cell state including protein localization and epigenetic state using the diverse suite of multimodal, multiplexed FISH and/or immunofluorescence-based measurements (*57*, *129*, *130*). iv) Methodological adaptations to assay thicker tissue sections in combination with emerging computational approaches would better enable three-dimensional reconstructions of the architectures of sampled tissues (*131*). v) Implementing novel lineage reconstruction algorithms (*e.g.* scalable maximum likelihood methods) may improve the quality of phylogenies generated with PEtracer (*132–135*). vi) Layering perturbations on top of phylogenetic reconstructions would enable direct interrogation of the genetic factors hypothesized to underlie cell state transitions and evolutionary dynamics (*69*). vii) Integrating emerging prime editor-based signal recording methodologies (*136*) would allow for decoration of phylogenies with LMs associated with signaling events of interest to assess their timing and impact on fate determination. With a recording space of 1,024 possible 5nt LMs per edit site, we can theoretically record hundreds of biological signals.

Understanding how cellular histories shape tissue-scale characteristics is a longstanding challenge in biology with broad biological significance. While current cell profiling methods and associated atlases provide an exceptional foundation for understanding these phenomena (*8*, *137–146*), they do not directly probe phylogenetic relationships between cells, which are necessary for deciphering cell fate determination and elucidating the principles of tissue organization. We propose that the PEtracer system will provide a foundation for probing cellular dynamics across diverse biological contexts including additional cancer models, development, stem cell dynamics, and tissue regeneration.

**Fig. S1.**
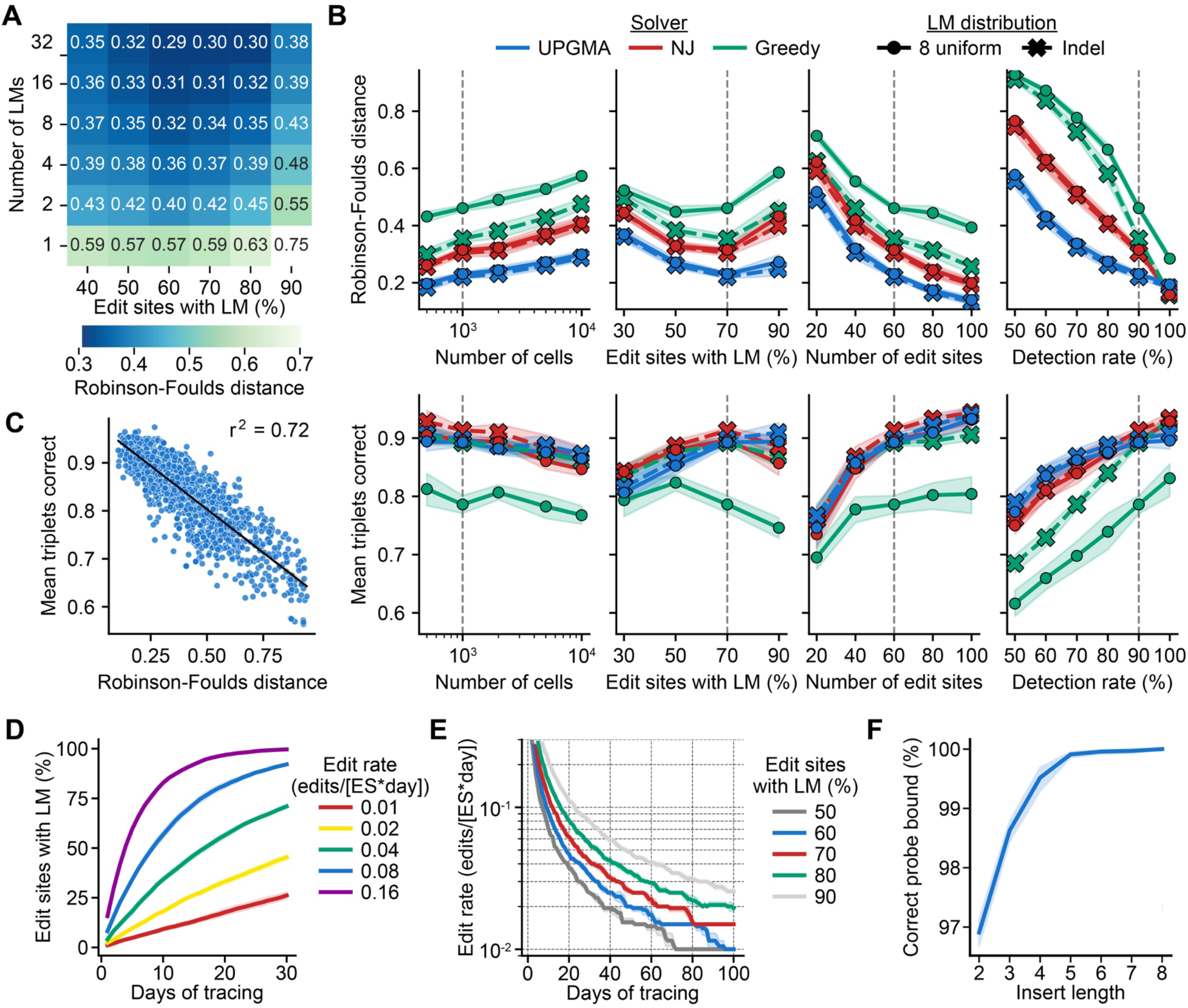
*In silico* modeling of key lineage tracing parameters. (**A**) Mean Robinson-Foulds reconstruction error for simulated phylogenies varying the number of lineage marks (LMs) and fraction of edit sites with an LM installed. (**B**) Mean Robinson-Foulds reconstruction error (top) and mean depth-normalized triplets correct (bottom) reconstruction accuracy for simulated phylogenies with LMs drawn from either an indel distribution with n = 9,662 LMs (crosses) or a uniform distribution with eight LMs (dots). Trees were simulated varying key experimental parameters: the number of cells (far left), fraction of edit sites with an LM (middle left), the number of edit sites (middle right), and detection rate (far right) and reconstructed using UPGMA (blue), Neighbor joining (NJ; red), and Greedy (green) algorithms. The Greedy solver was less accurate in reconstructing trees with eight LMs due to increased homoplasy at individual edit sites, which rendered greedy splits suboptimal; in contrast, methods leveraging information from all edit sites– such as UPGMA and NJ–remained robust to homoplasy. Values held constant for simulations where the experimental parameter was not varied are marked with grey dashed lines. (**C**) Correlation of Robinson-Foulds reconstruction error and mean depth-normalized triplets correct accuracy for simulated phylogenies. Black line depicts regression with ribbon indicating 95% confidence interval. (**D**) Simulated LM accumulation at edit sites with varying edit rates (edits/day). (**E**) Identification of optimal edit rates for lineage tracing experiments of varying durations. Optimal edit site saturation lies between 60-80% (colored); in some cases, lower edit site saturation (50%) or higher (90%) may still allow for accurate reconstructions. (**F**) *In silico* prediction of correct probe hybridization frequency as a function of LM insert length across ten simulated experiments with randomly sampled insert sequences. All simulations were run for ten iterations, with mean value reported; standard deviations shown as ribbons where appropriate.

**Fig. S2.**
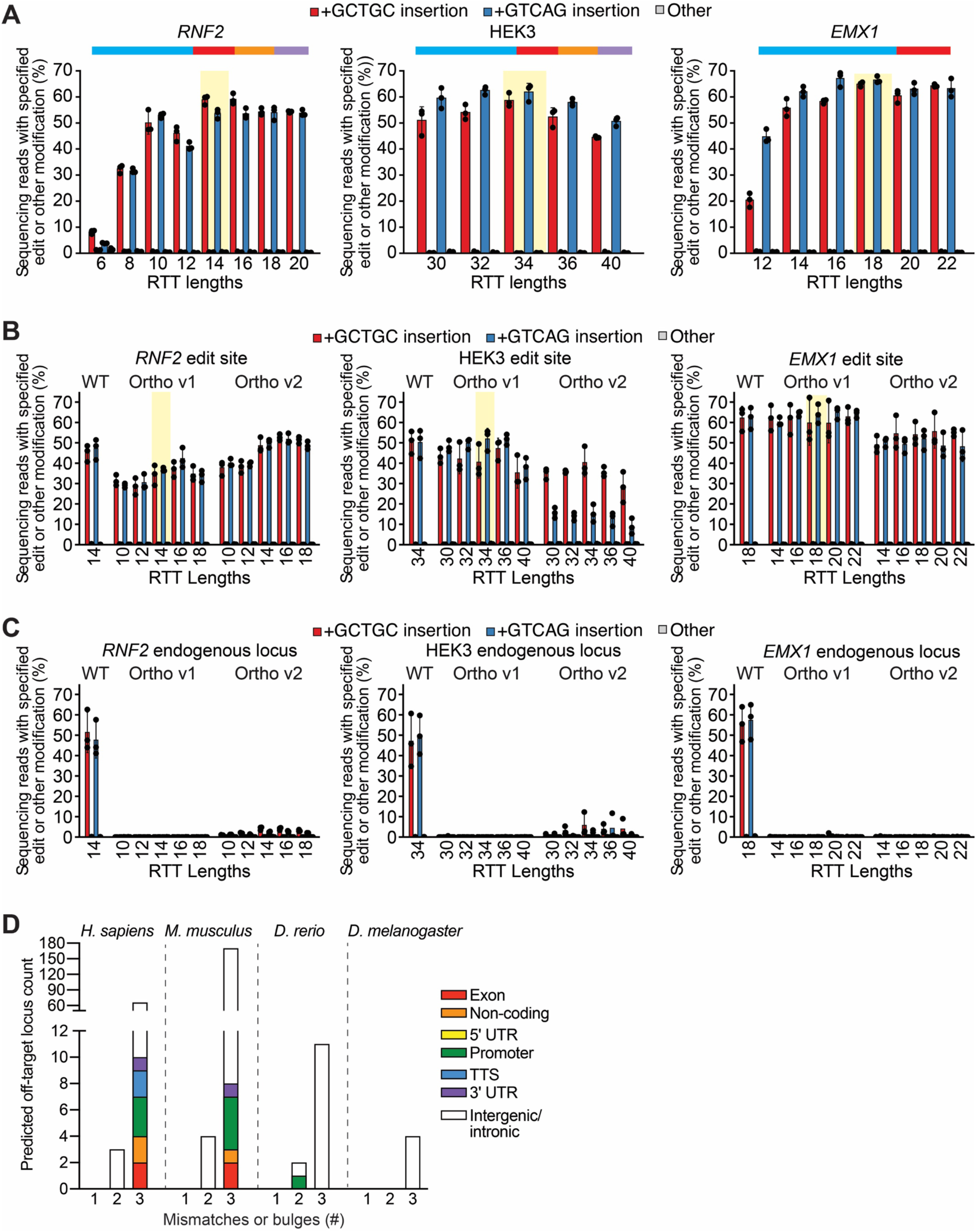
Edit site optimization and orthogonalization. (**A**) Fine-grained optimization of reverse transcription template (RTT) lengths for epegRNAs installing two representative five nucleotide (5nt) lineage marks (LMs) at endogenous loci in HEK293T cells. Optimal RTT lengths for each edit site sequence highlighted in yellow. (**B**) Editing efficiencies for representative LMs at tracing cassettes comprising *RNF2*, HEK3, and *EMX1* edit site sequences from the human genome (WT) or orthogonalized in two ways (Ortho v1–seed sequence complemented, v2–seed sequence reverse complemented). Editing efficiencies for each edit site variant and its cognate epegRNA with varying RTT lengths are shown (**C**) Editing of the endogenous *RNF2*, HEK3, and *EMX1* genomic loci in HEK293T cells when using WT, Ortho v1, or Ortho v2 epegRNAs. Mean of three biological replicates ± standard deviation depicted for (A-C). (**D**) *In silico* screening of potential genomic off-targets in *H. sapiens*, *M. musculus*, *D. rerio*, and *D. melanogaster* for Ortho v1 epegRNAs. Off-targets listed by species and sub-categorized by the number of mismatches or bulges in these off-targets. 18 putative off-targets in human and mouse non-intergenic regions were manually inspected for RTT homology and classified as unlikely to support prime editing. UTR = untranslated region. TTS = transcription termination sites.

**Fig. S3.**
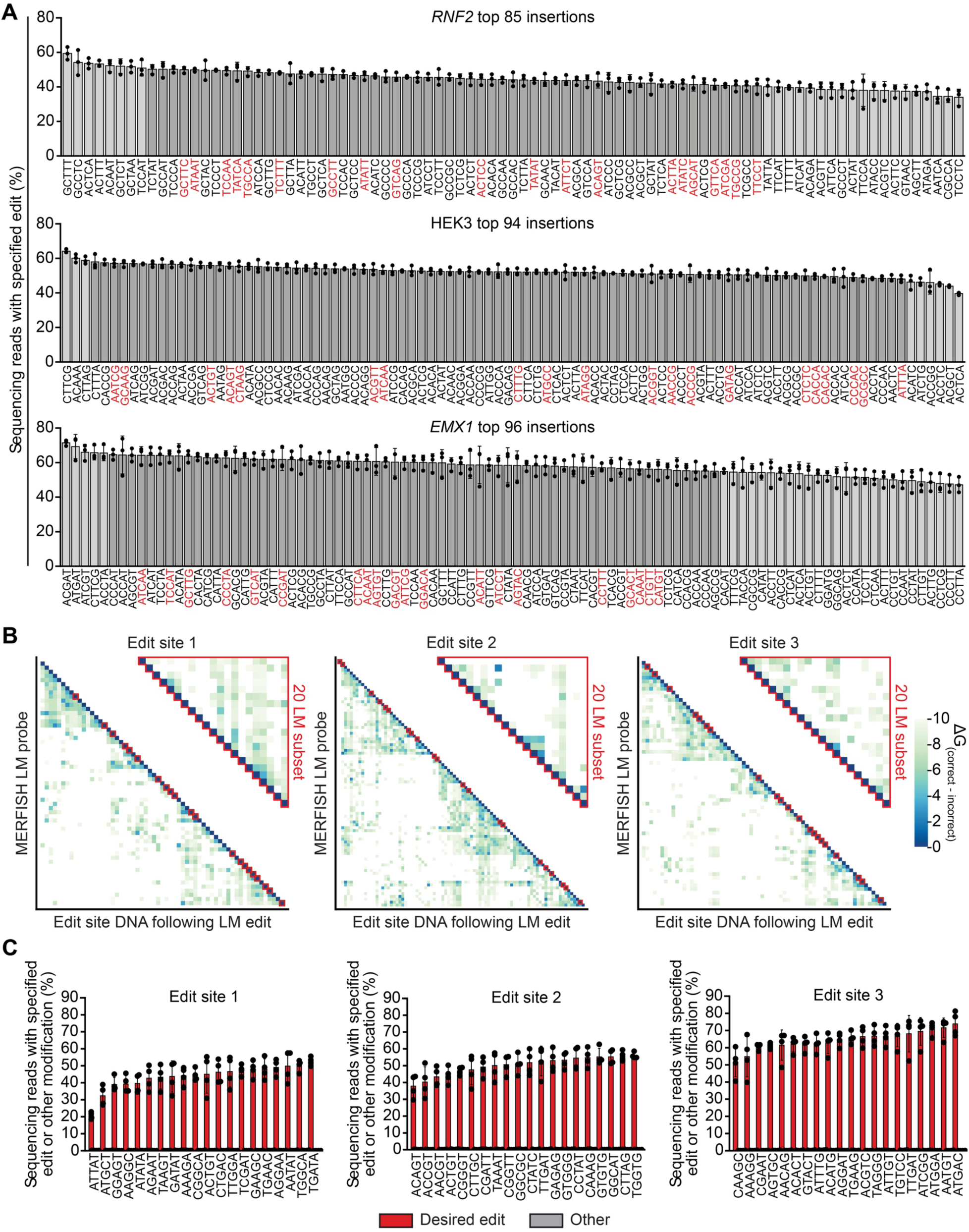
Screening for balanced and orthogonal LMs. (**A**) Installation efficiencies for best-performing five nucleotide (5nt) lineage marks (LMs) for each edit site. Mean of three biological replicates ± standard deviation depicted. *RNF2* = 85 LMs, HEK3 = 94 LMs, *EMX1* = 96 LMs. Complete set shown, those within 10% efficiency used for detection specificity measurements in (B) highlighted with dark grey bars and final 20 candidate 5nt LMs retested at orthogonal edit sites in (C) highlighted with red text. **(B)** Predicted probe hybridization specificity measurements (Gibbs free energy = ΔG) for discriminating all best-performing 5nt LMs from one another (bottom) or selected 20 LM subset with reduced cross-hybridization (top). (**C**) Editing efficiency of 20 LM subset for each edit site with optimized cross-hybridization retested at orthogonal edit sites in HEK293T cells. Mean of four biological replicates ± standard deviation depicted.

**Fig. S4.**
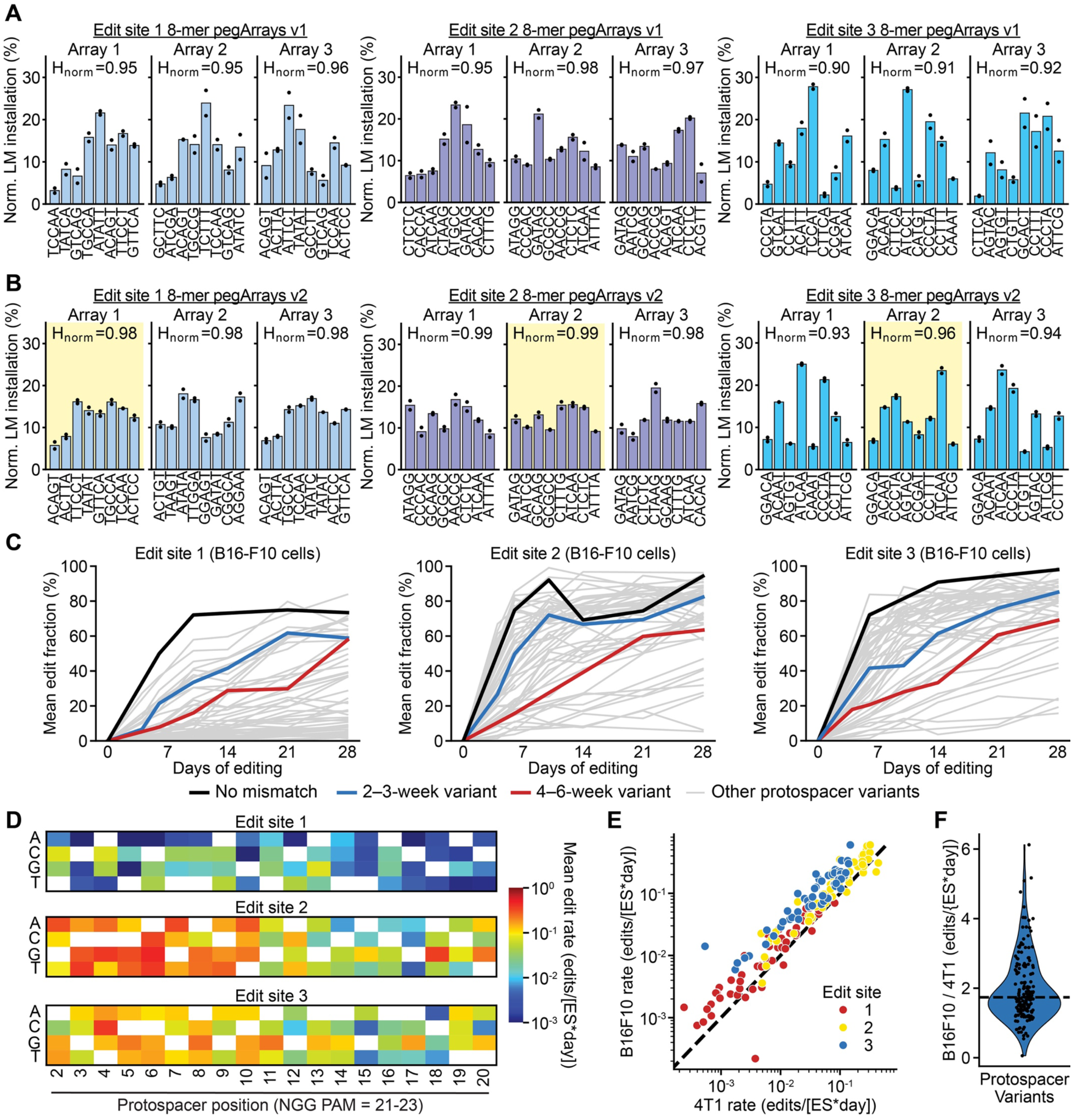
pegArray optimization and kinetic tuning. (**A**) Normalized (Norm.) lineage mark (LM) installation efficiencies for the first iteration (v1) of 8-mer pegArrays for edit sites 1, 2, and 3 in B16-F10 cells. Three arrays were tested per edit site. H_norm_ values provided for each 8-mer. Five nucleotide (5nt) LM sequence listed under its position in the 8-mer. (**B**) Normalized LM installation efficiency for the second iteration (v2) of 8-mer pegArrays for edit sites 1, 2, and 3 in B16-F10 cells averaged across two biological replicates. The final 8-mer used in the concatenated 24-mer is highlighted in yellow. Two biological replicates for (A) and (B). (**C**) Editing kinetics for protospacer mismatch variants at edit sites 1, 2, and 3 in B16-F10 cells. Labeling as in Fig. 2E; grey lines show all protospacer variants tested; no mismatch protospacers shown in black; protospacer mismatches selected for 2-3-week timescales shown in blue; 4-6-week protospacer mismatches in red. (**D**) Mean estimated edit rate associated with indicated protospacer mismatches along the length of the protospacer for each edit site. Protospacer mismatches decrease editing rates across three orders of magnitude in two tested MMR-competent cell lines (B16-F10 and 4T1) with slower editing rates for mismatches closer to the protospacer adjacent motif (PAM). The original base at each position is a white box in the heatmap. (**E**) Editing rate comparison for each variant in 4T1 and B16-F10 cells. Points colored by edit site. (**F**) Edit rate increase in B16-F10 cells relative to 4T1 cells for each protospacer variant with dashed line indicating median increase.

**Fig. S5.**
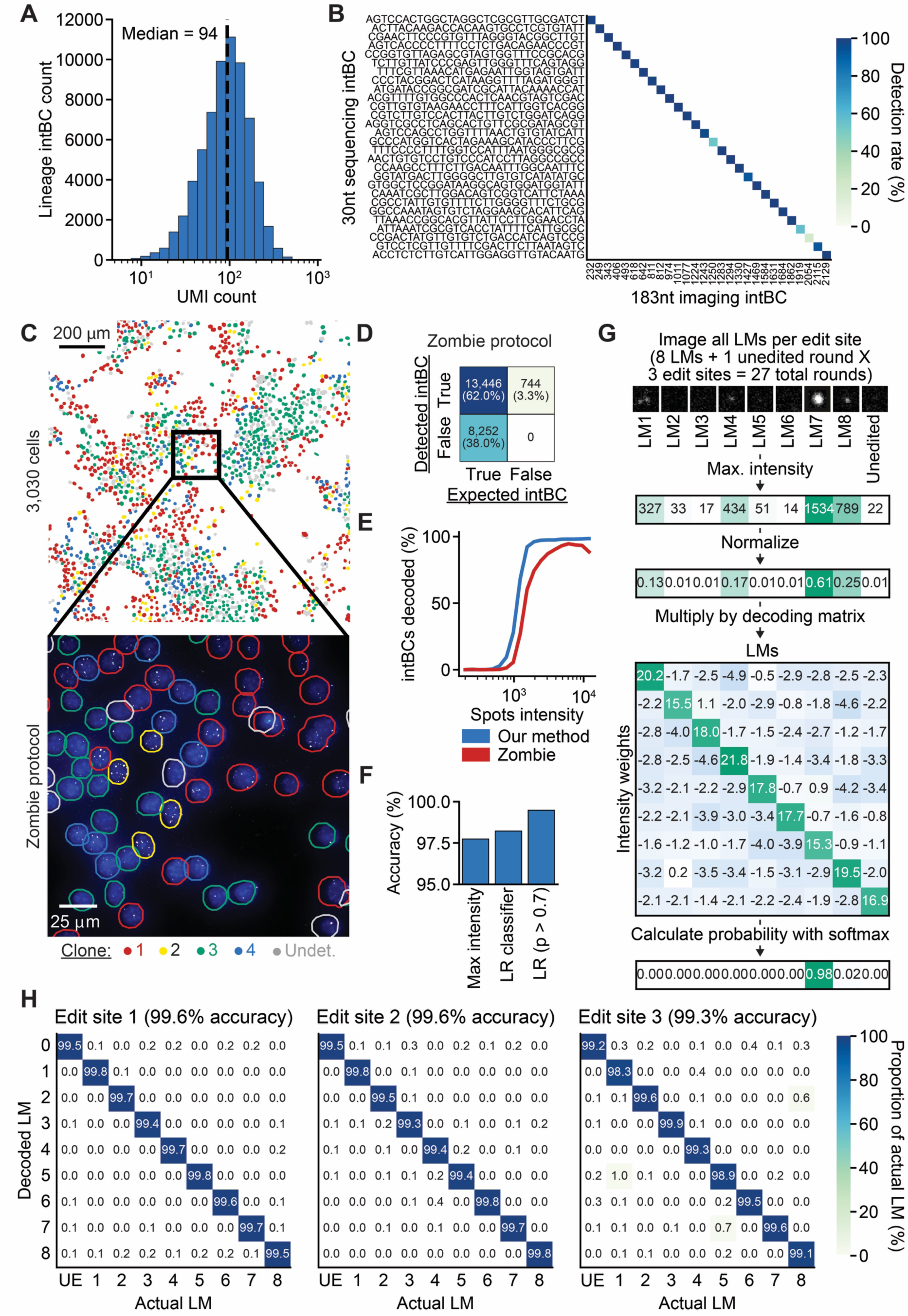
Optimization of *in vitro* PEtracer validation experiments. (**A**) Histogram of unique molecular identifiers (UMIs) per lineage cassette integration barcode (intBC) detected in fully-edited cell clones by droplet-based single-cell RNA-seq (scRNA-seq). (**B**) Mapping of 30nt sequencing intBCs to 183nt imaging-decoded intBCs by scRNA-seq. (**C**) Wide field of view (top) and higher magnification view (bottom) for experiments using T7 *in situ* transcription and imaging following the Askary *et al.* published Zombie protocol (*54*). Cells are colored by their clone identities where clone 1 = red, clone 2 = yellow, clone 3 = green, and clone 4 = blue with undetermined cells (Undet.) that could not be confidently assigned to a clone shown in grey. DAPI nuclear staining (blue) and the two common bits (C1 = green; C2 = magenta). Integration amplicons appear as colocalized common-bit puncta (white). (**D**) Confusion matrix for intBC decoding using the previously-published Zombie *in situ* T7 protocol. (**E**) Decoding efficiency as a function of spot intensity for *in situ* T7 polymerase-generated spots representing integrated lineage cassettes using our new protocol (blue line) versus Zombie protocol (red line). (**F**) Imaging-based readout accuracy of lineage marks (LMs) using the maximum (Max.) spot intensity, the logistic regression (LR) classifier, or the LR classifier with a p >0.7 cutoff with 5-fold cross validation. (**G**) Logistic regression classifier decoding workflow from images to LM assignment probabilities for an example spot and edit site. (**H**) Confusion matrix depicting the decoding accuracy with 5-fold cross validation for each individual LM and the unedited (UE) state using the PEtracer *in situ* T7 amplification protocol. For each intBC in each cell, the “actual” LM from the ground-truth intBC-LM mapping is compared to the decoded LM from the processing workflow in (G).

**Fig. S6.**
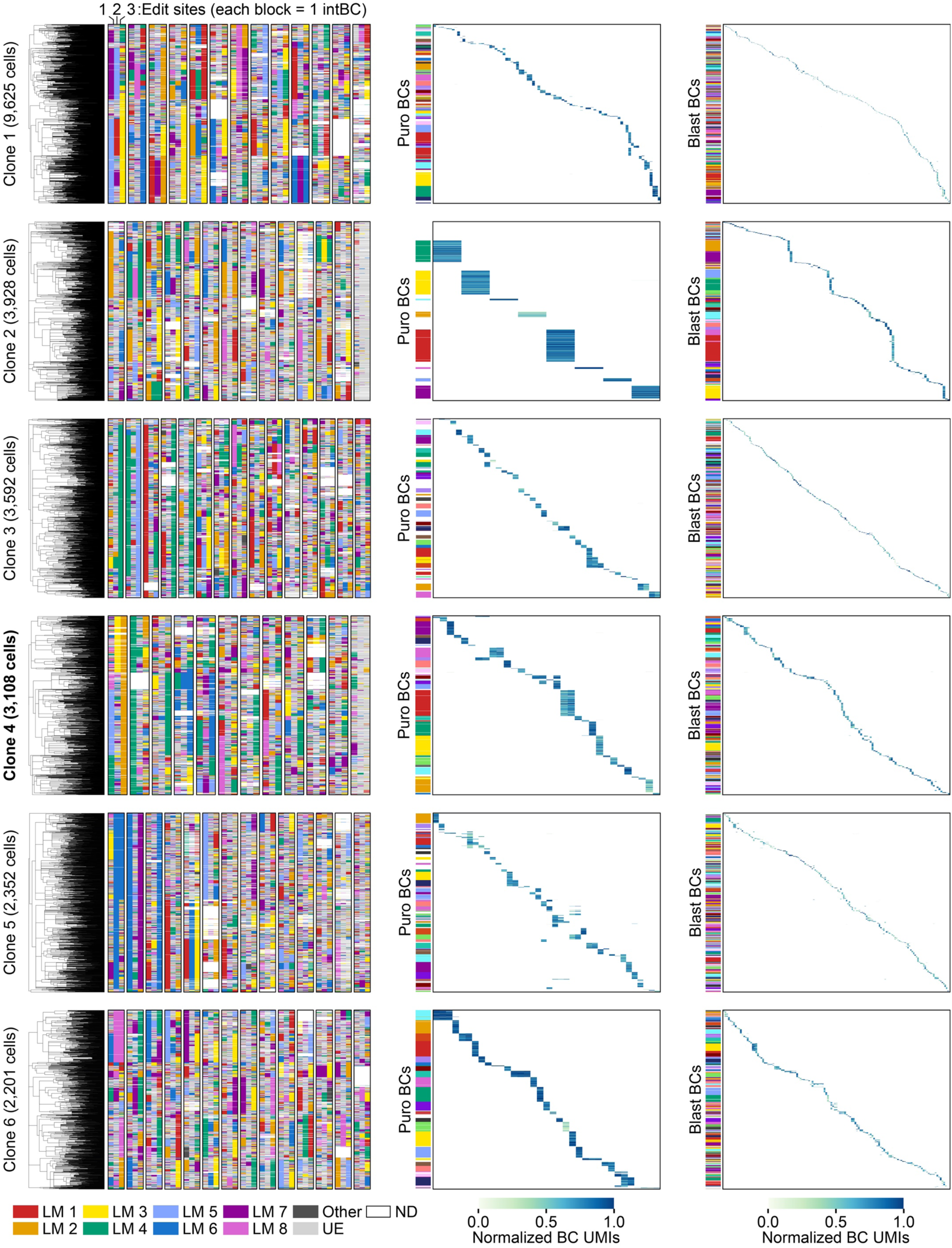
Phylogenetic reconstructions with paired puromycin and blasticidin barcode data. Character matrices and phylogenies for the 6 clones assayed in the single cell RNA sequencing *in vitro* barcoding experiment reconstructed with the neighbor joining algorithm and PEtracer evolving lineage marks (LMs) along with heatmaps of normalized unique molecular identifier (UMI) counts for Puromycin (Puro) and Blasticidin (Blast) static barcodes (BCs). Clone 4 is bolded as it is highlighted in the main text. Imperfect static BC groups indicated as gaps in called groupings. Notably, imperfect calling of barcode groups was partially responsible for FMI scores <1. Integration barcodes are denoted as black rectangles comprising three associated edit sites colored by LM identity. Color key for LM identities provided in bottom left; UE = unedited, ND = not detected. Puro and Blast BC groups based on normalized BC UMI counts denoted as colored blocks to the Puro BC and Blast BC heatmaps.

**Fig. S7.**
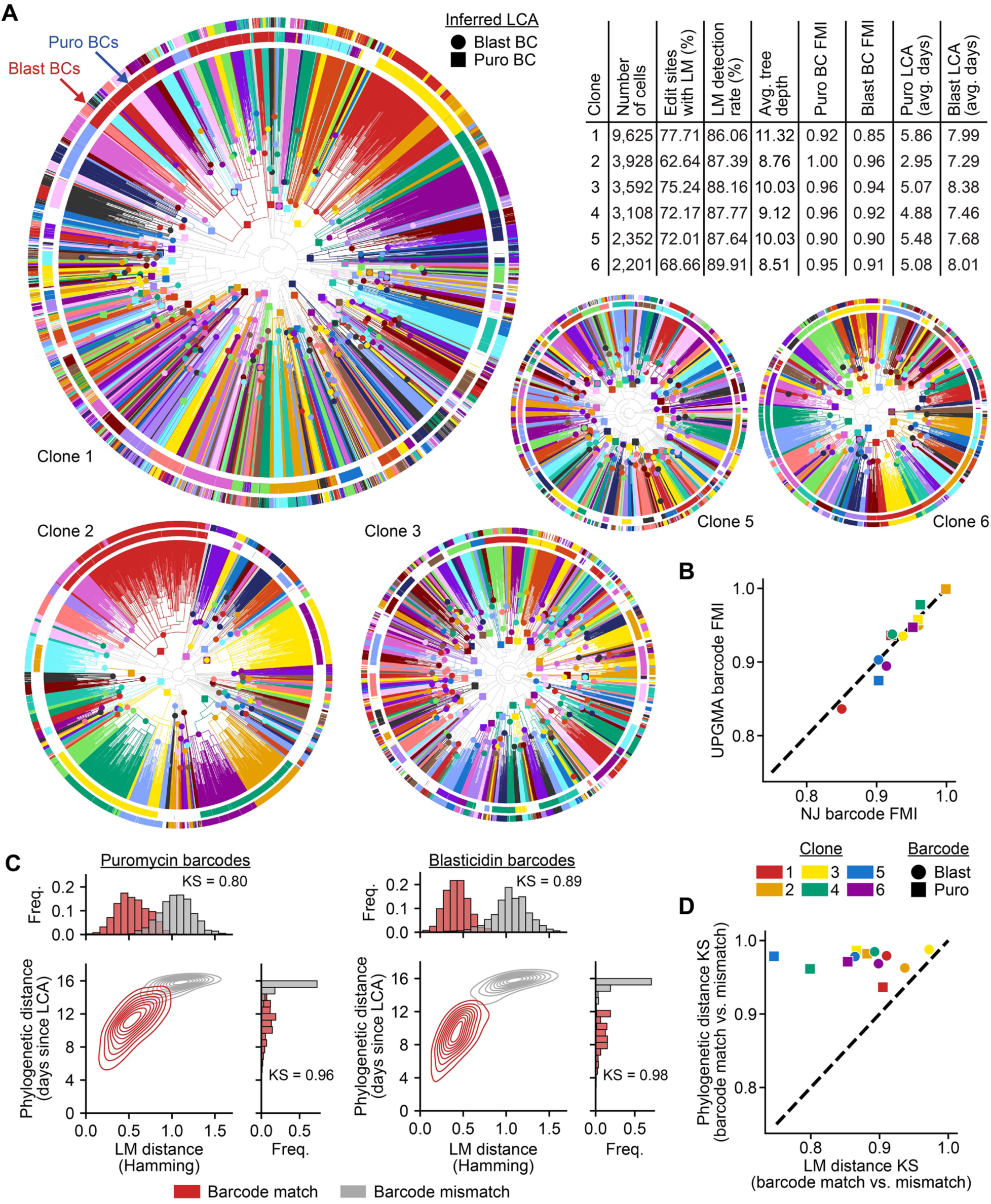
Characterization of phylogenetic reconstruction performance with scRNA-seq PEtracer data. (**A**) Reconstructed phylogenies for clones 1, 2, 3, 5, and 6. Coloring and plotting identical to Fig. 4D where rings are colored to show puromycin (Puro; inner ring) and blasticidin (Blast; outer ring) static barcode (BC) groups for each cell (white indicates missing data). Phylogeny branches colored to match BC assignments. The inferred lowest common ancestor (LCA) is marked and colored to match each BC group (Puro = square; Blast = circle). Embedded table (top right) shows useful statistics for each phylogeny. Average = avg. (**B**) Fowlkes-Mallows Index (FMI) scores measuring the agreement between LCA clades and both puromycin and blasticidin static BC groups when trees are reconstructed with the UPGMA and Neighbor Joining (NJ) algorithms. (**C**) Pairwise LM distance versus phylogenetic distance for cells sharing (red; barcode match) or not sharing (barcode mismatch; grey) puromycin (left) and blasticidin (right) static barcodes in Clone 4 (main text phylogeny). Kolmogorov-Smirnov (KS) statistic for the separation of matched and unmatched barcode distributions reported for each BC and distance metric. Freq. = Frequency. (**D**) Pairwise LM versus phylogenetic distance KS statistics for puromycin (square) and blasticidin (circle) BC groups across all 6 clones tested in this experiment.

**Fig. S8.**
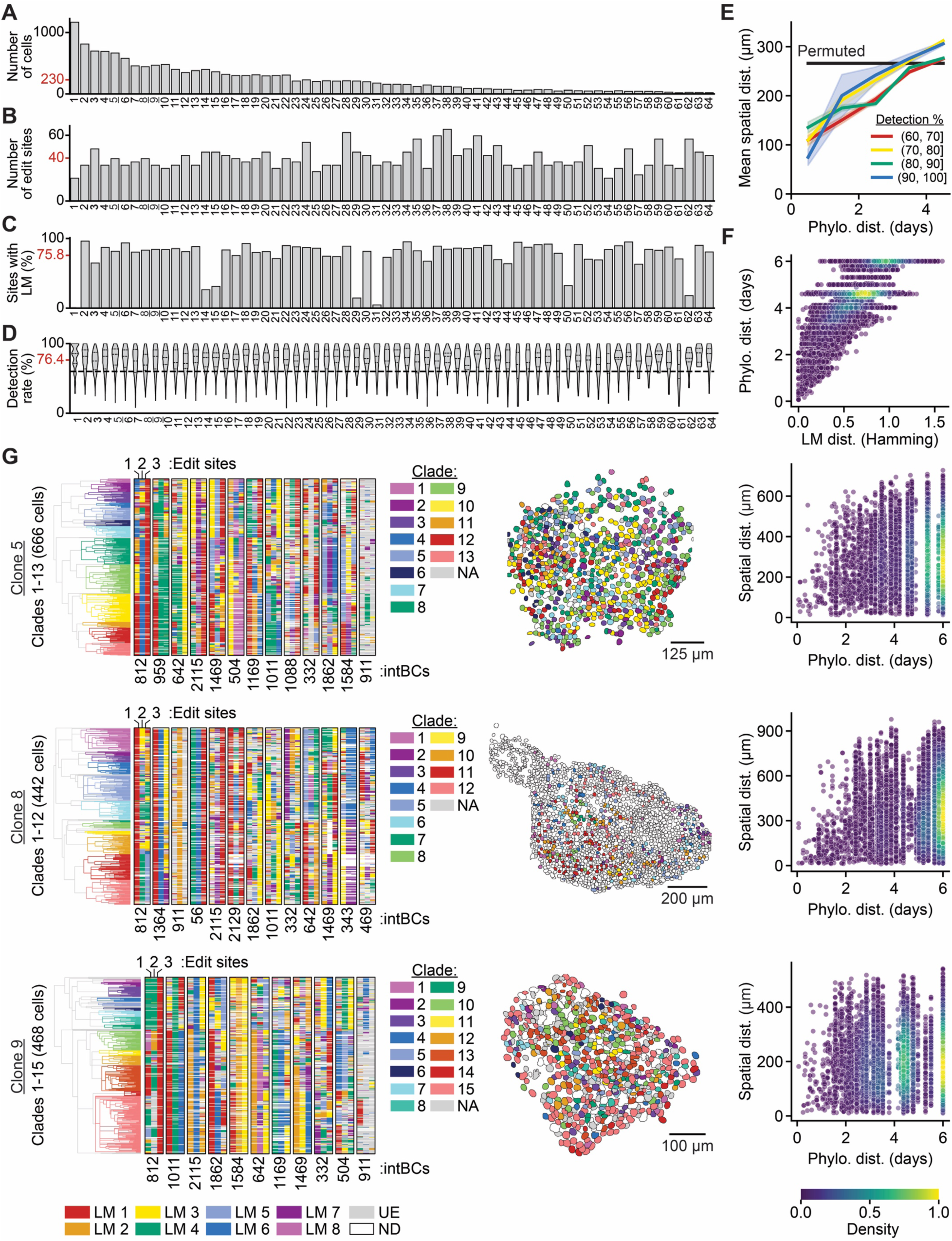
Evaluation of phylogenetic reconstruction performance with imaging-based PEtracer readout. The number of cells (**A**), number of edit sites (**B**), % of sites with a lineage mark (LM) modification (**C**), and the integration barcode (intBC) detection rate (**D**) for 64 evolving clones of 4T1 cells imaged on a single cover slip. Average value for each plot marked in red on the y-axis. Dashed lines on violin in (D) indicate median and dotted lines indicate 25^th^ and 75^th^ percentiles. Clone 3 is bolded as it is shown in the main text, Clones 5, 8, and 9 are underlined as they are depicted throughout this figure. Cells with detection rates <60% (indicated with dashed line in D) were excluded from downstream analyses. (**E**) For clone 3, mean pairwise phylogenetic versus spatial distance for cells grouped by detection rate decile. Ribbons show standard error and black line is the mean distance between randomly permuted pairs. (**F**) For clone 3, pairwise phylogenetic versus LM distance. (**G**) Data for clones 5, 8, and 9. Left shows the character matrix and phylogeny for each clone. intBCs are listed below each character matrix where the three edit sites are colored by LM identity; UE = unedited, ND = not detected. The number of cells in each clone is denoted and the phylogeny is colored by clade. Clades are listed for each clone and are used to pseudo-color nuclear masks for the colony shown in the middle. NA = not assigned. Grey cells = not assigned a clade; White cells = not included in phylogeny. Pairwise phylogenetic versus spatial distance for each clone provided on the right. Phylo. = phylogenetic and dist. = distance.

**Fig. S9.**
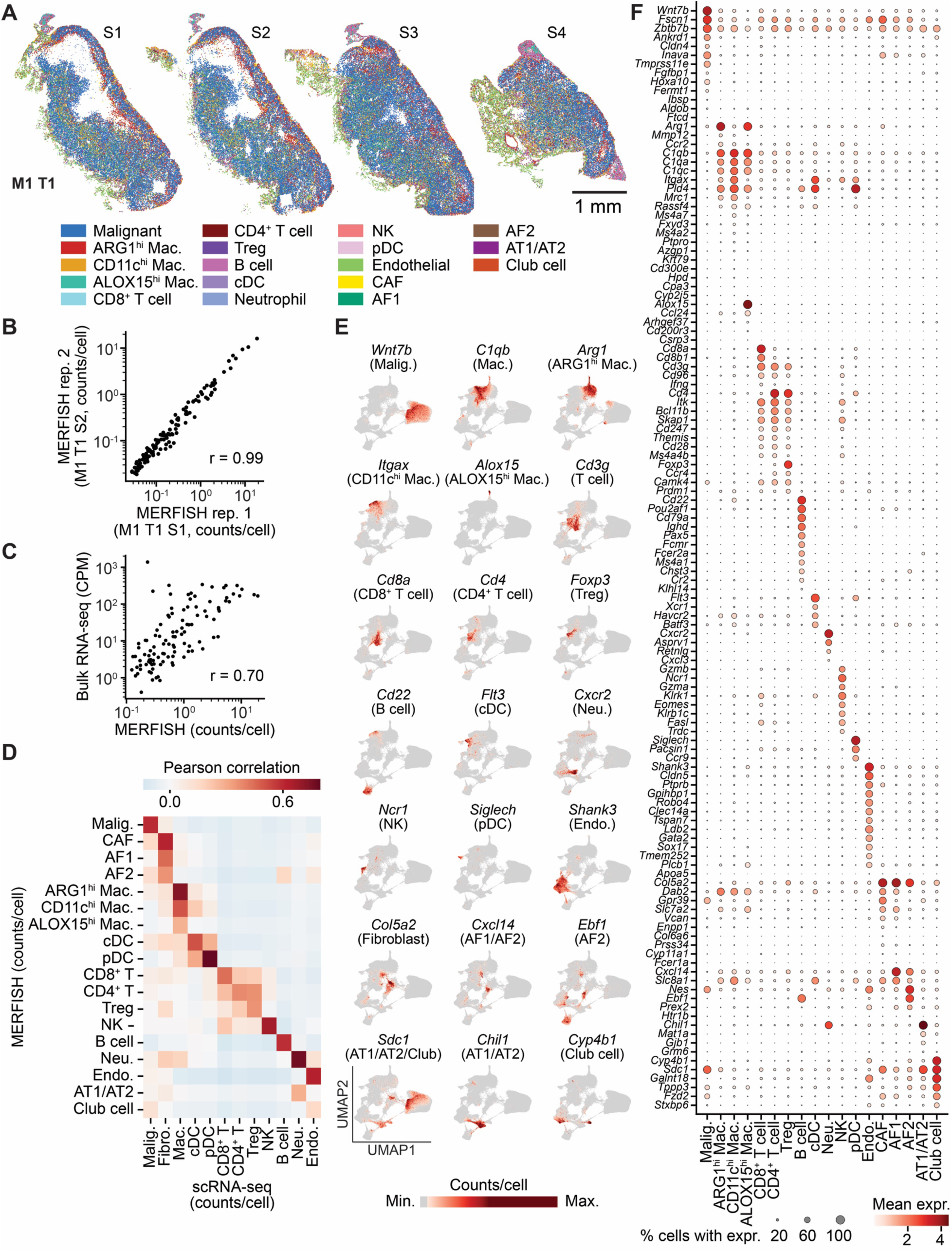
Reproducibility and additional markers for MERFISH transcriptomic data. (**A**) Spatial organization of cell types across four sections (S1-S4) of Mouse 1 Tumor 1 (M1 T1). Cell types colored to match Fig. 5. Scale bar = 1 mm. (**B**) Replicate correlation of transcript counts per cell of individual genes detected in MERFISH data acquired from M1 T1 S1 and M1 T1 S2. (**C**) Correlation between MERFISH counts per cell of individual genes across M1 T1 and bulk RNA-seq data for 4T1 lung metastases from Ferrer *et al* (*114*); r = Pearson correlation. (**D**) Pairwise Pearson correlation of mean counts per cell from our M1 T1 MERFISH dataset with mean counts per cell in scRNA-seq data generated from 4T1 primary tumors as well as lung and liver of tumor-bearing mice. (**E**) UMAP embedding of all 368,722 cells from Mouse 1 and Mouse 2 tumors colored by marker gene expression level for each of the 18 distinct cell types annotated with this library. (**F**) Gene expression (expr.) profiles for all 124 probed genes across the 18 cell types resolved with this library. Dot color indicates the mean gene expression for a gene within a cell type while dot size indicates the percent of cells within a cell type that express that gene. Cell type abbreviations are as follows: Malig. = malignant, Mac. = macrophage, Treg = regulatory T cell, NK = Natural Killer cell, cDC = Conventional Dendritic Cell, pDC = plasmacytoid Dendritic Cell, Neu. = Neutrophil, Endo. = endothelial, CAF = Cancer Associated Fibroblast, AF1/AF2 = Alveolar Fibroblast Type 1/2, AT1/AT2 = alveolar epithelial type 1/2.

**Fig. S10.**
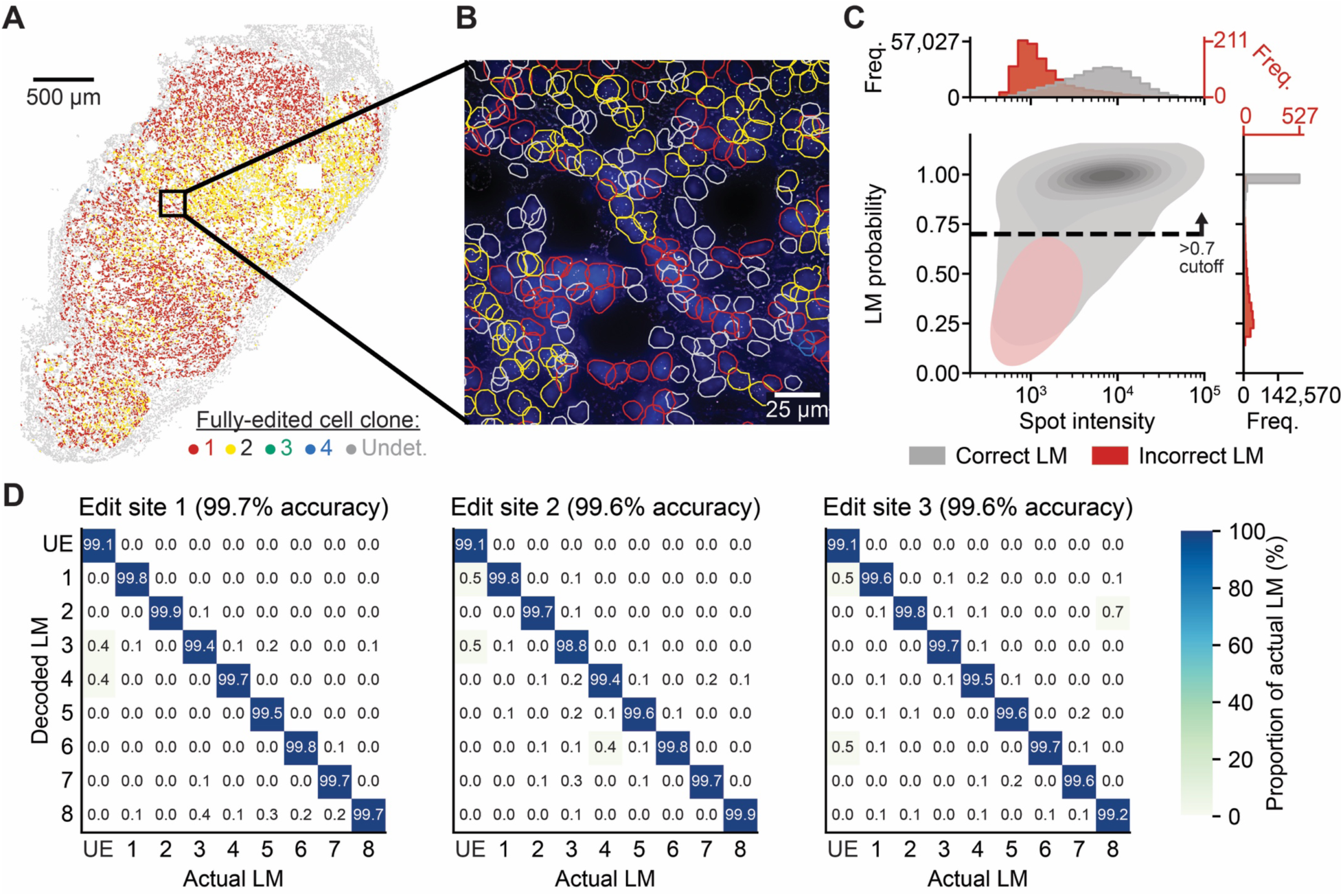
*In vivo* lineage mark detection using fully-edited 4T1 cells. (**A**) Spatial organization of fully-edited 4T1 cells seeded in the mammary fat pad of a female mouse. Nuclei are colored by their clone assignments, matching Fig. 3. Clone 1 = red, clone 2 = yellow, clone 3 = green, and clone 4 = blue and grey = undetermined (Undet.) and non-malignant cells which do not contain the PEtracer components. Scale bar = 500 µm. (**B**) Higher magnification view of a field of view in this tumor with DAPI nuclear staining (blue) and nuclei outlines colored by clone assignment. Scale bar = 25 µm. (**C**) Validation and performance of the logistic regression classifier for LM assignment from *in vivo* imaging-based PEtracer data. Assignment probability cutoff of p >0.7 for decoded LMs is noted as a dashed line. LM = lineage mark. Freq. = frequency. (**D**) Confusion matrix depicting the decoding accuracy for each individual LM and unedited (UE) state *in vivo*. For each intBC in each cell, the “actual” LM from the ground-truth intBC-LM mapping is compared to the decoded LM.

**Fig. S11.**
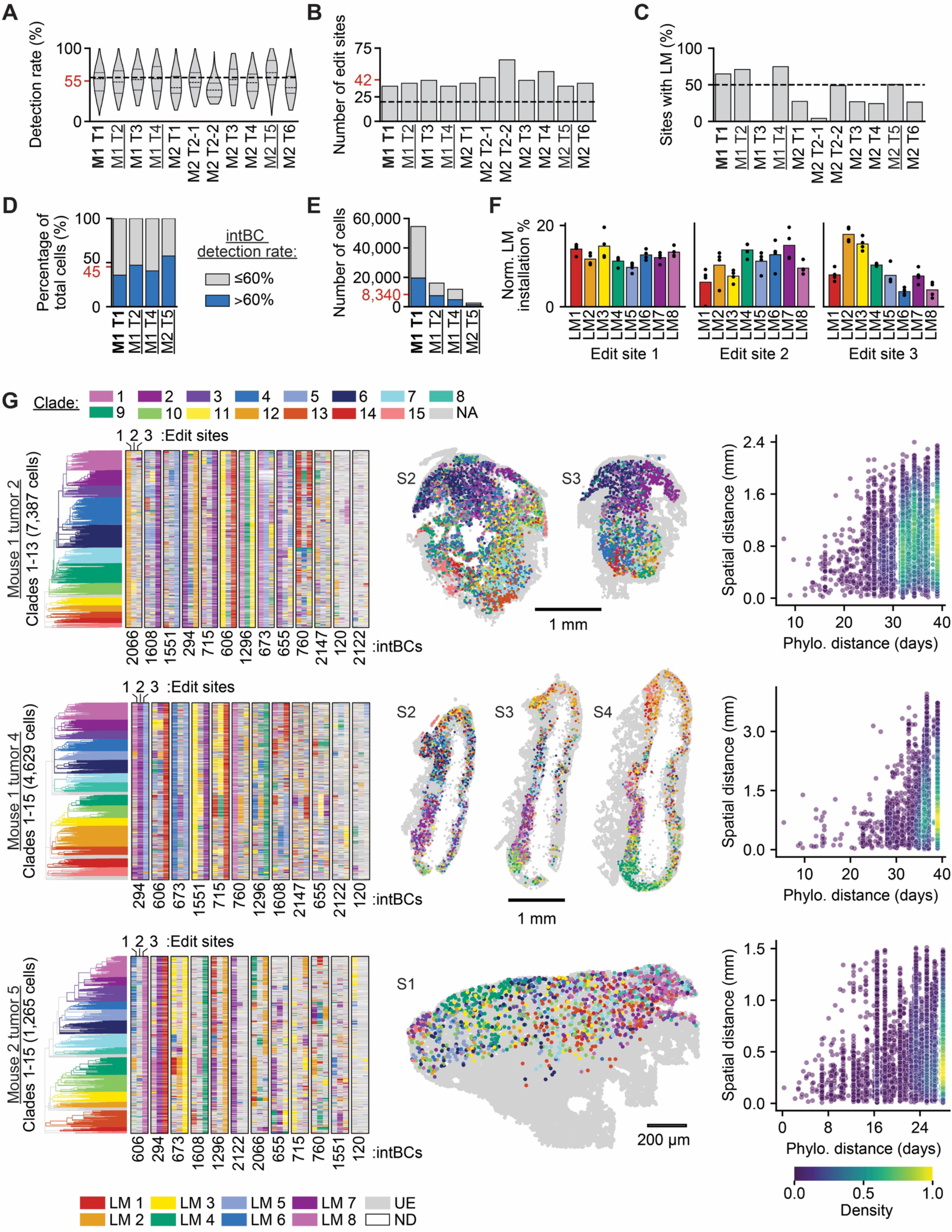
*In vivo* phylogenetic reconstructions with high-resolution MERFISH-based imaging data. The detection rate (**A**), number of edit sites (**B**), and fraction of sites with lineage mark (LM) modification (**C**) for all tumors across Mouse 1 and Mouse 2. Average value for each plot marked in red on the y-axis where relevant. Tumor clones with mean editing fractions <50% (indicated with dashed line in C) were excluded from downstream analyses. The percent of total malignant cells (**D**) and count of malignant cells (**E**) that satisfy our filtering criteria and are therefore included or excluded from phylogenies generated for the four tumors we further analyzed in this study. Dashed lines on violin in (A) indicate median and dotted lines indicated 25^th^ and 75^th^ percentiles. The M1 T1 phylogeny is shown in Fig. 6, while phylogenies for M1 T2, M1 T4, and M2 T5 are detailed below. (**F**) Normalized LM installation efficiency for all eight LMs at each of the three edit sites based on LM transition probability along branches. Mean of four analyzed tumors depicted. (**G**) Left shows the character matrix and phylogeny for the cells with >60% integration barcode (intBC) detection for each tumor. intBCs are listed below each character matrix where each edit site is colored by LM identity; UE = unedited, ND = not detected. LM color assignment is listed at the bottom and is consistent with (F). The number of cells in each clone is denoted and the phylogeny is colored by clade. Clades are listed and colored at the top left. NA = not assigned. All cells in the phylogeny across multiple sections (S#) are pseudo-colored by their clade assignment in the middle. Some tumors were not in all sections. Pairwise phylogenetic (Phylo.) versus spatial distance for each phylogeny is provided on the right. Scale bars provided for each tumor.

**Fig. S12.**
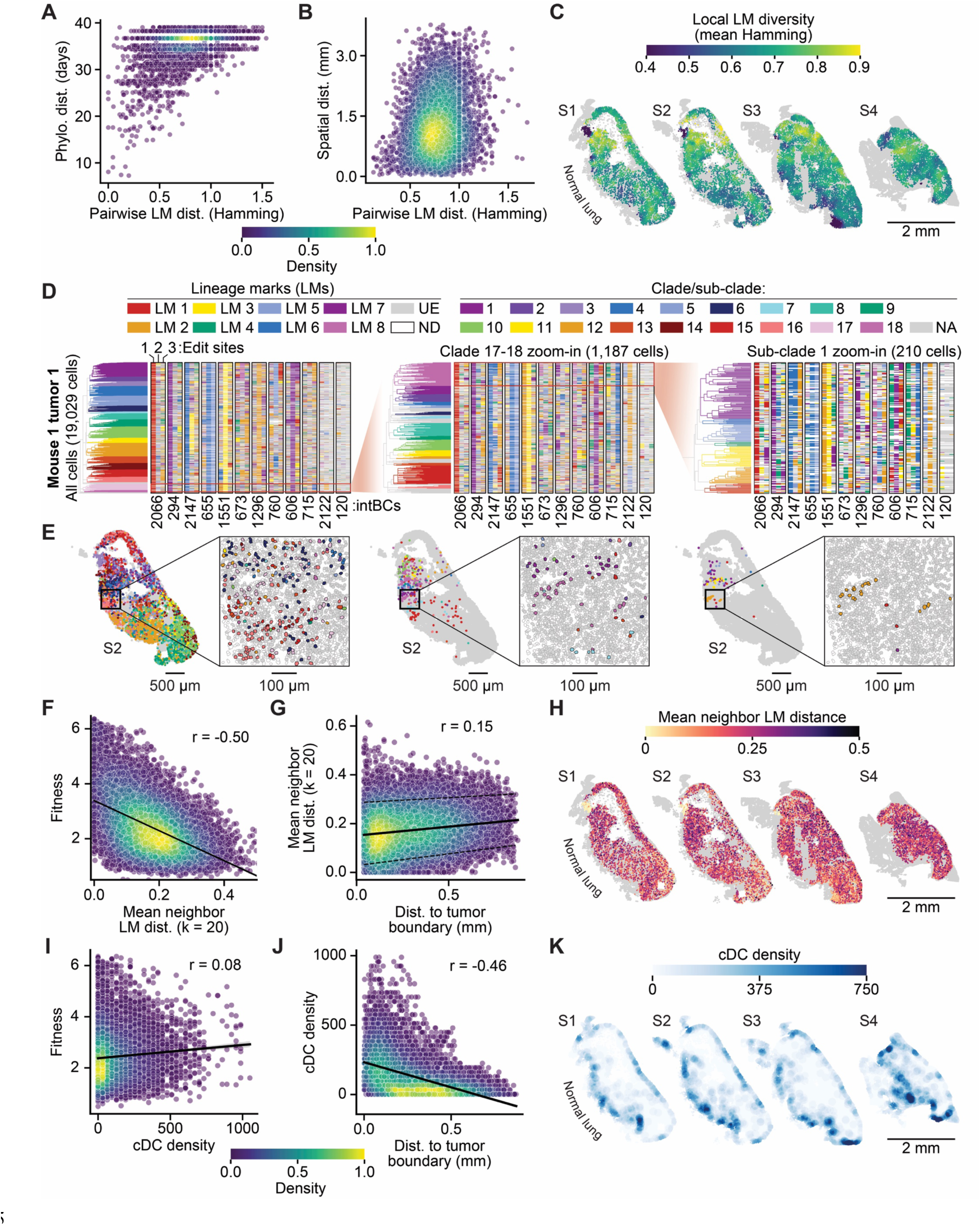
Phylodynamic analyses of tumor evolution *in vivo*. (**A**) Pairwise LM versus phylogenetic (phylo.) distance (dist.) for cells with >60% integration barcode (intBC) detection rate in Mouse 1 Tumor 1 (M1 T1). (**B**) Pairwise LM versus spatial distance for cells in (A). (**C**) Spatial organization of the local LM diversity, defined as mean pairwise Hamming distance within a 100 µm radius circle for all 19,029 cells in the M1 T1 phylogeny across four sections (S1 to S4) (**D**) Character matrix and phylogeny zoom-in for cells in the M1 T1 phylogeny across four sections (S1 to S4). intBCs are listed below the character matrix, which is colored by LM identity for each edit site; UE = unedited, ND = not detected. Clades 1 through 18 are colored on the phylogeny on the left with a key at top right; NA = not assigned. Zoom-ins of smaller portions of the tree proceed from left to right. These zoom-in phylogenies are recolored by sub-clade to aid in visual inspection of the spatial distribution of assigned sub-clades for each sub-tree. (**E**) Spatial positions in section S2 of cells in (D) colored by their clade or sub-clade assignment (same color scheme as (D)). Scale bar = 500 µm for full section; scale bar for inset = 100 µm. Higher magnification inset shows nuclear segmentation masks. Grey circles denote malignant cells not assigned to a clade or not included in the sub-trees; white cells denote non-malignant cells. (**F**) Mean LM distance (Hamming) to 20 nearest neighbors in the character matrix versus fitness. Black line depicts regression with ribbon indicating 95% confidence interval for (F, G, I, J). (**G**) Dist. to the tumor boundary versus mean neighbor LM distance. Dashed lines indicate top and bottom decile. (**H**) Spatial organization of mean LM distance for all cells in the M1 T1 phylogeny across four sections (S1 to S4). (**I**) Conventional dendritic cell (cDC) density versus malignant cell fitness. (**J**) Distance (dist.) to the tumor boundary versus DC density. (**K**) Spatial positions of cells colored by local cDC density across M1 T1 tumor four sections (S1 to S4).

**Fig. S13.**
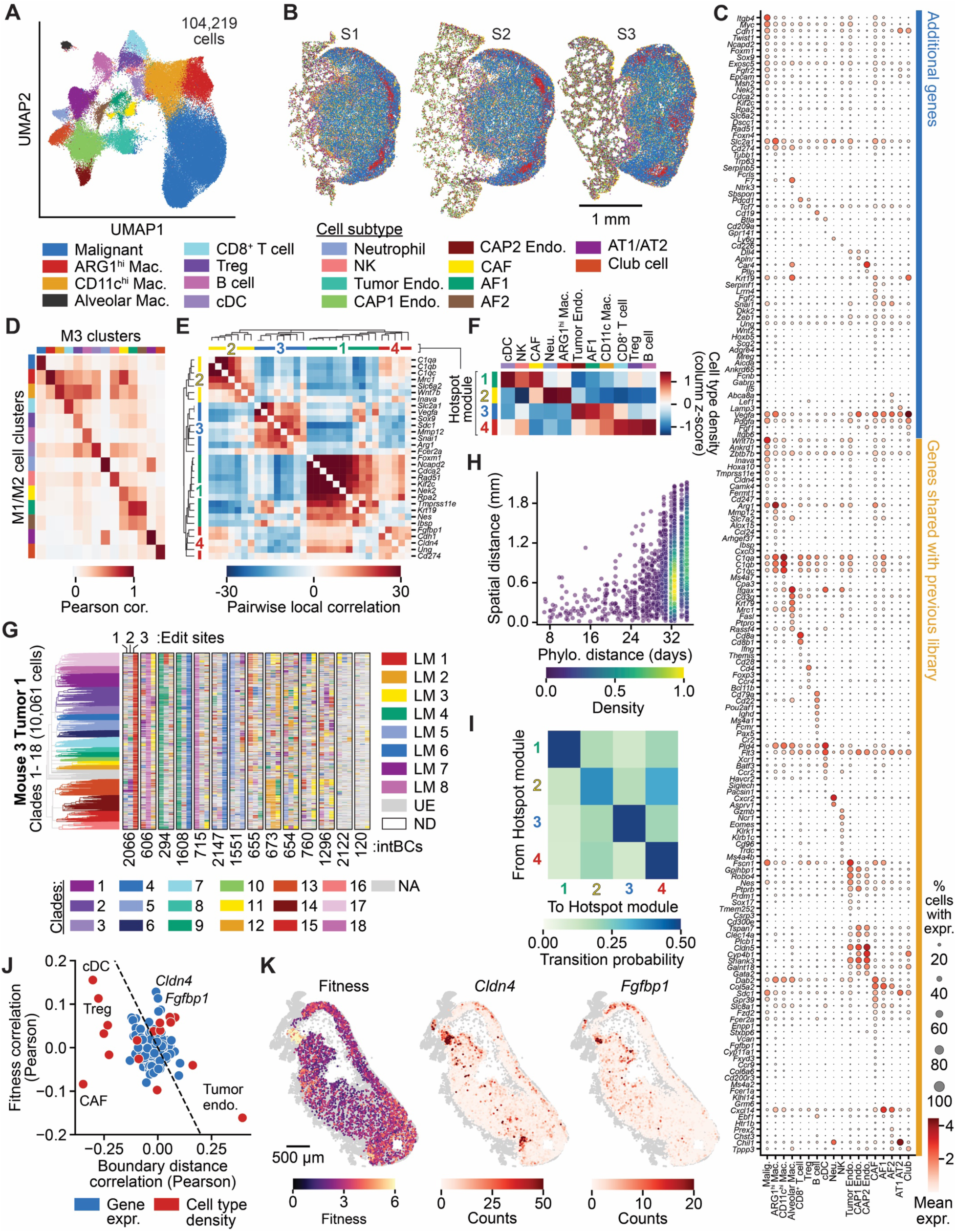
Consistent features driving metastatic 4T1 lung lesion growth *in vivo*. (**A**) UMAP embedding of 104,219 cells from Mouse 3 Tumor 1 (M3 T1) Sections 1-3 (S1-S3) where each cell is colored by its assignment to 18 cell types annotated with this library. Cell types and colors are listed below the UMAP. (**B**) Spatial organization of cell types in M3 T1 S1-S3. Cell types colored using the annotations in (A). Scale bar = 1 mm. (**C**) Gene expression profiles for all 175 probed genes across the 18 cell types annotated with this library. Dot color indicates the mean gene expression for a marker gene within a cell type, while dot size indicates the percent of cells within a cell type that express that gene. (**D**) Pearson correlation between average gene expression of shared genes in the 124-gene MERFISH library for M1/M2 versus the 175-gene library for M3 for each cell type resolved by both libraries. (**E**) Pairwise local correlation of gene expression MERFISH counts between Hotspot modules. (**F**) Heatmap of average cell type densities by Hotspot module. (**G**) Character matrix and phylogeny for the 10,061 cells with >60% intBC detection efficiency across S1 to S3 for M3 T1. Integration barcodes (intBCs) are listed below the character matrix, which is colored by lineage mark (LM) identity for each edit site; LM key shown to the right of the character matrix; UE = unedited, ND = not detected. Clades 1 through 18 are colored on the phylogeny with a key at the bottom. NA = not assigned. (**H**) Transition probabilities from one Hotspot module to another, quantified as the fraction of phylogenetic relatives (lowest common ancestor within 10 days) for each module. (**I**) Phylogenetic (Phylo.) versus spatial distance for reconstructed malignant cells in M3 T1. (**J**) Pearson correlation with malignant cell boundary distance versus Pearson correlation with malignant cell fitness for cell type densities and gene expression (expr.) signatures. Dashed line depicts y = -x as a guide. (**K**) Spatial organization of malignant cell fitness (left), *Cldn4* MERFISH counts (middle), and *Fgfbp1* MERFISH counts in M3 T1. S3 shown as a representative section. Scale bar = 500 µm. Cell type abbreviations are as follows: Malig. = Malignant, Mac. = macrophage, Treg = regulatory T cell, cDC = conventional Dendritic Cell, Neu. = Neutrophil, NK = Natural Killer cell, Endo. = endothelial, CAF = Cancer Associated Fibroblast, AF1/AF2 = Alveolar Fibroblast Type 1/2, AT1/AT2 = alveolar epithelial type 1/2, CAP1/CAP2 Endo. = Capillary type 1/2 Endothelial. See table S24 for relevant statistics.

**Fig. S14.**
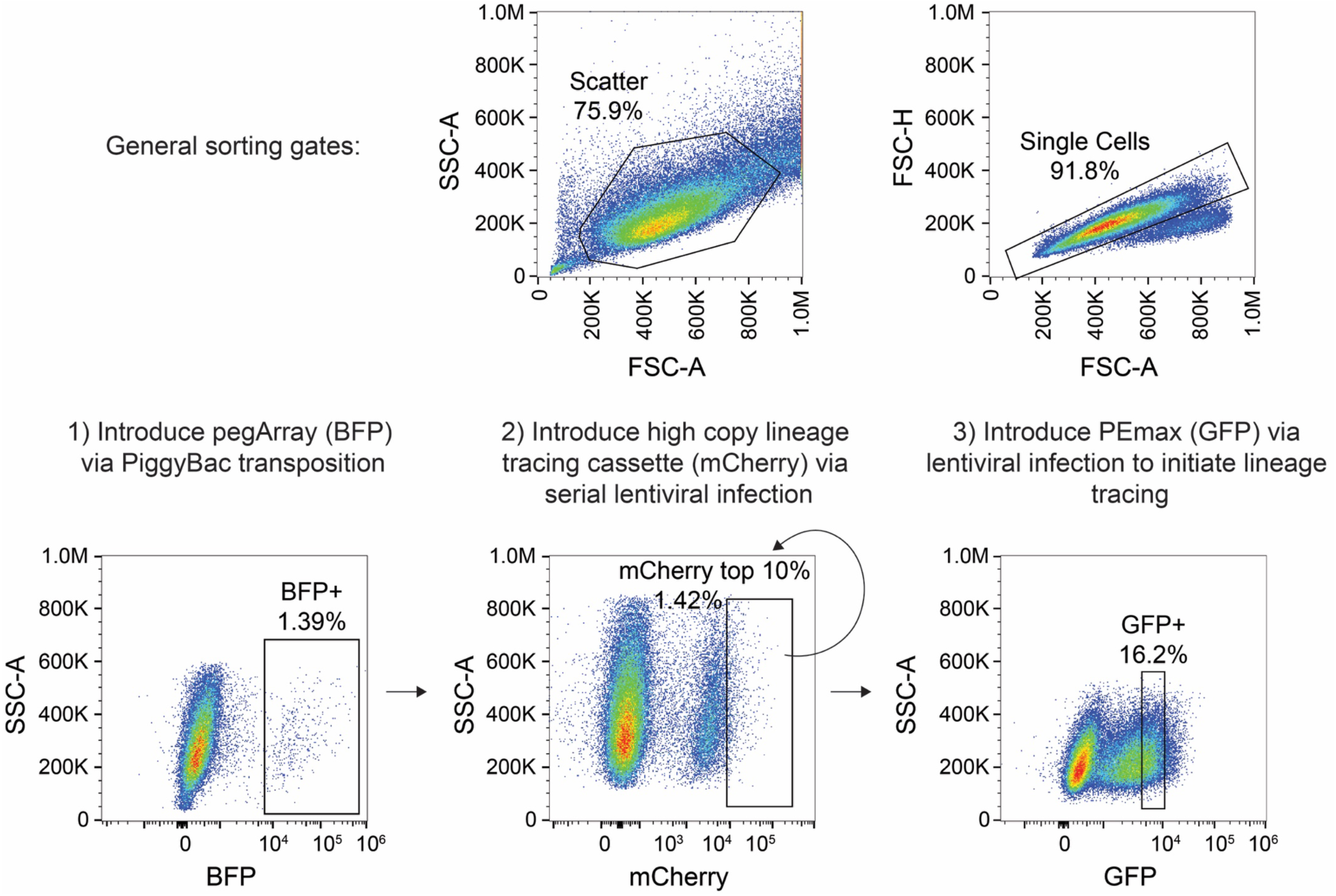
Representative flow cytometry sorting gates for cell line engineering with PEtracer components. Cells were sorted based on expression of the indicated fluorescent markers, with percentage of cells from the parent gate indicated. For generation of cells with high numbers of integrated tracing cassettes, cells were serially infected with lineage tracing cassette lentiviral libraries and the brightest 10% of mCherry expressing cells were sorted to isolate cells with high integration numbers. This process was repeated to increase integration count until the average number of integrations in the population was over 10 as evaluated by qPCR.

## Materials and Methods

### *In silico* lineage tracing simulation

Lineage trees and associated tracing data were simulated using the Cassiopeia framework (v2.0.0) (*86*). A birth-death process was employed to generate lineage trees, where internal nodes represented cell divisions, branch lengths corresponded to individual cell lifetimes, and leaves represented the cells observed at the experiment’s conclusion. The time between cell divisions was modeled using a log-normal distribution (mean = 1, sigma = 0.5) and the cell death rate was set at 0.25. Each simulation was run until the desired population size was reached or all cells died. These simulated trees were subsequently used to generate tracing data under varying conditions.

To evaluate how key experimental parameters influence the accuracy of reconstructed trees, tracing data was simulated while varying the following parameters:

- *N*: The number of extant cells at the end of the experiment
- *M*: The number of edit sites available for recording information
- *S*: The number of unique lineage marks (LMs) that could be installed at each edit site
- *X*: The distribution of LM installation probabilities
- *F*: The fraction of edit sites with an LM installed at the end of the experiment
- *D*: The efficiency of LM detection at each edit site

Simulations began at the tree’s root with *M* unedited sites. Edits occurred stochastically and were inherited according to the tree structure. To achieve the target edit fraction *F* over a total experimental time *T*, edits occurred at a rate *R* given by:

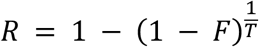

The LM installed during each edit was randomly selected based on the distribution *X.* At the end of the simulation, a fraction *1 – *D** of the LMs was randomly dropped out to account for imperfect detection.

### Distance-based tree reconstruction

For tree reconstruction with neighbor joining and UPGMA a *N*-by-*N* weighted Hamming distance matrix was computed. The distance between two cells *i* and *j* is defined as:

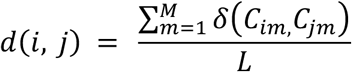

Here, *C* represents the *N* cell by *M* edit site matrix of detected LMs, *L* is the number of edit sites where both cells have an LM detected, and the distance for edit site *m* is defined as:

δ(*c_i_, c_j_)* = 0 if *c_i_* matches *c_j_* or either is missing

- if *c_i_* or *c_j_* is unedited
- otherwise (both edited and not equal)

To efficiently reconstruct large trees, optimized versions of neighbor joining and UPGMA were used (*92*). These algorithms employ dynamic programming to achieve an *O(n^2^)* runtime and are implemented in the CCPhylo software suite (v0.8.3, https://github.com/genomicepidemiology/ccphylo).

For rooting trees generated by neighbor joining, a synthetic cell with all sites unedited was added. The tree was then rooted using this synthetic cell as the outgroup, after which the synthetic cell was removed.

### Greedy tree reconstruction

We also evaluated tree reconstruction using the “vanilla” greedy algorithm from Cassiopeia (*86*). This algorithm builds a tree from the top down by recursively selecting lineage marks (LMs) that split the cells into progressively smaller groups. LMs are chosen based on their frequency within the group, with lower-frequency LMs favored because they are less likely to exhibit homoplasy (i.e., multiple independent occurrences of the same edit at the same edit site in separate cells). Although the greedy algorithm performed well for phylogenies labeled with the Cas9 indel LM distribution, it underperformed in distance-based tree reconstruction with only eight LMs due to increased homoplasy. Consequently, only the neighbor joining and UPGMA methods were used to reconstruct phylogenies generated with PEtracer LM data. Hybrid approaches, which pair a greedy algorithm for solving the top of the tree with a more accurate method, such as integer linear programming, for solving the bottom of the tree, can perform well but were not evaluated here since they require high-confidence greedy splits (*86*).

### Evaluation of simulated tree reconstruction

Two complementary metrics were used to evaluate the similarity between simulated ground-truth trees and reconstructed trees with UPGMA, neighbor joining, and Greedy tree reconstruction algorithms. The first metric was Robinson-Foulds (RF) distance, which quantifies topological differences by counting the number of node partitions unique to each tree (*87*). The RF metric was normalized by the total number of partitions, resulting in a range between 0 and 1. As a distance metric, lower values indicate greater similarity.

The second metric was depth-normalized mean triplets correct, which compares trees based on the relative ordering of leaf triplets rather than the set of partitions. To compute this metric, 1,000 triplets were sampled from the ground-truth tree, distributed evenly across lowest common ancestor (LCA) depths to minimize bias introduced by most triplets having an LCA near the tree’s root. The fraction of triplets with conserved ordering in the reconstructed tree was then calculated with values that range from 0 to 1. As a similarity metric, higher values indicate greater similarity.

All simulations were repeated 10 times, with reconstruction accuracies and parameters for each iteration reported **tables S1-3.** The mean value for each metric is presented with ribbons indicating standard deviation where applicable. In each iteration, the tree topology was kept constant while the simulated tracing data varied based on the specified parameters, except when assessing the effect of tree size on reconstruction accuracy. For values presented in heatmaps, the performance of the best algorithm for the given parameter combination is reported.

### LM distributions

The uniformity of LM installation probabilities was quantified using normalized entropy (*H_norm_*), calculated with the following formula:

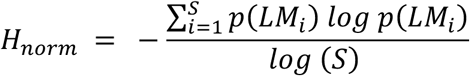

Here, *p*(*LM_i_*) represents the probability of the *i*-th LM being installed, and *S* is the total number of lineage marks.

In simulations, the LM distributions with a given *H_norm_* were generated by adjusting the λ parameter of an exponential distribution until the target *H_norm_* was achieved. Thus, the probability of the *i*-th LM is given by:

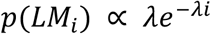

with probabilities normalized to sum to 1. The CRISPR-Cas9 indel distribution used in these simulations was empirically derived from data published by Yang, Jones, et al. 2022 (*82*).

### Edit rate simulations

A simplified simulation framework was used to model the relationship between edit rate and edit saturation over time. In this framework, it was assumed that all cells proliferate uniformly at a rate of one division per day, with no cell death. Instead of modeling all cells, which is computationally expensive because the number of cells grows exponentially, we model 100 cells over the duration of the simulation. Each parameter combination was simulated 10 times, and the mean values were reported.

To determine the edit rates required to achieve 50%, 60%, 70%, 80%, and 90% saturation, simulations were performed for edit rates ranging from 0 to 0.3 edits/day in increments of 0.001. For each experiment length, spanning 1 to 100 days, the edit rate closest to achieving the desired saturation value was identified.

To identify the minimum number of edit sites required for 70%, 80%, and 90% of cell divisions to be marked by an edit as a function of tree size, we simulated experiments from 1 to 30 days in length–corresponding to tree sizes between 2^1^ and 2^30^ cells. For each tree size, the number of edit sites was varied from 1 to 120 (in increments of 1), and edit rates were tested from 0 to 0.3 (in increments of 0.001). The minimum number of edit sites required to achieve the target edit fraction was determined for each tree size using the optimal edit rate for each experiment length. These values represent a lower bound on the required number of edit sites, as the simulations assume uniform division rates, perfect detection of all cells in the tree, and ideal tracing kinetics. Results from these simulations are reported in **table S4**.

### General mammalian cell culture conditions

4T1 cells (American Type Culture Collection (ATCC) CRL-2539) were maintained in RPMI 1640 medium (Thermo Fisher Scientific) supplemented with 10% FBS and 1% Penicillin-Streptomycin-Glutamine (PSG) (Thermo Fisher Scientific). B16-F10 cells (ATCC CRL-6475) were maintained in Dulbecco’s Modified Eagle’s Medium (DMEM; Thermo Fisher Scientific) supplemented with 10% FBS and 1% PSG. HEK293T (ATCC CRL-3216) and HEK293T/17 cells (ATCC CRL-11268) cells were maintained in DMEM supplemented with 10% FBS and 1% PSG. All cells were maintained at 37°C and 5% CO_2_, authenticated by their suppliers, and tested negative for mycoplasma.

### Mice

All mouse experiments described in this study were approved by the Massachusetts Institute of Technology Institutional Animal Care and Use Committee (IACUC, protocol numbers 0421-043-24 and 2311000598). To mitigate rejection of engineered 4T1 PEtracer cells, which are syngeneic in the Balbc/J background and express immunogenic components including PEmax (spCas9(H840A)–M-MLVRT) and mCherry (linked to lineage tracing cassettes), experiments were performed in F1 Balbc/J x SELECTIV (*147*) (LSL-Aavr-spCas9-mCherry) or Balbc/J x [SELECTIV x CMV-Cre (Aavr-spCas9-mCherry)] hybrids. Balbc/J (JAX #000651), SELECTIV (JAX #037553), and CMV-Cre (JAX #006054) mice were procured from JAX and bred in house. Tumor experiments were performed in female animals between 8 and 12 weeks of age. We performed tail vein lineage tracing experiments in three mice and eleven total tumors to capture biological variability. Statistical tests were not used to determine sample size and investigators were not blinded to experimental groups.

### Cloning of tracing cassette and editor constructs

DNA fragments for cloning were either ordered from Integrated DNA Technologies (IDT) or Twist Bioscience as dsDNA fragments or PCRed using Phusion Green Hot Start II High-Fidelity PCR 2x Master Mix (Thermo Fisher Scientific) with the following cycle conditions: 98°C for 2 minutes, [98°C for 10 seconds, Primer specific Tm for 20 seconds, 72°C for 30 seconds/kb of amplified insert]x34, 72°C for 2 minutes. GC rich sequences like universal chromatin opening elements (UCOEs) were amplified using the KOD Xtreme Hot Start polymerase for 34 cycles following manufacturer’s protocols (Sigma Aldrich, 71975-M). All PCR-amplified products were DpnI treated following manufacturer’s protocols (New England Biolabs, NEB). DpnI-treated fragments were run out on either 1% or 2% agarose gels (VWR) supplemented with 1 μL of ethidium bromide (Thermo Fisher Scientific) per 10 mL of 1x Tris-Acetate-EDTA (TAE) buffer. Amplified DNA was extracted from bands excised from the gel using the QIAquick Gel Extraction Kit (Qiagen). Gibson assembly was used to assemble fragments into completed plasmids according to manufacturer’s instructions. Specifically, we use 5 μL of NEBuilder 2x HiFi DNA Assembly master mix (NEB) combined with 0.05pmol backbone and 0.15pmol of each insert fragment brought to a final volume of 10 μL with ultrapure nuclease free water. Assemblies were incubated for 60 minutes at 50°C prior to transformation into NEB stable competent *E. coli* (C3040, NEB). Transformation included a 15-minute incubation of 50 μL of competent cells and 5 μL of assembly mix prior to a 30 seconds heat shock at 42°C and a subsequent 5-minute incubation on ice before plating on 100 μg/mL carbenicillin-containing agar plates. The Illustra TempliPhi 100 amplification kit (Cytivia) was used to perform rolling circle amplification (RCA) on bacterial colonies prior to Sanger sequencing or Whole Plasmid Sequencing (WPS) (Quintara Biosciences). Plasmid DNA was isolated from colonies with the correct sequence using the PureYield Plasmid Miniprep system (Promega) or with the ZymoPure II plasmid Midiprep system (Zymo Research).

The following tracing cassette, editor constructs, and other plasmids used in this manuscript were assembled using this protocol and are listed here for ease of reference. The sequence features of plasmids are described below. Addgene plasmid numbers are provided in parentheses where appropriate.

- Lentiviral PEmax editor that includes 5’-UCOE-EF1a short promoter (EFS)-PEmax-P2A-GFP-KASH-WPRE (Addgene # pending)
- Lentiviral lineage tracing cassettes that includes 5’-UCOE- EF1a promoter-mCherry-T7 promoter- T3 promoter- PacI digestion site-183 nucleotide (nt) MERFISH decodable integration barcode (intBC)- Forward primer for sequencing assays-30nt sequencing intBC- PacI digestion site- Edit Site 1 and flanking sequences- Edit Site 2 and flanking sequences- Edit Site 3 and flanking sequences- Reverse primer for sequencing assays- BGH polyA-500nt common sequence (Addgene # pending)
- PiggyBac lineage tracing cassettes that includes 5’-UCOE- mCherry- T7 promoter- T3 promoter- PacI digestion site-183nt MERFISH decodable intBC- Forward primer for sequencing assays- 30nt sequencing intBC- PacI digestion site- Edit Site 1 and flanking sequences- Edit Site 2 and flanking sequences- Edit Site 3 and flanking sequences- Reverse primer for sequencing assays- SV40 polyA-500nt common sequence
- Lentiviral puromycin static barcoding construct that includes 5’-UCOE- EF1a promoter- puromycin resistance cassette-10 N random barcode-BGH polyA (Addgene # pending)
- Lentiviral blasticidin static barcoding construct that includes 5’-UCOE- EF1a promoter-blasticidin resistance cassette-10 N random barcode-BGH polyA (Addgene # pending)
- Lentiviral CROP-seq BFP epegRNA expression vector that includes 5’-EF1a promoter- BFP- WPRE- Split 3’ LTR- hU6 promoter- pegRNA-Split 3’ LTR
- Three BsmBI-digestible PiggyBac 8-mer pegArray acceptor vectors and one PaqCI-digestible 24-mer pegArray acceptor vector that include 5’-UCOE- EF1a promoter-BFP- WPRE-SV40 polyA-Digestion Cassette for pegArray insertions (ES1 8-mer vector = #, ES2 8-mer vector = #, ES3 8-mer vector = #, 24-mer vector = Addgene # pending).

### Cloning of individual pegRNAs or epegRNAs

Individual pegRNAs and epegRNAs were cloned using Gibson assembly as described above. Briefly, hU6-pegRNA-GG-acceptor backbone (Addgene, #132777) expressing a guide RNA under the human U6 promoter was digested using BsaI-HFv2 (NEB, R3733L) according to the manufacturer’s protocol. Digested backbone gel extracted as described above from a 1% agarose gel. eBlock fragments were ordered from IDT or Twist Bioscience encoding the relevant pegRNA or epegRNA sequences which were resuspended in ultrapure nuclease-free water. Gibson assemblies, transformations, colony picking, and DNA isolation from colonies were carried out as described above.

### Testing of individual pegRNAs or epegRNAs

Individual pegRNAs or epegRNAs were evaluated with transfections using low passage HEK293T cells plated into tissue culture-treated (TC-treated) 96-well flat bottom dishes (Corning, CLS3799). Cells were maintained in TC-treated 15cm dishes. For transfections, media was removed, cells were rinsed once with sterile 1x PBS (Thermo Fisher Scientific), and trypsinized with 0.25% trypsin (Gibco). Following a 5-minute incubation at 37°C for 5 minutes to ensure complete trypsinization, cells were resuspended in DMEM + 10% FBS to inactivate the trypsin and carefully triturated prior to cell counting using the Countess II automated cell counter (Thermo Fisher Scientific). Cells were diluted to 18,000 cells/mL and 100 μL of diluted cells were plated into 96-well dishes. Cells were transfected 18 to 24 hours after plating when they were ∼60-70% confluent. For each well, transfections were carried out by combining 1) a DNA mixture comprising 200 ng of prime editor expressing plasmid (pCMV-PE2 plasmid = Addgene plasmid #132775; pCMV-PEmax plasmid = Addgene plasmid #174820) and 50 ng of a given pegRNA or epegRNA plasmid resuspended in 5 μL of OptiMEM (Thermo Fisher Scientific) with 2) 0.5 μL of Lipofectamine 2000 (Thermo Fisher Scientific, 11668019) and 4.5 μL of OptiMEM (Thermo Fisher Scientific, 31985062). This mixture of DNA and lipid were mixed and incubated at room temperature for 15 minutes prior to adding dropwise to cells. At 72 hours post-transfection, media was removed, cells were washed with 1x PBS, and genomic DNA (gDNA) was extracted by adding 35 μL of freshly-prepared lysis buffer (10 mM Tris-HCl pH 7.5, 0.05% sodium dodecyl sulfate (SDS, Thermo Fisher Scientific), 25 µg/ml Proteinase K (Thermo Fisher Scientific)) to each well prior to a 1-hour incubation at 37°C. Following this digestion, gDNA was transferred to a 96-well PCR plate (Thermo Fisher Scientific) and incubated on a deep-well PCR machine for 30 minutes at 80°C to inactivate the Proteinase K; we term this isolation protocol the Fast Lysis procedure.

### Sequencing and analysis of individual pegRNA and epegRNA editing outcomes

Genomic DNA isolated from HEK293T cells was amplified with two successive rounds of PCR as done previously (*71*). Briefly, PCR1 was conducted using a pool of forward and reverse primers where each primer was pooled in an equimolar fashion to enable sequencing on an Illumina MiSeq with minimal need for added PhiX. PCR1 was performed using 2 μL of Fast Lysis-isolated genomic DNA from HEK293T cells. Amplifications were carried out using Phusion Green Hot Start II High-Fidelity PCR 2x Master Mix with the following cycle conditions for the *EMX1*, HEK3, and *RNF2* endogenous genomic loci: 98°C for 2 minutes, [98°C for 10 seconds, 65°C for 20 seconds, 72°C for 1 minute]x30, 72°C for 2 minutes. PCR1 amplicons were all checked on 2% agarose (VWR) gels with 1 μL of ethidium bromide (Thermo Fisher Scientific) per 10 mL of 1x Tris-Acetate-EDTA (TAE) buffer. PCR2 amplification was carried out using TruSeq adapter PCR primers (listed in **table S28**) with the following conditions: 98°C for 2 minutes, [98°C for 10 seconds, 61°C for 20 seconds, 72°C for 1 minute]x8, 72°C for 2 minutes. 2 μL of PCR1 were used as template DNA for each PCR2 reaction and 500 nM final primer concentrations were used. Following PCR2, all reactions for a given genomic locus were pooled together, run on a 2% agarose gel, and gel extracted using the QIAquick gel extraction kit (Qiagen). Concentrations of purified libraries were determined using a Qubit double-stranded DNA high sensitivity Kit (Thermo Fisher Scientific) according to the manufacturer’s instructions. Libraries were diluted to 4 nM and sequenced on a MiSeq (Illumina) using an Illumina MiSeq v2 Reagent kit.

Samples were demultiplexed with the MiSeq Reporter (Illumina). CRISPResso2 (v2.2.6) (*148*) was used to analyze demultiplexed reads. For all prime editing yield quantification, CRISPRessoBatch was run with default settings using the unedited allele as the reference amplicon and desired allele as Amplicon1 (-a parameter) and a 10nt quantification window on each side of the cleavage position for a total length of 20nt (-w parameter). Editing yield was calculated as: (number of Amplicon1-aligned reads)/(total reads aligned to all amplicons). For all experiments, indel yields were calculated as: (number of indel-containing reads)/(total reads aligned to all amplicons).

**HEK Site 3 Genomic Locus Forward Primer Pool:** ACACTCTTTCCCTACACGACGCTCTTCCGATCT-[1N to 4N]-ATGTGGGCTGCCTAGAAAGG
**HEK Site 3 Genomic Locus Reverse Primer:** TGGAGTTCAGACGTGTGCTCTTCCGATCTCCCAGCCAAACTTGTCAACC
***RNF2* Genomic Locus Forward Primer Pool:** ACACTCTTTCCCTACACGACGCTCTTCCGATCT-[1N to 4N]-ACGTCTCATATGCCCCTTGG
***RNF2* Genomic Locus Reverse Primer:** TGGAGTTCAGACGTGTGCTCTTCCGATCTACGTAGGAATTTTGGTGGGACA
***EMX1* Genomic Locus Forward Primer Pool:** ACACTCTTTCCCTACACGACGCTCTTCCGATCT-[1N to 4N]-CAGCTCAGCCTGAGTGTTGA
***EMX1* Genomic Locus Reverse Primer:** TGGAGTTCAGACGTGTGCTCTTCCGATCTCTCGTGGGTTTGTGGTTGC

### Cloning of a comprehensive 1,024 5 nucleotide (nt) LM epegRNA libraries

Gibson assemblies between PCR amplified epegRNA backbones and short oligonucleotide libraries ordered from IDT were used to generate libraries representing all possible 5nt LM libraries for each of the three target sequences. First, epegRNA backbones with optimized primer binding sequence (PBS) and reverse transcription template (RTT) lengths were amplified as described in “Cloning of tracing cassette and editor constructs” using Phusion Green Hot Start II High-Fidelity PCR Master Mix with the primers listed below and 34 cycles of amplification. Two independent 5N LM libraries were ordered from IDT for each site, a top and bottom strand library. Libraries were diluted to 1 μM and 1 μL was used for each of 10 parallel Gibson assemblies with

0.05 pmol of amplified backbone for their respective target loci. All 100 μL of Gibson assembly product were cleaned up using PCR cleanup columns (Qiagen) to remove salts prior to electroporation into MegaX cells as previously described. Colonies were scraped into a combined pool, grown out for 8 hours in 100 mL terrific broth (TB) containing 100 μg/mL carbenicillin, and midi-prepped using the ZymoPure II Plasmid Midiprep Kit (Zymo Research) following manufacturer’s procedures.

**Forward primer for all:** CGCGGTTCTATCTAGTTACGCGTTAAAC
***EMX1* reverse primer:** CTCCCATCACATGCACCGAC
**HEK3 reverse primer:** TGATGGCAGAGGAAAGGAAGCC
***RNF2* reverse primer:** CTGAGGTGTTCGTTGCACCG

***EMX1* 5N library top:** GTCGGTGCATGTGATGGGAGNNNNNTTCTTCTGCTCGGACGCGGTTCTATCTAGTTAC G
***EMX1* 5N library bottom:** CGTAACTAGATAGAACCGCGTCCGAGCAGAAGAANNNNNCTCCCATCACATGCACCG AC
**HEK3 5N library top:** CTTCCTTTCCTCTGCCATCANNNNNCGTGCTCAGTCTGCGCGGTTCTATCTAGTTACG
**HEK3 5N library bottom:** CGTAACTAGATAGAACCGCGCAGACTGAGCACGNNNNNTGATGGCAGAGGAAAGGA AG
***RNF2* 5N library top:** CGGTGCAACGAACACCTCAGNNNNNGTAATGACTAAGATGCGCGGTTCTATCTAGTT ACG
***RNF2* 5N library bottom:** CGTAACTAGATAGAACCGCGCATCTTAGTCATTACNNNNNCTGAGGTGTTCGTTGCAC CG

### Testing of 1,024 LM epegRNA libraries

Testing of 1,024 LM epegRNA libraries for three test sites was carried out with transfections using low passage HEK293T cells plated into TC-treated 6-well flat bottom dishes (Corning, 353046). Cell maintenance and splitting via trypsinization were conducted identical to 96-well format transfections. Following cell counting, cells were resuspended to 400,000 cells/mL and 1 mL of cells was added to each well of a 6-well TC-treated plate to which an additional 1 mL of media was added to avoid evaporation. 18 to 24 hours after plating, three wells were transfected for each 1,024 LM library to ensure comprehensive coverage of the entire library and biological consistency. Each library was transfected as before by combining a mixture of DNA with a lipid mixture prior to a 10-minute incubation at room temperature prior to adding dropwise to 6-well plates. For 6-well formats DNA mixtures comprise 6 μg of PE2 plasmid and 1 μg of 1,024 LM library were resuspended in a final volume of 25 μL of OptiMEM while lipid mixtures comprise 15 μL of Lipofectamine 2000 and 10 μL of OptiMEM. At 72 hours post-transfection, media was removed, cells were washed with 1x PBS, and trypsinized as above to isolate cell pellets. Genomic DNA was extracted using the DNeasy blood and tissue kit (Qiagen) following manufacturer’s protocols.

### Sequencing and analysis of 1,024 LM epegRNA libraries

Plasmid DNA sequences were verified to ensure coverage of all edits 1,024 LMs across the library for each of the three loci. Two successive rounds of PCR were used to generate plasmid libraries using primers listed below. For plasmid libraries, PCR1 was performed using Phusion Green Hot Start II High-Fidelity PCR 2x Master Mix with the following cycle conditions: 98°C for 2 minutes, [98°C for 10 seconds, 65°C for 20 seconds, 72°C for 1 minute] x the number of cycles was empirically determined for each sample by qPCR, 72°C for 2 minutes. 10 reactions for each plasmid library where 500 ng of the plasmid library was used for each 25 μL amplification reaction along with 500 nM primer mix.

**epegRNA Plasmid Library PCR1 Forward primer pool:** TCGTCGGCAGCGTCAGATGTGTATAAGAGACAG-[1N to 4N]-GAAAAAGTGGCACCGAGTCGG

**epegRNA Plasmid Library PCR1 Reverse primer pool:** GTCTCGTGGGCTCGGAGATGTGTATAAGAGACAG-[1N to 4N]-TTTGTGATGCTCGTCAGGGG

For genomic DNA, 48 25 μL PCR1 reactions with 500 ng of gDNA and 500 nM primer mix were run to ensure broad coverage across all samples. Genomic loci primers are the same as used previously for these amplifications (“Sequencing and analysis of individual pegRNA and epegRNA editing outcomes”). Amplifications were performed using Phusion Green Hot Start II High-Fidelity PCR 2x Master Mix with the following cycle conditions: 98°C for 2 minutes, [98°C for 10 seconds, 65°C for 20 seconds, 72°C for 1 minute]x30, 72°C for 2 minutes. As before, longer extension times were used to ensure complete extension for each cycle. PCR1 amplicons were all checked on 2% agarose as above.

PCR2 amplification for both plasmid libraries and genomic DNA from transfected cells was carried out using Nextera adapter PCR primers (listed in **table S29**) with the following conditions: 98°C for 2 minutes, [98°C for 10 seconds, 61°C for 20s, 72°C for 1 minute]x8, 72°C for 2 minutes. 5 μL of PCR1 products were used for each PCR2 reaction and each of the reactions was independently barcoded in PCR2. Following PCR2, all reactions for each genomic DNA or plasmid sample were pooled together, run on a 2% agarose gel, and gel extracted prior to quantification, library generation, and sequencing as described above (“Sequencing and analysis of individual pegRNA and epegRNA editing outcomes”).

CRISPResso2 (v2.2.6) was used to analyze demultiplexed reads. For both input plasmid libraries and genomic DNA from edited cells, CRISPRessoBatch was run with default settings using the unedited allele as the reference amplicon and desired allele with a 5 N insertion as Amplicon1 (-a parameter) and a 10nt quantification window on each side of the cleavage position for a total length of 20nt (-w parameter). Frequencies of aligned reads were extracted from the output Alleles_frequency_table files. Aligned reads were trimmed to 25nt flanking the cleavage position (−10nt and +10nt from the cleavage position to capture 5nt insertions +10nt of additional flanking sequence for edited alleles) and counts for reads with expected 5 nt insertions were quantified. On average, 52% (range 43-61%) of aligned reads from each gDNA sample matched an expected 5nt insertion and 43% (range 30-55%) of reads matched the unedited sequence, with only 4.6% (range 1.7-8.2%) of reads representing unintended alleles (i.e. indels or sequencing/PCR errors). For the input plasmid libraries, on average 88% (range 79-93%) of reads matched the expected epegRNA sequence with a 5nt insertion. For subsequent analysis, only aligned reads matched an expected 5nt insertion were considered. Normalized counts per million values were log2 transformed, replicates averaged, and used to calculate the fold change for each 5nt insertion sequence in the edited gDNA relative to the input plasmid library. 5nt insertion sequences were ranked by log2 fold change (see **table S7**) to nominate the top-performing LMs at each test site.

### Cloning and testing of top-performing LMs at 3 test sites

The top 94, 96, and 85 LMs for the HEK3, *EMX1*, and *RNF2* loci were tested individually. epegRNAs for each of these LMs were cloned, plasmid DNA was isolated, transfected, sequenced and analyzed as described above (“Testing of individual pegRNAs or epegRNAs” and “Sequencing and analysis of individual pegRNA and epegRNA editing outcomes”).

### Generation and testing of orthogonal edit site sequences

To make edit site sequences orthogonal to the human genome as well as the genomes of other widely used model organisms, we reasoned that by making base changes in the 3nt seed region of our edit site sequences that we could preserve the primer binding site (PBS) sequence for each edit site while largely maintaining the base identity and annealing properties of the RTT to generate novel edit site sequences that are not directly present in various genomes of interest. We generated two orthogonal sequence variants per edit site. We designed new protospacer sequences where we used the opposite Watson-Crick-Franklin pairing DNA base at each position from the endogenous human locus for the 3nt genomic seed for Ortho v1 (i.e. if the edit site sequence were 5’-GCT-3’ we would use 5’-CGA-3’), and used the reverse complement of the 3nt seed for Ortho v2 (i.e. if the sequence were 5’-GCT-3’ we would use 5’-AGC-3’).

### Sequencing and analysis of orthogonal edit site sequences

To experimentally evaluate our edit site orthogonalization strategies, we generated synthetic cassettes that harbored each of the *RNF2*, HEK3, and *EMX1* edit site sequences and necessary flanking sequences to support efficient prime editing for the original, unmodified genomic edit site sequences and for each of the two new orthogonalized edit sites. These edit sites were ordered as e-blocks from IDT and cloned using Gibson assembly into lineage tracing cassette (LTC) vectors. This vector introduces edit site sequences into the 3’ UTR of an mCherry reporter gene. These alleles were integrated into HEK293T cell genomes and cells harboring these constructs were isolated by sorting for mCherry+ cells using fluorescence activated cell sorting (FACS) two weeks after integration into the genome (representative sorting gates shown in **fig. S14**). Cells were subsequently transfected with epegRNAs targeting the corresponding sequences, cloned as described above (“Cloning of individual pegRNAs or epegRNAs”).

Genomic DNA was isolated using the Fast Lysis procedure described for sequencing HEK293T genomic loci 72 hours post-transfection. Two successive rounds of PCR were used to amplify both synthetic edit sites (PCR1 primers listed below) as well as endogenous genomic loci as described above (PCR1 primers listed in “Sequencing and analysis of individual pegRNA and epegRNA editing outcomes”) with 2 μL of gDNA and 500 nM primer mix using the Phusion Green Hot Start II High-Fidelity PCR 2x Master Mix with the following cycle conditions: 98°C for 2 minutes, [98°C for 10 seconds, 65°C for 20 seconds, 72°C for 30 seconds]x28, 72°C for 2 minutes. PCR1 amplicons were all checked on 2% agarose as above. PCR2 amplification was carried out using Nextera adapter PCR primers with the following conditions: 98°C for 2 minutes, [98°C for 10 seconds, 61°C for 20s, 72°C for 30 seconds]x8, 72°C for 2 minutes. 5 μL of PCR1 products were used for each PCR2 reaction and each of the reactions was independently barcoded in PCR2. All steps through sequencing using the Illumina Miseq v2 Reagent kit were performed as above (“Sequencing and analysis of individual pegRNA and epegRNA editing outcomes”). Experiments to re-test optimal LMs for orthogonalized edit site sequences were carried out using this same protocol.

**LTC Forward primer pool:** TCGTCGGCAGCGTCAGATGTGTATAAGAGACAG-[1N to 4N]-GAATCCAGCTAGCTGTGCAGC

**LTC Reverse primer pool:** GTCTCGTGGGCTCGGAGATGTGTATAAGAGACAG-[1N to 4N]-CCTTAGCCGCTAATAGGTGAGC

CRISPResso2 (v2.2.6) was used to analyze demultiplexed reads. For all prime editing yield quantification, CRISPRessoBatch was run with default settings using the unedited allele as the reference amplicon and desired allele as Amplicon1 (-a parameter) and a 10nt quantification window on each side of the cleavage position for a total length of 20nt (-w parameter). Editing yield was calculated as: (number of Amplicon1-aligned reads)/(total reads aligned to all amplicons). For all experiments, indel yields were calculated as: (number of indel-containing reads)/(total reads aligned to all amplicons).

### *In silico* characterization of orthogonalized edit site sequences

To further ensure compatibility of Ortho v1 edit site sequences for use in widely used model systems, we used the Cas-OFFinder *in silico* prediction algorithm (*95*) (v2.4.1) to identify potential CRISPR/Cas9-dependent off-target loci with three or fewer mismatches or bulges to our three OrthoV1 protospacers of interest. Cas-OFFfinder nominated 69, 174, 13, and 4 Cas9-dependent off-targets with 3 or fewer mismatches or bulges (MMB) for *H. sapiens*, *M. musculus*, *D. rerio*, and *D. melanogaster* genomes, respectively. There were no predicted genomic off-targets with 1 mismatch in any of the analyzed genomes (**Supp. Fig. 2D**). All predicted 2 mismatch off-targets were within intergenic or intronic regions of the genome with the exception of one in the promoter of the *apoea* gene in the *D. rerio* genome. Among the 251 putative genomic off-targets with 3 mismatches to our sgRNAs, only 4 were within exons of genes– 2 each in the human and mouse genomes– and 14 were in other regulatory genomic elements. When each of these 18 putative off-target loci were examined, we did not identify homologous sequences within 100nt downstream of the protospacer likely to support prime editing.

### LM detection cross hybridization calculations

To select lineage marks suitable for accurate hybridization probe readout, we used NUPACK (v4.0.1.9) (*149*) simulations to evaluate off-target binding for various LM sequence combinations. Each simulation involved hybridizing an equimolar mixture of DNA readout sequences (20nt complementary region) with an equimolar mixture of DNA edit site sequences containing the selected LMs. Simulations were conducted using a 100-fold excess of readout sequences (10 nM vs. 0.1 nM) at 43°C in a sodium ion concentration of 0.3 M. For each LM sequence, the free energy difference (ΔG) between correct and incorrect probe binding was estimated. Cross-hybridization was quantified as the fraction of edit site molecules bound by incorrect readout sequences. Separate simulations were performed for the Ortho v1 Edit Site 1, Edit Site 2, and Edit Site 3 sequences. Results from these simulations are reported in **table S8.**

To investigate the effect of LM insert length on probe cross-hybridization, we simulated mixtures of unedited edit site sequences and edit site sequences containing random LM inserts of varying lengths (2–8nt). For each LM length, simulations were repeated 10 times using 8 different LM sequences while keeping the complementarity length and other hybridization parameters constant. Results from these simulations are reported in **table S4.**

### Design and cloning of lineage tracing cassette libraries

With three edit site sequences selected, we finalized our lineage tracing cassette design. These cassettes consist of the following sequence organization: 5’-UCOE-EF1a promoter-mCherry- T7 promoter- T3 promoter- PacI digestion site- 30nt sequencing barcodes-sequencing primer- MERFISH decodable intBC- Forward primer for sequencing assays-sequencing intBC- PacI digestion site- Edit Site 1 and flanking sequences- Edit Site 2 and flanking sequences- Edit Site 3 and flanking sequences- Reverse primer for sequencing assays-polyA. For lentiviral construct designs, we included 500nt of constant sequence downstream for optimizing imaging readout experiments and used a BGH polyA sequence. For PiggyBac transposase-compatible vectors, we introduced this 500nt constant sequence before an SV40 polyA.

For testing of 24-mer pegArrays and epegRNA mismatch libraries, we generated a PiggyBac lineage tracing cassette plasmid library with a random 10nt sequencing barcode (LTCv0).

For *in vitro* single cell lineage tracing experiments, we generated a lentiviral lineage tracing cassette plasmid library with 125nt MERFISH barcodes with an additional 30nt sequencing barcode (LTCv1). Specifically, 125nt MERFISH barcodes were generated by choosing 6 among 21 of previously published MERFISH readout sequences (*150*) (B37-B66, 20nt for each) and concatenating them together with an “A” between each pair of readout sequences. 30nt sequencing barcodes were randomly generated.

For the *in vitro* imaging experiments and all *in vivo* experiments, we generated a lentiviral lineage tracing cassette plasmid library with matching 30nt sequencing barcodes and 183nt MERFISH barcodes (LTCv2). To design these orthogonal barcodes, we first randomly selected 9 of the 25nt orthogonal DNA among 240k published DNA barcodes (*151*), concatenated them in an randomly permuted order and selected the best permutation by checking if all 15mer are unique to this barcode. We repeated this process to select up to 6,000 candidate barcodes. We then predicted the self-binding partition function by “pfunc” from Nupack (*149*) and selected the candidate barcodes with highest free energy to minimize self binding. After filtering, we inserted the sequencing primer at position 30 of candidate barcodes and designated the first 30nt as sequencing barcodes, and then selected the next 183nt as MERFISH barcodes. We concatenated the 5’ sequence (T7, T3 and PacI site) and 56nt of 3’ sequence (Edit site 1) together as candidate intBCs for cross-hybridization testing. We first partitioned each 183nt MERFISH barcode into 6 non-overlapping 30nt segments, and used the reverse complement of these segments as candidate probes. To evaluate probe cross-hybridization, we enumerated the pairs of candidate intBCs, generated a Nupack Tube class with the candidate probes and these two candidate intBCs, and predicted the free energy and partition function for all possible bindings. After prediction, we summarized probe cross-hybridization for all candidate intBCs and retained intBCs with less than 1% of predicted average probe off-targeting probability. For barcodes that passed these requirements, we further filtered to have no 17nt matching to human, mouse, zebrafish and drosophila genomes.

As such, we ordered libraries of 2,171 intBCs (**table S6**) from Twist Bioscience and amplified this pooled library using Phusion Green Hot Start II High-Fidelity PCR 2x Master Mix with the following cycle conditions: 98°C for 2 minutes, [98°C for 10 seconds, 65°C for 20 seconds, 72°C for 1 minute] where the number of cycles for amplification were determined by quantitative PCR, followed by 72°C for 2 minutes. Given that libraries can be provided with variable quality, this qPCR determination was important for ensuring library quality. Following library amplification, samples were cleaned up with a QIAquick PCR purification column (Qiagen) according to manufacturer’s protocols. These amplified DNA fragments were assembled into PacI-digested and Quick CIP-treated (NEB) acceptor backbone via Gibson assembly using NEBuilder 2x HiFi DNA Assembly master mix as large scale 100 μL reactions described above. The entire 100 μL of Gibson assembly product was cleaned up using PCR cleanup columns (Qiagen) to remove salts prior to electroporation into MegaX cells as previously described. Colonies were scraped into a combined pool, grown out for 8 hours in 500 mL terrific broth (TB) containing 100 μg/mL carbenicillin, and midi-prepped using the ZymoPure II Plasmid Midiprep Kit (Zymo Research) following manufacturer’s procedures.

**LTCv2 Cloning Forward primer:** AATTAACCCTCACTAAAGGGATAATTTAATTAA
**LTCv2 Cloning Reverse primer:** GTCGTAATGACTAAGATGACTGCCATTAATTAA

### Sequence validation and analysis of lineage tracing cassette libraries

Lentiviral plasmid libraries were sequence confirmed as described for the 1,024 epegRNA libraries with modest differences. As before, plasmid libraries were amplified with two successive rounds of PCR as done to sequence genomic loci. PCR1 was performed using Phusion Green Hot Start II High-Fidelity PCR 2x Master Mix with the following cycle conditions: 98°C for 2 minutes, [98°C for 10 seconds, 65°C for 20 seconds, 72°C for 1 minute] where the cycle number was determined by qPCR, followed by 72°C for 2 minutes, and a hold at 12°C. PCR2, gel extraction, library preparation, and sequencing were carried out as before.

**intBC Sequencing Validation Forward primer:** TCGTCGGCAGCGTCAGATGTGTATAAGAGACAG-[1N-4N]- GTAATTAACCCTCACTAAAGG
**intBC Sequencing Validation Reverse primer:** TCGTCGGCAGCGTCAGATGTGTATAAGAGACAG-[1N-4N]- CATCGATACCTAATACGACTCACTATAGGGAGAG

To assess the balance of library elements, we used the FastX toolkit (v0.0.14) to mask low quality bases (<q20) and collapse reads. We removed common flanking sequences and summed reads matching the whitelist of intBC barcodes for sequences with Hamming distance <4 to account for sequencing errors and the length of intBC sequences.

### Plasmid design and cloning strategy for 8-mer and 24-mer pegArrays

To clone 24-mer arrays comprising an 8-mer or each of the edit sites, we designed a two-step Golden Gate assembly cloning strategy. For the first step, 8-mer arrays are cloned into three distinct acceptor backbones, one for each edit site where the first cloning step is carried out with the Type IIS restriction enzyme BsmBI (NEB, R0739L). These backbones can then be used for a second cloning step using the Type IIS restriction enzyme PaqCI (NEB, R0745L) for the final 24- mer assembly. The overhangs for the 8-mer assembly are the same for each of 8-mer backbones. The backbone overhangs are 5’-GGAG-3’ and 5’-CTAA-3’. The intervening 8 epegRNA inserts use the following overhangs: 5’-GGAG-INS 1-TGCC-3’, 5’-TGCC-INS 2-GCAA-3’, 5’-GCAA- INS 3-ACTA-3’, 5’-ACTA-INS 4-TTAC-3’, 5’-TTAC-INS 5-CAGA-3’, 5’-CAGA-INS 6-TGTG-3’, 5’-TGTG-INS 7-GAGC-3’, 5’-GAGC-INS 8-CTAA-3’. Fragments with these overhangs were ordered from either IDT or Twist Bioscience using **table S30** as a design guide. The final 24-mer array has the three 8-mers in the following order to optimize for LM balance: Edit Site 2, Edit Site 3, then Edit site 1. To generate 24-mers, sequence verified 8-mer arrays were digested with PaqCI to generate 8-mer array fragments with the following overhangs: 5’-GGAG-[Edit Site 2 8-mers]- TGTG-3’, 5’-TGTG-[Edit Site 3 8-mers]-GAGC-3’, 5’-GAGC-[Edit Site 1 8-mers]-CTAA-3’. These fragments could then be cloned into a 24-mer acceptor backbone digested with PaqCI. All golden gate acceptor plasmids are provided on Addgene.

### Cloning protocol for 8-mer pegArrays

Three different acceptor backbones were generated for 8-mer assemblies, one for each of the three edit sites. These acceptor backbones include an EF1a promoter driving a BFP reporter gene and have BsmBI type-IIS restriction cut sites. DNA fragments were ordered from Twist Bioscience that include a U6 Pol III promoter upstream of epegRNAs for each of the 8 LMs for a given edit site. Golden gate assemblies were performed using 3 nM of each U6–epegRNA fragment (typically 1 μL of each ∼500nt fragment diluted to 20 ng/μL), 1.1 nM of the relevant backbone (typically 1 μL of the ∼8200nt BsmBI pre-digested backbone diluted to 25 ng/μL), 2 μL of NEBridge Golden Gate Assembly Kit (BsmBI-v2) (NEB, E1602L), 2 μL T4 ligase buffer, and 7 μL of nuclease free water. Assemblies were carried out following manufacturer’s recommendations: 99 cycles of 5 minutes at 42°C and 5 minutes at 18°C before a 10-minute 60°C incubation and subsequent 4°C hold. These reactions were transformed into NEB Stable Competent *E. coli* (NEB, C3040), incubating on ice for 20 minutes prior to a 30 second heat shock at 42°C. After heat shock, bacteria were incubated on ice for 5 minutes prior to plating on 100 mg/mL carbenicillin resistant agar plates. Plates were incubated overnight at 30°C prior to picking colonies into 5 mL of 100 mg/mL carbenicillin-containing terrific broth, outgrowth overnight at 30°C, and DNA isolation using the Promega PureYield DNA Miniprep System. Some bacteria were expanded into 100 mL cultures overnight while isolated plasmids were diluted to 100 ng/μL and submitted for whole plasmid sequencing. DNA was isolated from these larger cultures using the ZymoPure II Plasmid Midiprep Kit. Plasmids were sequence confirmed by whole plasmid sequencing and further validated to have the correct insert by gel electrophoresis to be the correct size following digestion with PaqCI.

### Cloning protocol for 24-mer pegArrays

24-mer assemblies require four pieces of DNA that include a PaqCI digested 24-mer backbone and a PaqCI digested fragment for each edit site 8-mer. PaqCI digestions were performed overnight using 4 μL of CutSmart buffer, 4 μL of PaqCI activator, 2 μL of PaqCI, and 30 μL of the plasmid of interest. These digested fragments were gel extracted using the QIAquick Gel Extraction Kit (Qiagen) following manufacturer’s recommendations. Digested fragments were assembled into 30 μL PaqCI golden gate assemblies with 3.33 nM of the PaqCI digested 8-mer fragments (typically 1 μL of the ∼4000nt fragment diluted to 150 ng/μL), 1 μL of PaqCI pre-digested 24-mer acceptor backbone diluted to 25 ng/μL, 2.5 μL PaqCI enzyme, 1.25 μL PaqCI activator, 10 μL NEBridge Ligase Master Mix (M1100L), and water to bring the reaction to 30 μL total volume. Assemblies were carried out following manufacturer’s recommendations, carrying out 99 cycles of 5 minutes at 37°C and 5 minutes at 16°C before a 10-minute 60°C incubation and subsequent 4°C hold. All subsequent steps of transformation, plating, colony picking, and DNA prep and sequence confirmation were carried out as described for 8-mer assemblies above.

### Testing of 8-mer and 24-mer pegArrays

We introduced PEmax editor and lineage tracing cassettes (LTCv0) into B16-F10 cells using sequential PiggyBac transposition. 1,000,000 cells were nucleofected with 1.5 μg of transfer plasmid and 150 ng of Super PiggyBac Transposase Expression Vector (System Biosciences PB210PA-1) using the SF Cell Line 4D-Nucleofector Kit L (Lonza V4XC-3024) with program CM-130 and sorted for GFP+ (PEmax+) cells and GFP+/mCherry+ (PEmax+/lineage tracing cassettes+) at least 8 days after nucleofection to ensure stable expression. We then introduced 8-mer and 24-mer pegArray variants into GFP+/mCherry+ cells using the same nucleofection conditions and sorted GFP+/mCherry+/BFP+ (PEmax+/lineage tracing cassettes+/pegArray+) at least 8 days after nucleofection. Representative sorting gates shown in **fig. S14**. Genomic DNA was isolated using QuickExtract DNA Extraction Solution (Biosearch Technologies, QE09050) following the manufacturer’s protocol.

### Sequencing and analysis of 8-mer and 24-mer pegArrays

To assess pegRNA competition, lineage tracing cassettes were amplified and sequenced to quantify lineage mark installation efficiency. Two successive rounds of PCR were used to amplify lineage tracing cassettes with 2 μL of gDNA and 500 nM primer mix using the Phusion Green Hot Start II High-Fidelity PCR 2x Master Mix with the following cycle conditions: 98°C for 2 minutes, [98°C for 10 seconds, 65°C for 20 seconds, 72°C for 30 seconds]x28, 72°C for 2 minutes. PCR1 amplicons were all checked on 2% agarose as above. PCR2 amplification was carried out using Nextera adapter PCR primers with the following conditions: 98°C for 2 minutes, [98°C for 10 seconds, 61°C for 20 seconds, 72°C for 30 seconds]x8, 72°C for 2 minutes. 5 μL of PCR1 products were used for each PCR2 reaction and each of the reactions was independently barcoded in PCR2. Following PCR2, all reactions for each genomic DNA or plasmid sample were pooled together, run on a 2% agarose gel, and gel extracted prior to quantification, library generation, and sequencing as described above.

**LTC Forward primer pool:** TCGTCGGCAGCGTCAGATGTGTATAAGAGACAG-[1N to 4N]- GAATCCAGCTAGCTGTGCAGC

**LTC Reverse primer pool:** GTCTCGTGGGCTCGGAGATGTGTATAAGAGACAG-[1N to 4N]- CCTTAGCCGCTAATAGGTGAGC

CRISPResso2 (v2.2.6) was used to analyze demultiplexed reads. CRISPResso was run with default settings using the unedited allele as the reference amplicon and desired allele with a 5N insertion as Amplicon1 (-a parameter) and a 10nt quantification window on each side of the cleavage position for a total length of 20nt (-w parameter). Frequencies of aligned reads were extracted from the output Alleles_frequency_table files. Aligned reads were trimmed to 25nt flanking the cleavage position (−10nt and +10nt from the cleavage position to capture 5nt insertions +10nt of additional flanking sequence for edited alleles) and counts for reads with expected 5nt insertions were quantified. For subsequent analysis, only aligned reads matched an expected 5nt insertion were considered to calculate the relative editing efficiency for each pegRNA. Read counts for each 5nt insertion and array are reported in **table S9.**

### Cloning of a comprehensive epegRNA mismatch library

We designed a library of epegRNA protospacer mismatches that includes all possible single letter variants at positions 2-20 of an SpCas9 protospacer (where the NGG PAM is 21-23). We preserved a guanine (G) at position 1 to ensure efficient transcription by the human U6 promoter. Inspired by the CROP-seq vector (*97*), we cloned this epegRNA library into the 3’ UTR of a lentiviral genome (modified from Addgene plasmid #86708 to replace puromycin with BFP and introduce BsmBI epegRNA cloning site). This vector ensures that the sequence of each epegRNA could be captured in droplet-based single cell RNA sequencing experiments while producing an active copy of the epegRNA through the genome integration process. epegRNA protospacer mismatches were either synthesized and cloned by Twist Bioscience or ordered as eBlock fragments from IDT and cloned into the BsmBI-digested CROP-seq vector backbone using Gibson assembly. All protospacer mismatches are listed in **table S10**.

### Lentiviral production

Lentiviral particles were generated as previously described (*152*). Briefly, HEK293T/17 cells (ATCC CRL-11268) were maintained in DMEM (Thermo Fisher Scientific) supplemented with 10% (v/v) FBS and 1% (v/v) PSG at 37°C with 5% CO2. Cells were split 1:8 two days prior to use to ensure rapid cycling. 23,000,000 cells were plated per 15cm dish on the day prior to transfection. The next day, cells were ∼90% confluent and were transfected with the following transfection mix: 200 μL FuGENE HD transfection reagent and a DNA mixture with 1.3 pmol of psPAX2 (Addgene plasmid #12260), 0.72 pmol pMD2.G (Addgene plasmid #12259), and 1.64 pmol the transfer plasmid. DNA and Fugene mix was resuspended in a final volume of 3 mL of OptiMEM serum free media. This mixture was vortexed gently and incubated for 15 minutes at room temperature before adding dropwise to cells. 6 hours post-transfection, transfection media was removed and replaced with Opti-MEM I Reduced Serum Medium (OPTI-MEM) with GlutaMAX Supplement (Invitrogen, 31985088) supplemented with 5%FCS, 1 mM sodium pyruvate (Fisher Scientific), and 1x MEM nonessential amino acids (FisherScientific) (cOPTI-MEM) supplemented with 1x ViralBoost reagent (AlStem). 48 hours post-transfection, media was collected, spun at 3000g for 15 minutes to remove cell debris and concentrated using Lenti-X-Concentrator (TakaraBio, 631232) following the manufacturer’s instructions. Concentrated virus was resuspended in OPTI-MEM in 1% of the original culture volume without supplements. Lentiviral particles were subsequently aliquoted and frozen at−80°C.

### epegRNA mismatch library profiling to tune lineage tracing kinetics

We introduced lineage tracing cassettes where each of the three edit sites were concatenated together (LTCv0) using PiggyBac transposition into two mismatch repair-competent cell lines, B16-F10 (a melanoma-derived line; ATCC CRL-6475) and 4T1 (a breast cancer-derived line; ATCC CRL-2539). We also introduced the PEmax editor using PiggyBac transposition and sorted for cells that were both GFP+ and mCherry+ to select for editor and tracing cassette containing cells, respectively. For PiggyBac transposition, one million cells were nucleofected with 1.5 μg of transfer plasmid and 150 ng of Super PiggyBac Transposase Expression Vector (System Biosciences, PB210PA-1). We used the SF Cell Line 4D-Nucleofector Kit L (Lonza V4XC-3024) with program CM-130 for B16-F10 cells and the SE Cell Line 4D-Nucleofector Kit L (Lonza V4XC-1024) with program CM-150 for 4T1 cells. We then treated GFP+/mCherry+ cells with lentivirus expressing the CROP-seq BFP epegRNA library, sorted BFP+ cells, and passaged these cells for 28 days, collecting samples throughout the duration of the experiment. Representative sorting gates shown in **fig. S14**.

Single cell RNA sequencing (scRNA-seq) was performed on harvested cells to link epegRNA mismatches with lineage tracing editing rates. scRNA-seq libraries were prepared using the 10X Genomics Chromium Next GEM Single Cell 3ʹ Kit v3.1 following the manufacturer’s instructions followed by targeted enrichment of CROP-seq epegRNA and lineage tracing cassette transcripts based on published protocols (*81*, *153*). To increase diversity in Read2 during sequencing, we used a pool of primers with an 8nt stagger following the Illumina Read2 primer binding site.

Two successive rounds of PCR were used to prepare lineage tracing cassette libraries from amplified full-length cDNA. For the first PCR reaction (PCR1), 2 μL of amplified cDNA and 600 nM primer mix listed below (LTC_10X_PCR1_0-8_F, 10X_PCR1_R) were used. Amplifications were performed using KAPA HiFi HotStart ReadyMix with the following cycle conditions: 95°C for 3 minutes, [98°C for 20s, 65°C for 15s, 72°C for 15s]x19-25, 72°C for 1 minute. PCR2 amplification was carried out using 4 μL of a 1:25 dilution of PCR1 product and 300 nM of the reverse primer (10X_PCR2_R) listed below and Nextera i7 adapter PCR primers (listed in **table S29**) with the following conditions: 95°C for 3 minutes, [98°C for 20s, 72°C for 30 seconds]x12, 72°C for 1 minute.

**scRNA-seq Lineage tracing cassette amplification primers:** PCR1 forward (LTC_10X_PCR1_0-8_F): GTCTCGTGGGCTCGGAGATGTGTATAAGAGACAG -[1N to 8N]- GAATCCAGCTAGCTGTGCAGC
PCR1 reverse (10X_PCR1_R): ACACTCTTTCCCTACACGACG
PCR2 forward (Nextera i7): CAAGCAGAAGACGGCATACGAGATNNNNNNNNGTCTCGTGGGCTCGG
PCR2 reverse (10X_PCR2_R): AATGATACGGCGACCACCGAGATCTACACTCTTTCCCTACACGACGCTC

Three successive rounds of PCR were used to prepare epegRNA libraries from amplified full-length cDNA. For the first PCR reaction (PCR1), 2 μL of amplified cDNA and 300 nM primer mix listed below (pegRNA_10X_PCR1_F, pegRNA_10X_PCR1_R) were used. Amplifications were performed using KAPA HiFi HotStart ReadyMix with the following cycle conditions: 95°C for 3 minutes, [98°C for 20s, 65°C for 15s, 72°C for 15s]x29-33, 72°C for 1 minute. PCR2 amplification was carried out using 1 μL of a 1:25 dilution of PCR1 product and 300 nM of the primer mix listed below (pegRNA_10X_PCR2_0-8_F, 10X_PCR2_R) listed below with the following conditions: 95°C for 3 minutes, [98°C for 20s, 65°C for 15s, 72°C for 15s]x19, 72°C for 1 minute. PCR3 amplification was carried out using 4 μL of a 1:25 dilution of PCR2 product and 300 nM of the reverse primer (10X_PCR2_R) listed below and Nextera i7 adapter PCR primers (listed in **table S29**) with the following conditions: 95°C for 3 minutes, [98°C for 20s, 72°C for 30 seconds]x12, 72°C for 1 minute.

**scRNA-seq epegRNA amplification primers:**
PCR1 forward (pegRNA_10X_PCR1_F): ACACTCTTTCCCTACACGACG PCR1 reverse (pegRNA_10X_PCR1_R): TTTCCCATGATTCCTTCATATTTGC
PCR2 forward (pegRNA_10X_PCR2_0-8_F): GTCTCGTGGGCTCGGAGATGTGTATAAGAGACAG-[1N-to-8N]- CTTGTGGAAAGGACGAAACAC
PCR2/PCR3 reverse (10X_PCR2_R): AATGATACGGCGACCACCGAGATCTACACTCTTTCCCTACACGACGCTC

Following PCR amplification, lineage tracing cassette and epegRNA libraries were purified and size-selected using SPRI magnetic beads (0.9x single-sided selection) and quantified by BioAnalyzer (Agilent) to assess the size and purity of final libraries and Qubit double-stranded DNA high sensitivity Kit (Thermo Fisher Scientific) to determine the concentrations. Libraries were diluted to 4 nM and sequenced on a Nextseq 2000 (Illumina). All PCRs were first monitored to determine optimal cycle number by qPCR by adding 0.6x SYBR Green (Thermo) to the PCR reactions.

### scRNA-seq data processing

All scRNA-seq data were processed using the Cellranger software package (v7.1.0). To capture lineage information, a custom reference genome was constructed by appending the 2,171 lineage tracing cassette sequences to the mouse mm10 genome build. Both gene expression and lineage tracing cassette libraries were then aligned to this reference using default parameters. Since the reference contains all 2,171 30nt barcode sequences, the intBC of each lineage tracing cassette read can be determined based on which element in the reference it aligns to.

A custom Python script was used to extract lineage information from the BAM alignment file generated by Cellranger. This script iterates through all reads aligned to a tracing cassette sequence. For each read, the LM sequence at each edit site was determined based on the alignment. For each UMI, the number of supporting reads was calculated, and the LM for each edit site was set to the LM with the most read support (ties broken randomly). UMIs were then aggregated by intBC, cellBC, and the set of LMs detected across the three edit sites to generate a final LM table.

### scRNA-seq LM quality control

To generate high-quality lineage trees from scRNA-seq data, rigorous LM quality control is essential due to common issues such as PCR errors, sequencing errors, and ambient RNA contamination in tracing cassette reads. Since the levels of erroneous reads can vary between experiments–depending on sequencing depth and cell quality–and between lineage cassette integrations (intBCs) due to differences in expression levels, we applied adaptive thresholding to filter alleles in the LM table. First, for each 10X capture, we fit a two-component Gaussian mixture model to the distribution of reads/UMIs per allele and removed alleles corresponding to the lower peak of the distribution. Next, for each 10X capture and intBC, we applied the mixture model to the distribution of UMIs per allele, again filtering out alleles in the lower peak. The first step eliminates PCR and sequencing errors, which are characterized by low read support, while the second step removes alleles derived from ambient RNA, which exhibit low UMI support that varies according to the expression level of the intBC.

When a lineage tracer is read out using endogenous RNA expression, another source of error arises from the lag time between the integration of the LM into the genome and the point when RNA molecules containing that LM become the predominant species. This issue is exacerbated by the long RNA half-lives of lineage tracing cassettes, particularly in experiments conducted over short timescales, leading to errors in calling LMs that occur later in the experiment. To mitigate this problem, instead of resolving conflicting alleles by simply selecting the allele with the highest UMI count, we considered the edit state of the alleles. If the conflicting alleles differed at a single edit site–where one allele was edited and the other was not–we selected the edited allele, provided it accounted for >20% of the total UMIs for that intBC in the cell.

With only 2,171 distinct intBCs, duplicate integrations–where two tracing cassettes with the same intBC integrate into the genome of a single clone–can occasionally occur. If not filtered out, these duplicates could introduce errors, as they accrue different LMs, and we are unable to resolve which LM occurred at which integration. To address this, we identified and removed alleles from duplicate intBCs by detecting allele conflicts. For each clone and intBC, we calculated the total fraction of UMIs in cells containing more than one allele and excluded intBCs where this fraction exceeded 25%.

### scRNA-seq cell quality control

Low-quality cells in scRNA-seq experiments were filtered out based on mitochondrial content and total UMI counts. Cells with >10% of UMIs assigned to mitochondrial transcripts were excluded. For the kinetics experiment, cells with <100 UMIs assigned to the transcriptome were also removed. The UMI threshold varied between experiments due to differences in transcriptome sequencing depth. In the scRNA-seq of fully-edited clones and tracing cells with barcoding, the threshold was set at 1,000 UMIs instead of 100.

Lineage tracing enables robust and accurate detection of scRNA-seq doublets–droplets containing more than one cell–since conflicting alleles reveal the presence of multiple distinct lineages within a single droplet. Doublets were filtered out by identifying these conflicting alleles, as it is unlikely that two random cells would share the same set of LMs. Similar to the process used for removing duplicate intBCs, we calculated the total fraction of UMIs linked to intBCs with conflicting alleles for each cell and excluded cells where this fraction exceeded 25%.

### epegRNA kinetics analysis

epegRNA libraries were processed using the “CRISPR Guide Capture” feature of Cellranger. epegRNA protospacer mismatch calls supported by <3 UMIs were filtered out and careful quality control performed to exclude cells with multiple epegRNA integrations as these cells could confound our estimation of protospacer mismatch kinetics. Cells with multiple epegRNA calls were removed as well as cells with lineage cassette reads where more than one edit site had an LM installed.

To calculate the edit fraction over time for each epegRNA protospacer mismatch in each cell line, we first determined the fraction of intBCs with an edit in each cell. We then calculated the mean edit fraction across cells for a given epegRNA call at each time point. Time points with fewer than 20 cells for a particular protospacer mismatch were excluded due to excessive noise in these data points.

To estimate the edit rate for each epegRNA protospacer mismatch, we fitted a saturating exponential curve to the edit fraction data. The curve is defined by the equation:

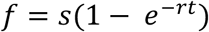

where *f* is the edit fraction, *t* is the time point (days), *r* is the edit rate (edits/[edit site * day]), and *s* is the saturation level which accounts for a small fraction of cells that do not undergo editing despite expressing the epegRNA. The curve was fitted using non-linear least squares, with *s* constrained to the range [0.8,1] and *r* constrained to the range [10^-4^,0.6]. Estimated edit rates in 4T1 and B16-F10 cells for each protospacer variant are reported in **table S10.**

### Generation of 4T1 PEtracer cells with high numbers of integrated tracing cassettes

4T1 cells were maintained in typical culture conditions. Infections with lineage tracing cassette lentiviral libraries were carried out by plating 100,000 cells per well of a 6-well dish 18-24 hours prior to transduction. Immediately before adding the virus, media was changed on these cells to include 10ug/mL polybrene (Millipore Sigma). 50 μL of high titer virus was incubated on cells for 24 hours prior to a media change and outgrowth of cells. The brightest 10% of mCherry expressing cells were sorted 1 week after transduction to isolate cells with high integration numbers. Representative sorting gates shown in **fig. S14**. This process was repeated serially to increase integration count until the average number of integrations in the population was over 10 as evaluated by qPCR using the following primer pairs and a single multiplicity of infection (MOI) line where infection rate was ∼20% as an internal 1x integration control. qPCR reactions were carried out using 50 ng of genomic DNA isolated using DNeasy blood and tissue kit (Qiagen) with 500 nM primer mix and 1X DyNAmo ColorFlash SYBR Green Master Mix (Thermo) and the following cycling conditions: 95°C for 7 minutes, [95°C for 10 seconds, 60°C for 30 seconds]x40. Relative integration number was calculated by the ddCt method compared to β-actin.

<u>Mouse β-Actin Control Forward:</u> TTTGATGTCACGCACGATTT
<u>Mouse β-Actin Control Reverse:</u> AGGGCTATGCTCTCCCTCAC
<u>mCherry Forward:</u> AGGGCGAGATCAAGCAGAG
<u>mCherry Reverse:</u> CTCGTTGTGGGAGGTGATGT

### Generation of 4T1 PEtracer cells edited to completion (fully-edited cells)

4T1 were nucleofected to introduce a no mismatch 24-mer pegArray (all epegRNAs in the array had protospacers with no mismatches that would slow their editing rates). Nucleofections with 4T1 cells were carried out using the Lonza 4-D nucleofector according to manufacturer’s instructions with the SE buffer, pulse code CM-150, 1.8 μg of donor pegArray plasmid and 180 ng of PiggyBac transposase plasmid (SBI). Cells were allowed to recover for 48 hours, split for >10 days to ensure loss of unintegrated plasmid, and BFP+ cells were isolated using Fluorescence Activated Cell Sorting (FACS) on a BD FACS Aria II (BD Biosciences). BFP+ cells were then engineered with high numbers of integrated tracing cassettes (LTCv2) by serial lentiviral infection as described above (“Generation of 4T1 PEtracer cells with high numbers of integrated tracing cassettes”). BFP+/mCherry+ cells were plated and infected with PEmax-P2A-GFP lentivirus as described for the tracing cassette lentivirus. BFP+/mCherry+/GFP+ triple positive cells were FACS sorted >5 days post transduction, passaged for >3 weeks, and then re-sorted at 5 cells/well. Representative sorting gates shown in **fig. S14**. Clones were expanded, evaluated by bulk sequencing, and pooled to cover all unedited and edited states.

### scRNA-seq of fully-edited 4T1 cells

Fully-edited 4T1 cells were passaged in RPMI as described above. Prior to droplet-based single cell sequencing using the 10X Genomics Chromium Next GEM Single Cell 3ʹ Kit v3.1, cells were split, counted, and diluted according to manufacturer’s protocols. Lineage tracing cassette libraries were generated as described above (“epegRNA mismatch library profiling to tune lineage tracing kinetics”).

We anticipated recombination during cloning and lentiviral production steps and therefore register a link between MERFISH and sequencing intBCs in fully-edited 4T1 cells by generating targeted sequencing libraries from amplified cDNA using T3 specific primers (located 5’ of both the MERFISH and sequencing intBCs).

Two successive rounds of PCR were used to prepare intBC libraries from amplified full-length cDNA. For the first PCR reaction (PCR1), 2 μL of amplified cDNA and 600 nM primer mix listed below (T3_10X_PCR1_1-4_F, 10X_PCR1_R) were used. Amplifications were performed using KAPA HiFi HotStart ReadyMix with the following cycle conditions: 95°C for 3 minutes, [98°C for 20s, 65°C for 15s, 72°C for 15s]x25, 72°C for 1 minute. PCR2 amplification was carried out using 4 μL of a 1:25 dilution of PCR1 product and 300 nM of the reverse primer (10X_PCR2_R) listed below and Nextera i7 adapter PCR primers (listed in **table S29**) with the following conditions: 95°C for 3 minutes, [98°C for 20 seconds, 72°C for 30 seconds]x12, 72°C for 1 minute.

**scRNA-seq intBC amplification primers:**
PCR1 forward (T3_10X_PCR1_1-4_F): GTCTCGTGGGCTCGGAGATGTGTATAAGAGACAG -[1N to 4N]- GTAATTAACCCTCACTAAAGG
PCR1 reverse (10X_PCR1_R): ACACTCTTTCCCTACACGACG
PCR2 forward (Nextera i7): CAAGCAGAAGACGGCATACGAGATNNNNNNNNGTCTCGTGGGCTCGG
PCR2 reverse (10X_PCR2_R): AATGATACGGCGACCACCGAGATCTACACTCTTTCCCTACACGACGCTC

Following PCR amplification, lineage tracing cassette and intBC libraries were purified and size-selected using SPRI magnetic beads (0.9x single-sided selection) and quantified by BioAnalyzer (Agilent) to assess the size and purity of final libraries and Qubit double-stranded DNA high sensitivity Kit (Thermo Fisher Scientific) to determine the concentrations. Libraries were diluted to 4 nM and sequenced on a Nextseq 2000 (Illumina). All PCRs were first monitored to determine optimal cycle number by qPCR by adding 0.6x SYBR Green (Thermo) to the PCR reactions.

Reads from the T3 library were processed similarly to reads from the standard tracing cassette library, with read/UMI and UMI thresholds determined using Gaussian mixture models. The 183nt MERFISH barcode for each T3 read was identified based on alignment, while the 30nt sequencing barcode was extracted from the read sequence and corrected using a whitelist of known barcodes.

To map 183nt MERFISH barcodes to 30nt sequencing barcodes, we assigned each MERFISH barcode to a sequencing barcode in each cell based on UMI support. We then calculated the fraction of cells supporting each mapping for each clone. This approach allowed us to confidently assign MERFISH barcodes to 29 of the 31 sequencing barcodes present in the fully-edited 4T1 cells, revealing multiple instances of barcode swapping (**fig. S5B** and **table S11**). In cases of barcode swapping, integrations are referred to by their MERFISH barcode identifier.

### Clone calling using intBCs

Non-negative matrix factorization (NMF) was used to call clones which we define as groups of phylogenetically related cells that share a set of intBCs. Given the UMI counts for each intBC in each cell after quality control filtering, we represented the data as a cell-by-intBC matrix *X.* NMF was then applied to factorize *X* into the product of two matrices, *W* and *H*:

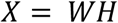

Here, *W* is a cell-by-clone matrix representing the assignment of cells to clones, while *H* is a clone-by-intBC matrix representing the assignment of intBCs to clones. The number of clones for each experiment was manually determined based on visual inspection of the matrix *X*.

To filter out scRNA-seq doublets, which often contain cells from distinct clones, we identified cells with intBC sets that more closely matched the combination of two clones than any single clone. First, cells were putatively assigned to clones using the matrix *W*. Next, for each intBC, we assigned it to a clone if it was detected in >50% of the cells within that clone. Using this clone-to-intBC mapping, we calculated the Jaccard similarity between the set of intBCs detected in each cell and the intBC sets for both individual clones and all pairwise combinations of clones. Cells were assigned to the clone with the highest Jaccard similarity unless the highest match was to a pair of clones, in which case the cell was marked as a doublet. If the Jaccard similarity between a cell of interest and all possible clone assignments was <0.5, the cell was marked as unassigned.

### Quantification of lineage tracing data accuracy in fully-edited cells

To quantify the accuracy of lineage tracing data in fully-edited 4T1 cells, we generated a whitelist (**table S11**) for each clone containing the LMs installed at each edit site for each intBC. This whitelist was derived from the scRNA-seq data, where cells were grouped into four clones based on their intBCs, as described above. For each clone, we then identified the most common LM at each edit site for each intBC, which formed the basis of the whitelist.

We assessed the accuracy and sensitivity of both intBC and LM detection using this whitelist. Based on each cell’s clone assignment, we identified the expected set of intBCs and LMs and compared these to the actual intBCs and LMs detected. For intBCs, we calculated the true positive and false negative rates from the fraction of expected intBCs that were detected, and the false positive rate from the fraction of detected intBCs that were not expected. For LMs, we calculated overall call accuracy, as well as accuracy per edit site and per LM. In scRNA-seq data, the LM detection rate matches the intBC detection rate, as LMs can be called for all detected intBCs. However, in imaging data, the LM detection rate is slightly lower than the intBC detection rate, since for some amplicons the intBC can be accurately called but the LM at one or more edit sites cannot.

### Microscope setup for image acquisition

Image acquisition was performed using a custom-built microscope system, as previously described (*154*) with some modifications. The system was built around a Nikon Ti2-U microscope body with a Nikon CFI Plan Apochromat Lambda D 60x oil immersion objective with 1.42 NA. The system also included a Nikon CFI Plan Fluor 10x objective with 0.3 NA for quick identification of the sample position. Illumination of samples was through a Lumencor CELESTA light engine (a fiber-coupled solid-state laser-based illumination system) with the following wavelengths: 405 nm, 477 nm, 545 nm, 637 nm, and 748 nm. This system was used with a penta-bandpass dichroic mirror (Semrock, FF421/491/567/659/776-Di01-25X36), a penta-bandpass emission filter (Semrock, FF01-441/511/593/684/817-25), and a penta-bandpass excitation filter (Semrock, FF01-391/477/549/636/741-25). A scientific CMOS camera (Hamamatsu C14440 with factory calibration for single-molecule imaging) was used for image acquisition. Each camera field of view (FOV) consisted of 2,304 x 2,304 pixels, with a camera pixel corresponding to 107 nm in the X and Y dimensions in the imaging plane for the 60x oil immersion objective with 1.42 NA. Sample position in three dimensions was controlled using a XYZ stage (Ludl Electronic Products). A custom-built auto-focus system was used to maintain a constant focal plane over prolonged time intervals. This was achieved by comparing the relative position of two IR laser (Thorlabs, LP980-SF15) beams reflected from the glass-fluid interface and imaged on a separate CMOS camera (Thorlabs, CS165MU1).

These different components were controlled using a National Instruments Data Acquisition card (NI PCIe-6321, X Series DAQ) and custom software (see the “Software for controlling experimental components” section below).

### MERFISH fluidics system configuration

The fluidics system consisted of several core components similar to previous designs with some modifications (*154*): a syringe pump to control flow speed and volume; a set of daisy-chain-connected valves to select flow-in buffers; a flow chamber in which the sample was mounted; a set of tubing, connectors, and needles; and a 3-D printed chassis to hold a 24-well deep-well plate (Agilent, 204023-100). Four MVP4 High Torque valves (Hamilton, 94747-02) with 8-way connector heads (Hamilton, 97472-01) were daisy-chain connected (port 8 of each valve is connected to the input of the next valve), and the output of the first valve was connected to the FCS2 flow chamber (Bioptechs, 03060319-2-NH-30) in which samples were placed together with a 0.5 mm thick rubber gasket (Bioptechs, DIE# F18524). The chamber output was connected to the input port of the syringe pump (Hamilton, PSD6) which controlled the flow speed and volume. The output port of the syringe pump was further connected to a waste collecting vessel.

This system was controlled using a custom software (see the “Software for controlling experimental components” section below). Overall, this system allowed for 24 rounds of hybridization. We also added two 3-way adaptor sets (Bioptechs, 162003-1) and connected these two adaptors directly with tubing to allow fluidic bypass of the FCS2 chamber, which allows us to perform washing of the fluidic system without unmounting the sample if needed.

### Software for controlling experimental components

All system components were controlled using custom-built software available at https://github.com/ZhuangLab/storm-control. The software package is composed of several main modules that work in concert:

- “Hal” is the software package used to control and synchronize all illumination and microscope components. We note that in some cases it is necessary to install drivers for components which are not included in this package, for example, the camera for autofocus system (Thorlabs, CS165MU1) requires Thorlabs official drivers. Specific imaging parameters including laser intensities, z-scan steps, exposure time for each frame and the sequence of shutter wrapped in xml files are loaded and executed by Hal. These parameter files were generated by Python scripts provided in our Github repository (see “Code availability” below).
- “Steve” is a module used to generate mosaic images (i.e., a composite of many individual fields of view (FOV)) of a sample to select FOV for imaging in experiments.
- “Kilroy” is the software used to control fluidics components, and to define pre-programmed sequences of operations to be performed as sets (e.g., the set of operations that initiate a new round of hybridization).
- “Dave” can issue commands to both Hal and Kilroy and is used to automate the performance of data collection by pre-programmed the complete set of fluidics system and microscope operations used for an experiment (i.e. the order and time-lag in which fluidics and imaging steps are to be performed).

The general outline of an experiment is as follows: before the experiment starts, Hal and Kilroy are loaded with the parameters and specifications to be used. After the sample is loaded and the chamber is filled with the imaging buffer, a mosaic image of the DAPI channel is taken using Steve, and FOVs of interest are selected. A file is then generated to specify the sequence of operations throughout the entire experiment and is loaded to Dave, together with the coordinates of the selected regions of interest. The rest of the experiment is automated. If the number of rounds in an experiment exceeds the capacity of the flow system, we can specify a Dave file to automatically run up to the capacity of the system, then use the bypass tubing to clean up the fluidic system without perturbing the mounted sample. After cleaning up, all buffers are replaced and a new Dave file is created. This is repeated until all rounds of imaging are completed.

### Design of intBC encoding probes

A combinatorial detection scheme for our 2,171 intBCs was inspired by MERFISH experiments (*56*) and used a modified Hamming distance 4, Hamming weight 6 21-bit codebook derived from covering designs (https://www.dmgordon.org/cover/, with parameters: v=21, k=6, t=5). Each of the 21 total bits in the codebook are assigned to a unique readout sequence and each intBC was assigned to a six bit code in the codebook (**table S6**). We designed six encoding probes per intBC consisting of a 30nt targeting sequence and three readout sequences. For each intBC, all assigned readout sequences are represented across the set of encoding probes. The 183nt MERFISH barcode within each integration barcode was partitioned into 6 non-overlapping 30nt segments. These 30nt sequences have been tested for cross-hybridization during intBC design procedures, therefore the reverse-complement of these 30nt sequences were directly used as targeting sequences in probes. Readout sequences were appended to each targeting sequence separated by an “A”, with all assigned readout sequences evenly represented across the set of encoding probes designed for that intBC. 20nt forward and reverse primers were then added to the 5’ and 3’ of these probes. Then the fully assembled probes were further screened for cross-hybridization, so that any probes with a 17nt segment reverse-complementing to other probes were removed. Full probe sequences can be found in **table S12**.

### Design of LM and unedited probes

A unique probe was designed to hybridize to each LM installed at its cognate edit site. Specifically, edit site sequences modified with each LM were generated *in silico* and probes were designed to hybridize to each modified edit site. Probes were designed to be 20nt in length and centered around each 5nt inserted LM. For each LM, a specific MERFISH readout sequence was chosen and two copies of this readout sequence were appended to the 5’ side of each edit site/LM probe. All LM probes were directly purchased from IDT. Several combinations were tested and the final set of probes with the highest signal to noise ratios (SNRs) were merged. Full probe sequences can be found in **table S13**.

### Design of tracing cassette common sequence probes

Target sequences of common bits were directly chosen from the 500nt common sequence located 3’ of the edit sites in the lineage tracing cassette (Addgene # pending). Common sequence probes were designed to target 24 30nt sequences within this 500nt common sequence. Two 20nt MERFISH readout sequences (**table S16**) were appended to the 5’ and 3’ of these targeting sequences flanked by a set of forward and reverse primers. Full probe sequences could be found in **table S14**.

### Encoding probe amplification

Encoding probes were ordered as oligo pools from Twist Bioscience. Each oligo pool was first dissolved in 100 µL TE buffer (Thermo Fisher Scientific, AM9849). Encoding probes were amplified from the template library using a previously described amplification protocol (*154*) with additional modifications:

1. The initial oligo pool was amplified using limited-cycle PCR for approximately 11-15 cycles dependent on the library size, where the reverse primer used for this amplification included a T7 promoter sequence to enable *in vitro* transcription in the next step. PCR primer sequences are listed below.
2. The resulting PCR product was purified via column purification (DNA Clean & Concentrator-25, Zymo Research, D4033).
3. The purified PCR product underwent further amplification and conversion to RNA by a high-yield *in vitro* T7-mediated transcription reaction (HiScribe® T7 Quick High Yield RNA Synthesis Kit, NEB, E2050).
4. The resulting RNA product was purified by column (Monarch® RNA Cleanup Kit, NEB, T2050).
5. The purified RNA product was converted back to single-stranded DNA (ssDNA) by a reverse transcription reaction (Maxima H Minus Reverse Transcriptase, Thermo Fisher Scientific, EP0753) using the forward primer from step 1 in this protocol. The final product was purified by adding 3x volume of the self-made SPRI beads, eluted in TE buffer followed by an ethanol precipitation to concentrate to 25 nM for each probe in TE buffer. This probe stock could be stored in −20°C. All primers were purchased from IDT.

**Probes for common sequence in lineage cassette were amplified by:**
primer: CGGGTTTCGTTGCGCACACC
primer: TAATACGACTCACTATAGGGCTTGTGCATCGCGCCAAAGA

**Probes for intBC were amplified by:**
primer: CCCGCAATGGCTGACAACCG
primer: TAATACGACTCACTATAGGGATTGCCGCATGGTTTCCG

**Probes for 124-gene MERFISH library were amplified by:**
primer: CCCGCAATGGCTGACAACCG
primer: TAATACGACTCACTATAGGGATTGCCGCATGGTTTCCG

**Probes for 175-gene MERFISH library were amplified by two sub-pools:**
Amplification of 150-gene Sub-pool Forward primer: CCCGCAATGGCTGACAACCG
Amplification of 150-gene Sub-pool Reverse primer: TAATACGACTCACTATAGGGATTGCCGCATGGTTTCCG

Amplification of 25-gene Sub-pool Forward primer: CGGGTTTCGTTGCGCACACC
Amplification of 25-gene Sub-pool Reverse primer: TAATACGACTCACTATAGGGCTTGTGCATCGCGCCAAAGA

### Glass coverslip cleaning and silanization

40 mm diameter, 0.17 mm thickness glass coverslips (Bioptechs, 40-1313-03192) were first cleaned by placing coverslips in a plastic holder (Entergris, A23-0215) and performing 3x 5 minute rinses with MilliQ water in a glass dish (Electron Microscopy Sciences, 70312-31) prior to a 30-minute incubation in a chemical hood at room temperature with a mixture of equal volumes of Methanol (Sigma, 34860) and 37% HCl (Sigma-Aldrich 258148). After this incubation, 4x 5-minute MilliQ water rinses were performed prior to 2x 5-minute washes with 70% ethanol. Coverslips were gently blown dry using compressed air and left to fully dry.

Cleaned coverslips were placed in a glass holder and incubated for 30 minutes at room temperature in a mixture of 1L chloroform (Sigma-Aldrich 319988), 1 mL triethylamine (Sigma-Aldrich 81101), and 2 mL allyl-trichlorosilane (Sigma-Aldrich 107778) which were all stored and handled according to provider’s instructions. Using glass pipetting agents is essential to generating high-quality coverslips. Following this incubation, coverslips were washed by performing 2x 5-minute chloroform washes followed by a 1x 5-minute wash in 100% ethanol. After this ethanol wash, coverslips were gently blown dry by compressed air and left to fully dry. Following silanization, coverslips were stored in a dry, dark place in a sealed container with a dedicated set of desiccant cartridges (VWR, 76538-672).

### Poly-D lysine coating silanized glass coverslips

Silanized coverslips were incubated with 0.1 mg/mL Poly-D lysine (PDL; Thermo Fisher Scientific, A3890401) overnight at room temperature in a 3-D printed box with parafilm sealing top. Following PDL coating, coverslips were cleaned with 3x 5-minute rinses with MilliQ water followed by a 1x rinse with 100% ethanol before being gently blown dry with compressed air and left to dry. Coverslips were stored in 3-D printed cases at 4°C with the same desiccant cartridge and sterilized with ultraviolet light for 20 minutes immediately prior to use.

### Seeding of fully-edited cells prior to imaging

Fully-edited cells were cultured as described above prior to seeding 400,000 cells on cleaned, silanized, and PDL-coated 40 mm diameter round coverslips in 60 mm diameter petri-dishes (Corning, 430166) in RPMI media supplemented with 10% FBS and 0.1% PSG. Samples were then prepared for FISH-based lineage readout using either the “Zombie” or “In-gel T7” protocols described in the following sections. 59 fields of view (FOVs) were imaged for one coverslip prepared with the “Zombie” protocol and 69 FOVs for one coverslip prepared with the “In-gel T7” protocol as enabling comparison of these methods.

### Sample preparation for Zombie experiments

The Zombie protocol was adapted from previously published methods (*42*, *54*). In brief, samples were washed 2x with 1x PBS for 5 minutes each and subsequently fixed with a 3:1 (vol:vol) mix of methanol and acetic acid at room temperature for 20 minutes. Cells were then washed 4x with 1x PBS prior to a 1x wash with nuclease-free water. Samples were subsequently incubated with T7 *in vitro* transcription mix (MEGAscript Transcription Kit; Invitrogen) supplemented with 2.5 mM aminoallyl-UTP (Thermo Fisher Scientific, R1091) at 37°C for 16 hours. After transcription, cells were fixed with 4% formaldehyde in PBS for 20 minutes at room temperature followed by 2x washes with 1x PBS prior to the same hybridization as in the section “Lineage probe hybridization” below.

### Sample preparation for In-gel T7 amplification

Samples were washed 2x with 1x PBS for 5 minutes each and then fixed by treatment with 4% paraformaldehyde in 1x PBS for 10 minutes at room temperature. Samples were subsequently washed three times with 1x PBS and stored in 1x PBS supplemented with 1:500 RNase Inhibitor Murine (NEB, M0314L) for up to 8 weeks.

The fully-edited cells fixed on coverslip were embedded in a hydrogel prior to tissue clearing to reduce tissue autofluorescence and off-target probe binding. First, the sample was incubated in gel solution consisting of 300 mM NaCl, 4% (vol/vol) of 19:1 acrylamide/bisacrylamide, 60 mM Tris-HCl pH 8, 0.1% (vol/vol) TEMED, 1/40,000 (vol/vol) 505/515 nm fiducial beads (Invitrogen F8803), and 0.5% (wt/vol) ammonium persulfate for 5 minutes at room temperature. We then placed a 60 μL droplet of the gel solution onto a hydrophobic glass slide treated with GelSlick (Lonza). The coverslip bearing the sample was then inverted onto the droplet to form a uniform layer of monomer solution. The sample was allowed to polymerize completely for >1 hour at room temperature.

The coverslip bearing the gel-embedded sample was then removed from the glass slide with a thin razor blade. The sample was then incubated in digestion buffer containing 2% (wt/vol) SDS, 0.5% (vol/vol) Triton-X-100 (Sigma, X100), and 1% (vol/vol) Proteinase K (NEB P8107S) in 2x SSC for 24 hours at 47°C and a subsequent 24 hours at 37°C. Following digestion, the sample was washed by performing 6x 20 minute washes with 2x SSC buffer supplemented with 0.2% (vol/vol) Rnase Inhibitor Murine (NEB) at room temperature. The sample was further digested in 800 mM Guanidine HCl (IBI Scientific, IB05080), 1 mM EDTA, 50 mM Tris-HCl pH=8.0, and 0.5% Triton-X-100 supplemented by 1% (vol/vol) Proteinase K for 24 hours at 37°C.

After Guanidine HCl digestion, samples were transferred to new petri-dishes with fresh 2x SSC for one quick wash followed by a 10-minute incubation in 2x SSC supplemented with 1 mM phenylmethylsulfonyl fluoride (PMSF) protease inhibitor (Thermo Fisher Scientific, 36978). Samples were then thoroughly washed by performing 5x 15-minute rinses with 2x SSC on a shaker followed by 3x 15-minute washes with nuclease free water on the shaker. The remaining water was aspirated from the coverslips, and each coverslip was inverted onto a 100 µL droplet of T7 *in vitro* transcription mix (MEGAscript Transcription Kit; Invitrogen AM1334) supplemented with 2.5 mM aminoallyl-UTP (Thermo Fisher Scientific, R1091) on a parafilm-covered 60 mm diameter petri-dish. Transcription was carried out for 16 hours at 37°C. Optionally, samples were remounted onto fresh T7 transcription mixture after this 16 hours incubation for an additional 3 hours at 37°C.

After transcription, samples were fixed with 4% formaldehyde solution in 1x PBS for 20 minutes at room temperature followed by 2x washes with 1x PBS. After these washes, probe hybridization was carried out as described in the section “Lineage probe hybridization” below.

### Lineage probe hybridization

Following either the Zombie or in-gel T7 amplification protocols above, probes for lineage intBCs, common sequences, and LMs were hybridized similar to published MERFISH protocols (*155*). Two separate hybridizations were performed: in the first, probes for the intBCs and common sequences were hybridized, while in the second LM-encoding probes were hybridized.

For the first hybridization, samples were washed once with 2x SSC prior to equilibration in encoding-probe wash buffer (30% (v/v) formamide [Thermo Scientific, AM9344] in 2x SSC) supplemented with 0.1% Tween-20 (Sigma, P9416) for 10-15 minutes at room temperature. Wash buffer was then aspirated from the coverslip, and coverslips were inverted with the sample side facing down onto a 50 µL droplet of probe mixture on a parafilm-covered 60 mm petri-dish. The probe mixture comprised encoding probes for each intBCs at 1 nM and encoding probes for the two-color common sequence at 10 nM in 2x SSC with 30% v/v formamide, 0.1% wt/v yeast tRNA (Life Technologies) and 10% v/v dextran sulfate (Sigma, D8906). Probe hybridization was carried out by incubating at 37°C for 24 hours in a humidified chamber. After hybridization, samples were washed 1x with encoding-probe wash buffer for 30 minutes at 47°C to remove excess probes prior to a 1x wash with 2x SSC before proceeding directly to LM probe hybridization.

Samples were equilibrated in LM-probe wash buffer (10% (v/v) formamide in 2x SSC) supplemented with 0.1% Tween-20 for 10-15 minutes at room temperature. Wash buffer was then aspirated from the coverslip, and each coverslip was inverted with the sample side facing down onto a 50 µL droplet of probe mixture on a parafilm-covered 60 mm petri-dish as above. This probe mixture comprised 300 nM of each LM probe (directly purchased from IDT) in 2x SSC with 30% v/v formamide, 0.1% wt/v yeast tRNA (Life Technologies) and 10% v/v dextran sulfate (D8906, Sigma). Samples were incubated at 37°C for 24 hours in a humidified chamber. After hybridization, the sample was washed 2x with the encoding-probe wash buffer for 15 minutes at room temperature to remove excess probes.

Following these two hybridizations, samples were embedded in a second 4% poly-acrylamide gel as described in the section “In-gel T7 amplification for fully-edited cells” above to further immobilize T7 amplicons. After this embedding, samples can be stored at 4°C in 2x SSC supplemented with 1:1000 Rnase Inhibitor Murine for up to 4 days.

### Imaging of integration barcodes and lineage marks

After lineage probe hybridization and secondary gel-embedding, samples were assembled into the FCS2 flow chamber (Bioptechs, 060319-2). Samples were imaged by the home-built imaging platform described in the “Microscope setup for image acquisition” and “MERFISH fluidics system configuration” sections. All fluid exchanges on the microscope use the custom-built fluidics system described in the “Fluidics system configuration” section. For each coverslip, nuclear staining by DAPI was used to help select FOVs of interest from mosaic images generated at 10x magnification using “Steve” as described in “Software for controlling experimental components”.

Throughout the course of imaging, we performed multiple rounds of hybridization and imaging. Specifically, we imaged 2 colocalizing bits for common sequences used to nominate potential amplions in round 0; 21-bit combinatorial scheme for integration barcode readout in rounds 1-7, and 27 sequential bits for LMs and unedited states in rounds 8-16 (three colors per round). We designed 51 adaptor probes to convert bit-specific readout sequences in the encoding probes into common readout sequences. Each adaptor probe consists of: a 5’ sequence that is complementary to one of the bit-specific readout sequences used in encoding probes, and a 3’ sequence with two copies of the reverse-complement of one common readout sequence (**table S16**) (*154*). We designed 3 common readout sequences in total. Each common readout probe contains a readout sequence that is complementary to the adaptor probe and has a fluorescent dye (Alexa750, Alexa647 or Atto565) linked to the 5’ of the oligo via a disulfide bond **(table S15)**.

### For each round, samples were imaged as follows unless otherwise described

1. Samples were hybridized with 3 adaptors probes labeling 3 different bits at 100 nM in secondary hybridization buffer (2x SSC, 30% v/v formamide) for 15 minutes at room temperature.
2. Samples were washed with the secondary hybridization buffer for 3 minutes at room temperature.
3. Samples were then hybridized with a set of fluorescently labeled oligonucleotide probes termed “common readout probes” (synthesized by IDT) (**table S15**) at 20-25 nM in secondary hybridization buffer for 15 minutes at room temperature.
4. Samples were washed again with the secondary hybridization buffer, for 3 minutes at room temperature.
5. Samples were then incubated and kept in imaging buffer comprising 5 mM 3,4-dihydroxybenzoic acid (Sigma, P5630), 100 µM of Trolox and Trolox quinone mixture (generated by UV radiation of a Trolox solution ∼50 µM of Trolox quinone final quantified by UV-vis spectrometry; Sigma, 238813), 1:500 recombinant protocatechuate 3,4-dioxygenase (rPCO; OYC Americas, 46852904), 1:1000 Murine RNase inhibitor (NEB,

M0314L), 10 mM Tris-HCl pH=7.5 (Thermo Fisher Scientific, 15568025), and 5 mM NaOH (to adjust pH to 8.0; Sigma, S2770) in 2x SSC.

1. Samples were imaged using a 60x oil immersion objective, and signals for the common readout probes and fiducial beads were collected in the 748 nm, 637 nm, 545 nm, and 488 nm channels, respectively at a rate of 10 Hz. All channels were imaged for 50-60 consecutive 0.6 µm-thick z-stacks for each FOV of interest.
2. After imaging all FOVs, the signal from the previous round was extinguished by incubating the sample in cleavage buffer comprising 2x SSC, 30% formamide supplemented with 50 mM Tris (2-carboxyethyl) phosphine (TCEP; Sigma, 646547) for 10 minutes at room temperature. TCEP treatment cleaves the disulfide bond connecting fluorophores to readout probes. Cleavage buffer also contained 167 nM common readout probes without fluorophores to block unoccupied common readout sequences on the adaptor probes from interfering with the next round of hybridization.

All rounds of imaging were carried out as described above except for the first round. For the first round, before the imaging buffer step (step 5), samples were additionally washed in 2x SSC once and then incubated in 2x SSC containing 2.5 µg/mL DAPI (Thermo Fisher Scientific, D1306) for 10 minutes to stain nuclei. DAPI signal was collected in the 405 nm channel, and either the 405 nm channel or all channels in this round were imaged for 50-60 consecutive 0.6 µm-thick z-stacks for each FOV of interest, depending on the experiment. Nuclear DAPI signal was imaged together with common sequences in round 0, which was later used as reference for segmentation and fine alignments described here: “Sample alignment between MERFISH and lineage imaging”.

### Nuclei segmentation for *in vitro* samples

Cellpose (v3.1.0) was used to segment cell nuclei in 3-D based on DAPI signal (*109*). For each field of view (FOV) the DAPI image was adjusted to account for the illumination profile of the microscope and then downsampled by a factor of four in the x-y plane. Cellpose was run on a GPU using the "nuclei" model with a diameter of 25 pixels and a cell probability threshold of −4. The resulting nuclei masks were converted into polygons with a tolerance of 0.5 pixels.

Nuclei polygons from all FOVs were then aggregated. Since adjacent FOVs overlap, the same nuclei could be segmented in multiple FOVs, leading to overlapping polygons that required correction. For large overlaps (polygons from neighboring z planes that overlap >20% of their area), the two polygons were merged, as they were likely the same nuclei segmented twice. For smaller overlaps, the intersecting volume was subtracted from the smaller polygon, as these were likely segmentation errors. After aggregating all polygons and correcting overlaps, polygons with a volume <500 μm³ were removed.

3-D nuclei polygons, represented as a collection of 2-D polygons for each z-slice, were used to assign T7 amplicons and MERFISH transcripts to cells, as described in detail below. For 2-D visualization, nuclei outlines were generated by computing the union of all 2-D polygons along the z-axis.

### T7 amplicon detection and quantification

After T7 amplification, each lineage tracing cassette integration should be marked by a local population of RNA molecules centered at the site of transcription. These amplicons were detected using hybridization probes targeting the common sequence shared by all integrations. To maximize the detection rate, the common sequence was probed in two independent rounds of hybridization. To identify amplicons, the two hybridization images were maximum-projected, and a 2-D unsharp mask (sigma = 10 pixels) was applied to reduce background. Spherical intensity peaks were then identified in 3-D using the Laplacian of Gaussian method (sigma = 2-10 pixels, minimum intensity = 200). The spot detection parameters were optimized to prioritize recall of real amplicons, accepting false positives that could be filtered out during integration barcode decoding.

Following amplicon detection, the intensity of each hybridization round for each amplicon was quantified. A 2-D unsharp mask (sigma = 10 pixels) was applied to each channel, and then the maximum signal intensity within a 10-pixel diameter circle in the x-y plane centered at each amplicon position was calculated. This quantification was performed in 3-D to resolve amplicons that overlapped in the z-dimension. To correct for x-y drift between imaging rounds, fiducial beads imaged in each round were used for registration. The drift was calculated using phase cross-correlation between bead images in each round compared to the first round, and amplicon positions were adjusted accordingly.

### T7 amplicon integration barcode decoding

The intBC for each T7 amplicon was determined by probing the 183nt MERFISH barcode across 7 rounds of 3-color imaging. After detecting potential T7 amplicons and quantifying their intensity across the 21 bits as described above, data from all fields of view were combined into a *N-by-21* intensity matrix, *X,* where *N* is the total number of potential amplicons. With this representation, intBC decoding becomes an optimization problem, where we seek to assign each row in *X* to the most likely intBC codeword (21-bit Hamming distance 4, Hamming weight 6 code) accounting for variability in T7 amplification and probe hybridization efficiency across rounds. Following the approach of Moffit et al. (2016) we use an expectation maximization (EM) algorithm to solve this optimization problem (*156*).

First, *X* is color normalized to correct for differences in laser power and fluorophore intensity by rescaling each column so that its mean intensity is consistent across channels. Then, to account for variability in amplicon intensity and probe hybridization efficiency, *X* is normalized by the dot product of mean spot and bit intensities:

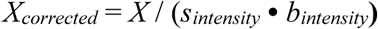

Here, *s_intensity_* is defined as the mean intensity of each spot across the “1” bits indicated by its corresponding codeword, and *b_intensity_* is the mean intensity of each bit across spots (after normalization by spot intensity). Initial estimates are obtained by setting *s_intensity_* to the 95th percentile of each row and *b_intensity_* to the 95th percentile of each column (after scaling by *s_intensity_*). Finally, *X_corrected_* is adjusted for each bit’s noise profile:

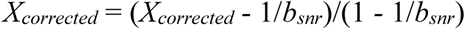

where *b_snr_* is the signal-to-noise ratio for each bit; values in *X_corrected_* >1 are clipped to 1.

In the expectation step, *X_corrected_* is used to assign each spot to an intBC codeword from the predefined list of 2,171 valid binary codewords (listed in **table S6)**. A KDTree is used to efficiently perform a nearest-neighbor search that identifies the intBC codeword minimizing the Euclidean distance to each column in *X_corrected_.* Spots with a distance ≥1.7 between the codeword and corrected intensity vector are masked out. Spots with a distance ≥1.7 are masked out, a threshold that balances error correction with the exclusion signals that do not correspond to true integration amplicons.

In the maximization step, intBC identity assignment in the previous expectation step is used to update *s_intensity_*, *b_intensity_*, and *b_snr_,* where *s_intensity_* is recalculated for all rows, while only spots with confidently assigned intBCs contribute to the updates of *b_intensity_* and *b_snr_*. For each bit, *b_snr_* is computed as the ratio of the mean intensity for spots where the bit is present (“1” in the codeword) to the mean intensity for spots where it is absent (“0” in the codeword).

After 10 iterations–sufficient for convergence–each amplicon is decoded based on its assigned intBC. Amplicons with a matching distance of ≥1.7 are filtered out and excluded from downstream analysis.

### T7 amplicon lineage mark decoding

Each LM is detected by a separate hybridization probe, so decoding the eight LMs for each edit site as well as the unedited state requires 9 rounds of 3-color FISH imaging (27 bits total, 9 bits per edit site x 3 edit sites). For spots with confident intBC assignments, data from all fields of view were combined into an N-by-27 intensity matrix *X,* as described above. *X* is then color-balanced by rescaling each column so that its mean intensity is consistent across channels, correcting for differences in laser power and fluorophore brightness.

To select an LM for each edit site, we extract the relevant columns from *X* to form an N-by-9 intensity matrix, *X_site_*. Each row in *X_site_* is normalized so that the sum of its values equals 1, accounting for variation in overall spot intensity. While simply choosing the LM corresponding to the maximum value in each row of *X_site_* works well, we found that training a logistic regression classifier further improves decoding accuracy. The classifier learns to account for variations in probe affinity and cross-hybridization that affect the intensity profile of each LM across hybridization rounds.

We trained the logistic regression classifier using the known linkage between intBCs and LMs in our fully-edited cells (see “Quantification of lineage tracing data accuracy in fully-edited cells” for details). For each row in *X_site_*, we determined the ground-truth LM based on the decoded intBC and then trained the classifier to predict this value. *In vitro* decoding accuracies for both our in-gel tissue-clearing protocol and the standard Zombie protocol were evaluated using 5-fold cross-validation. The classifier was subsequently retrained on all *in vitro* data from the in-gel protocol and then applied to decode all subsequent experiments, including *in vivo* data from fully-edited cells, which serve as an out-of-sample control.

Separate classifiers were trained for each edit site. Accuracies were reported for all decoded LMs as well as for the subset of LMs confidently decoded by each classifier (with prediction probability p ≥0.7). Only LMs with p ≥0.7 were used in downstream analyses such as tree reconstruction.

### Assignment of T7 amplicons to cells

Decoded T7 amplicons were assigned to nuclei segmentation masks in 3-D using the geopandas package (v1.0.1). Using the 3-D coordinates of each amplicon and the 3-D polygons representing nuclei, amplicons falling within a polygon were assigned to the corresponding nucleus, then to account for segmentation errors, amplicons located just outside a mask (<4 µm) were assigned to the nearest nucleus. The Euclidean distance from each amplicon to its assigned mask was recorded, and amplicons outside segmentation masks were only used in tree reconstructions if no amplicon with the same intBC was detected within the mask.

### Imaging readout LM quality control

Similar to scRNA-seq LM quality control (see “scRNA-seq LM quality control” for reference), we identify a set of high-confidence LM calls for each intBC in every imaged cell. We first remove amplicons with poor LM detection by excluding those with a mean LM assignment probability <0.7 across the three edit sites. To account for instances where the same amplicon is detected multiple times by the spot detection algorithm, we group intBCs with identical LM sets within each cell and retain only the brightest spot from each group. To eliminate duplicate integrations (i.e., two lineage tracing cassettes sharing the same intBC), we exclude intBCs detected in more than one amplicon with different LMs in >40% of the cells for a given clone. Additionally, to remove segmentation doublets–cases where two cells are not properly separated–we filter out cells with >50% conflicting amplicons. Finally, to uniquely assign LMs to intBCs, we resolve conflicting amplicons by selecting the brightest spot in each cell, with a preference for those contained within the nuclear mask.

After quality control the accuracy of image-based intBC and LM decoding was quantified using fully-edited cells as described in the “Quantification of lineage tracing data accuracy in fully-edited cells” section.

### Cloning and evaluation of puromycin/blasticidin lentiviral static barcode libraries

Lentiviral plasmids expressing puromycin or blasticidin resistance genes were cloned with Gibson assembly as described above. We introduced a 10N random barcode that could be efficiently assayed using scRNA-seq-based capture of 3’ polyadenylated transcripts. These libraries were cloned by introducing oligonucleotide libraries ordered from IDT using Gibson assembly. The same strategy used for 5N epegRNA libraries described above (see section “Cloning of a comprehensive 1,024 5 nucleotide (nt) LM epegRNA libraries” for details) was used where top and bottom oligo libraries were ordered, cloned, and transformed before libraries were pooled, expanded through outgrowth for 8 hours and Midiprepped as detailed above. Lentiviral plasmid libraries were sequence confirmed as described for the epegRNA libraries with modest differences. As before, plasmid libraries were amplified with two successive rounds of PCR as done to sequence genomic loci. PCR1 was performed using Phusion Green Hot Start II High-Fidelity PCR 2x Master Mix with the following cycle conditions: 98°C for 2 minutes, [98°C for 10 seconds, 64°C for 20 seconds, 72°C for 1 minute] where the number of cycles was empirically determined by qPR, followed by a 72°C for 2 minutes. PCR2, gel extraction, library preparation, and sequencing were carried out as before.

**Puromycin 10N library top:** CAGAAAGCCTGGCGCCTGACNNNNNNNNNNGTTTAAACCCGCTGATCAGCCTCGAC TG
**Puromycin 10N library bottom:** CAGTCGAGGCTGATCAGCGGGTTTAAACNNNNNNNNNNGTCAGGCGCCAGGCTTTC TG
**Blasticidin 10N library top:** TTATGTGTGGGAGGGCTAACNNNNNNNNNNGTTTAAACCCGCTGATCAGCCTCGACT G
**Blasticidin 10N library bottom:** CAGTCGAGGCTGATCAGCGGGTTTAAACNNNNNNNNNNGTTAGCCCTCCCACACATA A

**Forward primer for puromycin library amplification:**
TCGTCGGCAGCGTCAGATGTGTATAAGAGACAG -[1N to 4N]-
CCTGGTGCATGACCAGAAAGC
**Forward primer for blasticidin library amplification:** TCGTCGGCAGCGTCAGATGTGTATAAGAGACAG -[1N to 4N]- GTGAATTGCTGCCCTCTGGTTATGTG
**Reverse primer for puromycin and blasticidin library amplification:** GTCTCGTGGGCTCGGAGATGTGTATAAGAGACAG -[1N to 4N]- GCACAGTCGAGGCTGATCAG

To assess the balance of library elements, we used FastX toolkit (v0.0.14) to mask low quality bases (<q20) and collapse reads. We removed common flanking sequences and summed reads for unique barcode sequences. Lentivirus was produced from these plasmid libraries as described above (“Lentiviral production”).

### *In vitro* tracing with ground-truth barcoding readout with scRNA-seq

4T1 cells with high numbers of integrated LTCv1 tracing cassettes were nucleofected with a pegArray tuned to a 2-to-3-week timescale (24-mer comprises: edit site 2 protospacer mutant 8, edit site 3 protospacer mutant = 34, edit site 1 protospacer mutant = 32; see **table S10**). >10 days post-nucleofection, BFP+/mCherry+ double positive cells were isolated by FACS sorting and expanded. On day 0 of the experiment, tracing cassette and pegArray containing cells were subsequently infected with PEmax-T2A-GFP lentivirus. 48 hours post-infection (on day 2), BFP+/mCherry+/GFP+ triple positive cells were sorted as 2 cells/well into 96-well tissue culture-treated plates (Corning). Representative sorting gates shown in **fig. S14**. On day 5, sorted cells were infected with 10 µL of the puromycin static barcode lentiviral library. Media was changed on Day 6 to prevent continued infection events. On day 7, cells were infected with 10 µL of the blasticidin static barcode lentiviral library. On day 8, cells were split into a fresh TC-treated 96-well plate to continue expansion of the clones. Robust wells were expanded until treatment on day 12 with 4 µg/mL blasticidin-containing selection media. Cells were passaged in blasticidin selection media until day 14 when cells were split into media containing both 4 µg/mL blasticidin and 2 µg/mL puromycin. Double selection continued until day 16, at which points cells were split, counted, and droplet-based single cell sequencing was performed by pooling one to three clones per capture for a total of 6 clones and 3 captures using the 10X Genomics Chromium Next GEM Single Cell 3ʹ Kit v3.1. Lineage tracing cassette libraries were generated as described above (“epegRNA mismatch library profiling to tune lineage tracing kinetics”).

### Sequencing and analysis of *in vitro* tracing with ground-truth barcoding readouts

Two successive rounds of PCR were used to prepare puromycin and blasticidin barcode libraries from amplified full-length cDNA. For the first PCR reaction (PCR1), 2 μL of amplified cDNA and 600 nM primer mix listed below (Puro_10X_PCR1_1-4_F/Blast_10X_PCR1_1-4_F, 10X_PCR1_R) were used. Amplifications were performed using KAPA HiFi HotStart ReadyMix with the following cycle conditions: 95°C for 3 minutes, [98°C for 20s, 65°C for 15s, 72°C for 15s]x18-20, 72°C for 1 minute. PCR2 amplification was carried out using 4 μL of a 1:25 dilution of PCR1 product and 300 nM of the reverse primer (10X_PCR2_R) listed below and Nextera i7 adapter PCR primers (listed in **table S29**) with the following conditions: 95°C for 3 minutes, [98°C for 20s, 72°C for 30 seconds]x12, 72°C for 1 minute.

**intBC amplification primers:**
PCR1 forward (Puro_10X_PCR1_1-4_F): GTCTCGTGGGCTCGGAGATGTGTATAAGAGACAG -[1N to 4N]- CCTGGTGCATGACCAGAAAGC

PCR1 forward (Blast_10X_PCR1_1-4_F): GTCTCGTGGGCTCGGAGATGTGTATAAGAGACAG -[1N to 4N]- GTGAATTGCTGCCCTCTGGTTATGTG

PCR1 reverse (10X_PCR1_R): ACACTCTTTCCCTACACGACG

PCR2 forward (Nextera i7): CAAGCAGAAGACGGCATACGAGATNNNNNNNNGTCTCGTGGGCTCGG

PCR2 reverse (10X_PCR2_R): AATGATACGGCGACCACCGAGATCTACACTCTTTCCCTACACGACGCTC

Following PCR amplification, lineage tracing cassette and Puro/Blast barcode libraries were purified and size-selected using SPRI magnetic beads (LTC libraries: 0.9x single-sided selection, Puro/Blast barcode libraries: 1.2x single-sided selection) and quantified by BioAnalyzer (Agilent) to assess the size and purity of final libraries and Qubit double-stranded DNA high sensitivity Kit (Thermo Fisher Scientific) to determine the concentrations. Libraries were diluted to 4 nM and sequenced on a Nextseq 2000 (Illumina). All PCRs were first monitored to determine optimal cycle number by qPCR by adding 0.6x SYBR Green (Thermo) to the PCR reactions.

scRNA-seq processing and quality control were performed as described in the “scRNA-seq data processing” section, with additional puromycin and blasticidin barcode libraries aligned using Cellranger to a custom reference. This reference included puromycin and blasticidin resistance cassette sequences with ‘N’s indicating the positions of the 10nt random barcodes. A custom Python script was used to extract the barcode sequence from each read aligned to these elements, generate a consensus barcode for each UMI, and aggregate UMIs by barcode for each cell. To separate barcode calls corresponding to real integrations from sequencing errors and ambient RNA, for each 10x capture a Gaussian mixture model was fitted to the distribution of UMIs per barcode per cell. Barcode calls corresponding to the lower peak were excluded as likely errors.

Quality control, clone calling, and phylogenetic tree reconstruction for each clone using neighbor joining were performed as described above.

### Algorithm for calling barcode groups

A greedy algorithm was used to define barcode groups, which are clusters of cells sharing a set of puromycin or blasticidin barcode sequences. Starting with UMI counts for each barcode in each cell after quality control filtering, the algorithm iteratively performs the following steps until all cells are assigned to a barcode group or no viable seeds remain:

1. <u>Select a seed:</u> Choose the barcode detected in the highest fraction of unassigned cells that has not been used in a previous iteration.
2. <u>Identify the cluster:</u> Identify all other barcodes present in more than 80% of the cells where the seed barcode is detected.
3. <u>Assign cells:</u> Assign all unassigned cells to the barcode group if the barcodes in the cluster account for more than 80% of the total barcode UMIs in those cells.
4. <u>Validate the group:</u> Retain the barcode group if at least 5 cells are assigned; otherwise, discard the group and mark the cells as unassigned.

This algorithm accurately assigns the vast majority of cells to barcode groups, accounting for cases where groups are distinguished by multiple barcodes and instances where barcodes are occasionally reused between groups (**fig. S6**).

### Quantification of agreement between phylogeny and barcode groups

The Fowlkes–Mallows index (FMI), a metric for quantifying the similarity between two clusterings, was used to assess the agreement between the phylogeny and barcode groups derived from puromycin and blasticidin static barcoding. FMI is defined as:

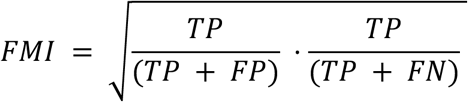

where *TP* is the number of true positives, *FP* is the number of false positives, and *FN* is the number of false negatives.

To calculate the FMI score for each clone, barcode groups were compared to clades representing the descendants of the ancestral cell that originally received each barcode. The lowest common ancestor (LCA) of each barcode group was inferred by computing the FMI score for all internal nodes in the phylogeny and selecting the node that maximized the FMI for that group. For each clone, FMI scores were calculated separately for puromycin and blasticidin barcode groups, using phylogenies reconstructed with both neighbor joining (NJ) and UPGMA. Cells lacking a barcode excluded from the FMI calculation. FMI scores were also calculated after randomly permuting the barcode group associated with each cell in the phylogeny. FMI values are reported in **table S17**.

### Inference of ancestral LM states

To estimate branch lengths in the phylogeny and collapse branches that are not supported by a lineage mark (LM), we inferred the ancestral LM state for each internal node in the tree using the Sankoff algorithm (*157*). This algorithm was chosen because it allows for low-probability LM-to-LM transitions, thereby reducing the impact of noise and errors in phylogeny reconstruction on ancestral state inference. For each edit site, the algorithm minimizes the total transition cost across the tree by assigning ancestral states that are most consistent with the LMs observed at the leaves. We implemented the following transition costs:

- Unedited → unedited: 0
- LM → same LM: 0
- Unedited → LM: 0.6
- LM → unedited or different LM: 1

These constraints reflect the assumption the state of an edit site should remain the same unless an LM is installed while allowing for some LM-to-LM transitions to account for errors.

### Inference of branch lengths

After reconstructing phylogenies and inferring ancestral states, we used ConvexML (v2.0.0) to estimate branch lengths in the tree (*158*). This maximum likelihood algorithm integrates a statistical model of the editing process with inferred ancestral LM states to efficiently estimate branch lengths. ConvexML assumes that editing occurs at a constant rate, meaning that longer branches–representing ancestral cells that take longer to divide–tend to accumulate more LMs than shorter branches. The agreement between our kinetics data and the expected saturating exponential curve suggests that the constant editing rate assumption holds true for PEtracer data for B16-F10 cells *in vitro* and for 4T1 both *in vitro* and *in vivo* (**fig. S4E** and **table S10**).

To generate the final phylogenies, we collapsed all branches that were not supported by a LM. This step introduced multifurcations in the tree, reflecting cases where the relationships between descendant cells could not be fully resolved, such as when one or more cell divisions were not marked by an LM. To perform this collapse, we conducted a reverse depth-first traversal of the tree’s edges. For each edge, if the inferred LM state of a parent node matched that of its child (i.e., no new LMs were acquired along that branch), we removed the edge and directly connected the child to its grandparent. This approach simplified the tree structure, preserving only branches supported by the available lineage tracing data.

Branch lengths were rescaled to reflect the tracing duration for each experiment, and node depths– defined as the number of edges from the root–were calculated for the collapsed tree. For each phylogeny, the “average tree depth” was determined by computing the mean depth of its leaves.

### Plotting phylogenetic trees

Phylogenetic trees were stored using TreeData (v0.1.3, https://github.com/YosefLab/treedata) and visualized with Pycea (v0.0.1, https://github.com/YosefLab/pycea). The plots display inferred branch lengths, with internal nodes sorted by subtree size. When intBCs are shown, they are ordered based on the maximum frequency of lineage marks (LMs) at the corresponding edit sites, positioning earlier edits on the left. High-confidence clades were manually selected and numbered according to their location in the phylogeny. Note that similar clade colors do not necessarily indicate close phylogenetic relationships due to multifurcations and arbitrary sorting of internal nodes.

### Downsampling analysis of phylogenetic reconstruction accuracy

To assess the impact of detection rate and number of edit sites on the accuracy of phylogenetic reconstruction, we performed a downsampling analysis on the scRNA-seq lineage tracing data with ground-truth barcodes. For each clone, we simulated detection rates ranging from 0.1 to 1 (in increments of 0.1) by randomly dropping out LMs to achieve the target detection rate. Similarly, we simulated different numbers of edit sites ranging from 5 to 45 (in increments of 5) by randomly removing edit sites. Phylogenies were then reconstructed using neighbor joining, and the Fowlkes– Mallows index (FMI) was calculated for puromycin and blasticidin barcode groups to quantify reconstruction accuracy. If the target detection rate or number of edit sites exceeded the detection rate or number of edit sites for that clone no downsampling was performed and the true detection rate or number of edit sites was reported. Each simulation was repeated 10 times per parameter combination, with FMI scores for each iteration reported in **tables S19,20** and mean FMI scores reported in the text.

### Comparison of pairwise phylogenetic, character, and spatial distance

To compare various distance metrics between cells in a phylogeny, 20,000 random cell pairs were selected, unless the total number of possible pairs was fewer than 20,000, in which case all pairs were used. The distances between each pair were calculated as follows:

1. <u>Spatial distance:</u> The Euclidean distance between the x-y coordinates of the two cells. For phylogenies spanning multiple sections, pairs from different sections were excluded.
2. <u>LM distance:</u> The Hamming distance between the set of LMs for each cell in the pair, as described in the “Distance-based tree reconstruction” section.
3. <u>Phylogenetic distance:</u> The LCA of the cell pair was identified using breadth-first search, and the total path length between the LCA and each of the two cells computed. We use inferred branch lengths for this calculation, so path lengths represent the estimated number of days since the LCA divided.

### *In vitro* 4T1 colony sample preparation and imaging

4T1 cells with high numbers of LTCv2 integrated lineage tracing cassettes and a 24-mer pegArray with no protospacer mismatches that edits to completion within 5 to 7 days were generated as described above (“Generation of 4T1 PEtracer cells edited to completion (fully-edited cells)”). These cells were infected with PEmax-T2A-GFP lentivirus on day 0 and sorted 48 hours later for high expression of the mCherry-containing tracing cassettes, consistent expression of the BFP-linked pegArray, and for consistent expression of GFP-linked PEmax. Representative sorting gates shown in **fig. S14**. After sorting, 11,000 actively-editing 4T1 cells were seeded on cleaned, silanized, and PDL-coated 40 mm diameter round coverslips in 60 mm diameter petri-dishes with RPMI media as described for fully-edited *in vitro* cell samples above.

After 6 days, cells were fixed by treating with freshly made 4% paraformaldehyde in 1x PBS for 10 minutes prior to 3x washes with 1x PBS and storage in 1x PBS supplemented with 0.2% RNase Inhibitor Murine (NEB, M0314L) for up to 8 weeks. Samples were prepared for FISH-based lineage readout as described in the “Sample preparation for In-gel T7 amplification” section and then imaged and analyzed as described above. Representative colonies of varying sizes were imaged across 262 FOVs of one coverslip.

### Phylogenetic reconstruction of *in vitro* 4T1 colonies

Segmented nuclei were grouped into spatially distinct colonies using the Density-Based Spatial Clustering of Applications with Noise (DBSCAN) algorithm (ε = 60 µm) (*159*), with colonies containing fewer than 25 cells filtered out. Since colonies could originate from multiple single cells that landed next to each other and therefore consist of more than one clone, clone calling was performed within each colony as described in the “Clone calling using intBCs” section. The only difference between scRNA-seq and imaging-based clone calling was that spectral clustering was used for initial assignments instead of non-negative matrix factorization (NMF) due to the lower proportion of doublets in imaging data.

LM quality control and phylogenetic tree reconstruction using neighbor joining were performed as described above. However, based on thresholds determined through *in silico* benchmarking and downsampling of scRNA-seq lineage tracing data, only nuclei with >60% intBC detection were included in tree reconstruction with neighbor joining. In some cases, phylogenies contained multiple clones that could not be distinguished by the clone-calling algorithm due to similar intBC compositions. These phylogenies were manually split by clone based on visual inspection of LM data: if no single LM was shared by all cells within a phylogeny, major clades marked by distinct LMs were separated into independent phylogenies.

Branch length estimation was performed as described in the “Inference of branch lengths” section and comparison of phylogenetic and spatial distances performed as described in the “Comparison of pairwise phylogenetic, character, and spatial distance” section. Statistics including number of cells and intBC detection rate for the 64 clones are listed in **table S20**.

### Generation of primary 4T1 tumors

4T1 cells engineered with PEmax editor and lineage tracing cassettes (LTCv0) were generated as described above (“epegRNA mismatch library profiling to tune lineage tracing kinetics”) and barcoded using low MOI lentiviral infection with an sgRNA library that co-expresses PuroR-T2A-BFP (Addgene #1000000091) as described above (“Lentiviral production” and “Generation of 4T1 PEtracer cells with high numbers of integrated tracing cassettes”). Cells were selected with puromycin (2 µg/mL) for five days starting 72 hours after lentiviral infection. 500,000 or 1,000,000 cells were subcutaneously injected into the mammary fat pad of female Balbc/J mice. Following injection, mice were monitored in accordance with IACUC guidelines prior to sacrifice at six weeks post-injection. Mice were sacrificed by CO_2_ asphyxiation and primary tumor, liver, and lung tissue collected.

### Single cell RNA sequencing of primary 4T1 tumors

To inform the design of MERFISH gene panels, we generated scRNA-seq libraries from cancer, immune, and stromal cells isolated from primary 4T1 tumors as well as the lungs and livers of two tumor-bearing mice. Tissue was dissociated into single cell suspension by mincing followed by enzymatic digestion with 300 U/mL collagenase/100 U/mL hyaluronidase (STEMCELL Technologies) and 150 µg/mL DNase I (Sigma Aldrich) in RPMI 1640 (Thermo Fisher Scientific) at 37°C with rotation for 25 minutes (tumor) or 20 minutes (lung/liver). Homogenized tissue was smashed through a 70 μm sterile strainer and incubated with ACK Lysing Buffer (Thermo Fisher Scientific) for 5 minutes at room temperature. Single-cell suspensions were stained in flow buffer (2% fetal bovine serum in 1x PBS) with the following antibodies (1:500 dilution):

- anti-CD45 (clone 30-F11, Alexa Fluor 700, Biolegend Cat. No. 103127, RRID AB_493714)
- anti-CD11b (clone M1/70, APC, Biolegend Cat. No. 101211, RRID AB_312794)
- anti-CD11c (clone N418, APC/Cyanine7, Biolegend Cat. No. 117323, RRID AB_830646)
- anti-TCRb (clone H57-597, PE/Cyanine7, Biolegend Cat. No. 109221, RRID AB_893627)
- anti-Ly-6G/Ly-6C (Gr-1) (clone RB6-8C5, Brilliant Violet 711, Biolegend Cat. No. 108443, RRID AB_2562549)
- anti-CD19 (clone 6D5, Brilliant Violet 605, Biolegend Cat. No. 115539, RRID AB_2563067)

Cells were stained for viability using eF506 dye (1:1000, eBioscience). Cells were sorted on BD FACS Aria II for viable cancer cells (mCherry^+^ CD45^-^), stromal cells (mCherry^-^ CD45^-^), neutrophils (CD45^+^ Gr-1^+^) and other immune cells (CD45^+^ Gr-1^-^). After sorting, cells were pooled together by tissue for three total captures and captured using the 10X Genomics Chromium Next GEM Single Cell 3ʹ Kit v3.1. scRNA-seq libraries were prepared following the manufacturer’s instructions and sequenced on a NovaSeq 6000 (Illumina).

### Single cell RNA analysis of primary 4T1 tumors

scRNA-seq data was processed with Cellranger as described above (“scRNA-seq data processing”). Filtered gene-cellBC matrices that contained only cellBCs with UMI counts that passed the threshold for cell detection were used for further analysis. Additional analysis was performed in R (v.4.2.3) using Seurat (*160*) (v.4.3.0) with default function parameters unless otherwise noted. Doublets were predicted using DoubletFinder (*161*) (v.2.0.3). Cell types were predicted using SingleR (*162*) (v.2.0.0) based on mouse bulk RNA-seq reference data (MouseRNAseqData) from celldex (v.1.8.0). Cells with <200 genes detected, >10% mitochondrial RNA content, or predicted doublets from DoubletFinder and cells annotated as Erythrocytes by SingleR were excluded from analysis. We normalized and identified variable features using the Seurat functions NormalizeData and FindVariableFeatures. We then regressed out percent mitochondrial RNA content and number of UMIs and scaled data with Seurat’s ScaleData and ran PCA using variable features with RunPCA. Clusters were identified using shared-nearest-neighbor-based clustering based on the first 10 PCs with resolution = 0.2 with FindNeighbors and FindClusters. The same principal components were used to generate the UMAP projections with RunUMAP. Cell clustering for fine-resolution cell types was performed as described above for malignant/stromal, myeloid, and T cell subsets with the following parameters:

<u>Malignant/stromal:</u> dims = 1:10, resolution = 0.25
<u>Myeloid:</u> dims = 1:15, resolution = 0.25
<u>T cells:</u> dims = 1:10, resolution = 0.3

For cell subsets, cell cycle scoring was performed using Seurat’s CellCycleScoring and S phase/G2M phase scores were regressed out during data scaling. Cell types were assigned using SingleR annotations and manually refined based on expression of known marker genes.

### *In vivo* PEtracer sample generation

4T1 cells with high numbers of LTCv2 integrated lineage tracing cassettes and a 24-mer pegArray with protospacer mismatches designed to tune editing to ∼70% completion over a 4-to-6-week timescale were generated as described above (“Generation of 4T1 PEtracer cells edited to completion (fully-edited cells)”. These cells were infected with lentivirus expressing PEmax-T2A-GFP on day 0 of the experiment and sorted on day 2 for high mCherry signal, consistent BFP signal, and high GFP signal. Representative sorting gates shown in **fig. S14**. For mouse 1, these cells were expanded in culture for an additional week prior to a second sort on day 9 of the experiment. These cells were again expanded *in vitro* prior to injection into mice on day 12 of the experiment. For mouse 2 and 3, higher cell numbers were infected on day 0, an initial sort was performed on day 2 and the second sort was performed on day 5 prior to injection into animals on day 6.

Between 200,000 and 500,000 cells undergoing evolving tracing were injected into the tail vein of female Balbc/J x SELECTIV mice (mouse 1) or Balbc/J x [SELECTIV x CMV-Cre] mice (mouse 2 and mouse 3). Following injection, mice were monitored in accordance with IACUC guidelines prior to sacrifice either at a predetermined time point or when mouse health deteriorated to a point that humane sacrifice was necessary (mouse 1 = 39 days post-injection, mouse 2 = 28 days post-injection, mouse 3 = 35 days post-injection). Mice were sacrificed by CO_2_ asphyxiation prior to perfusion with 1% diethyl pyrocarbonate (DEPC) in sterile PBS. Following perfusion, mouse lungs were dissected out, incubated with DEPC in sterile PBS at 4°C for 3 hours, washed 2x with fresh 1x PBS, and soaked in optimal cutting compound (OCT) for 5-10 minutes at room temperature prior to degassing in a vacuum chamber. Degassed, OCT-embedded samples were frozen on dry ice and stored at −80°C until sectioning.

### *In vivo* MERFISH sample preparation

Frozen lungs were sectioned around −15 to −20°C on a cryostat (Leica CM3050s). Tissue slices were collected on silanized and PDL-treated 40 mm coverslips, and fixed by treating with freshly made 4% paraformaldehyde in 1x PBS for 15 minutes and were washed three times with 1x PBS and stored in 70% ethanol at 4°C for at least 18 hours to permeabilize cell membranes. Tissue sections that did not directly proceed to MERFISH hybridization were stored in 70% ethanol at 4°C for no longer than 2 months.

### Design of MERFISH encoding probes

To discriminate transcriptionally distinct cell types in 4T1 tumor and probe the related cell physiology, we assembled the 124-gene MERFISH panel from two categories:

1. Differentially expressed genes calculated from our annotated 10X scRNA-seq results. A Wilcoxon test was performed to identify the most significant differentially expressed genes.
2. Manually selected genes including canonical cell type markers.

For these candidate genes, we identified all possible 30nt probe binding sites within annotated transcripts and kept candidate probe sequences that satisfy the following criteria using the MERFISH probe designer (https://github.com/xingjiepan/MERFISH_probe_design): 68°C<Tm<83°C given the salt concentration in 2x SSC buffer; 40%<GC content<70%; no 15-mer match with rRNA and tRNA; transcriptome specificity >99%. After filtering probe binding sites, genes with fewer than 400 possible probe binding sites (roughly 500nt in transcript length) were removed.

For the selected genes, we designed a codebook by importing a covering design (https://ljcr.dmgordon.org/cover/links.html <u>with parameters: v=16 or 18, k=4, t=3)</u>, and then converting this design into a Hamming weight 4, Hamming distance 4 N-bit Hamming code (N=16 for the 124-gene MERFISH library, N=18 for the 175-gene MERFISH library) by removing codewords with less than 4 Hamming distance to other codewords. We then assigned the Hamming codes to genes by balancing the cell-type averaged UMI counts from reference scRNA-seq data across all bits and all cell types.

We then followed standard MERFISH library design procedures to generate the probe library. For each gene, we first assembled candidate probes by appending 3 readout sequences to the probe targeting sequences (reverse-complement of the probe targeting sequences designed above). These 3 readouts were randomly chosen from 4 assigned readouts assigned to this gene as indicated in the codebook and an extra “A” was added in between readout sequences and probe targeting sequences. Next, we appended 20nt forward and reverse primers to the 5’ and 3’ of these candidate probes. We then selected 90 to 96 of the overlapping probes for each gene among all possible candidate probes while minimizing the overlap binding to the target transcripts between neighboring probes. Then these fully assembled probes were further screened for cross-hybridization and any probe sharing >17nt of reverse-complement sequence with other probes was dropped. Full probe library sequences could be found in **table S21**.

To improve cell type resolution and monitor tumor progression we assembled the 175-gene MERFISH library from the following categories:

1. Genes from our initial 124-gene MERFISH library after removing genes with no obvious spatial or cell type specific enrichment (110 genes)
2. Cell cycle marker genes for identifying cell cycle states (7 genes)
3. EMT markers (5 genes)
4. Hypoxia markers (2 genes)
5. Additional FGF pathway markers (3 genes)
6. Finer cell type markers from existing references (34 genes)
7. Hits from a CRISPR screen in 4T1 cells (*122*) (4 genes)

Probes for this final set of selected genes were filtered with the same design criteria as the 124-gene library described above. For this 175-gene MERFISH library we used 18 choose 4 Hamming codes. Full probe library sequences could be found in **table S25**.

### MERFISH encoding probe hybridization and tissue clearing

Both *in vitro* and *in vivo* samples were washed once with 2x SSC prior to equilibration in encoding-probe wash buffer (30% formamide in 2x SSC supplemented with 0.1% Tween-20) for 10-15 minutes at room temperature. Encoding-probe wash buffer was then aspirated from the coverslip, and the coverslip was inverted onto a 50 µL droplet of probe mixture on a parafilm-covered 60 mm diameter petri-dish. The probe mixture comprised 1 nM of each encoding probe for MERFISH imaging, approximately 10 nM of each encoding probe for the two-color mCherry single-molecule FISH imaging, and 1 µM of a polyA-anchor probe (/5Acryd/TTGAGTGGATGGAGTGTAATT+TT+TT+TT+TT+TT+TT+TT+TT+TT+T, where T+ represents a locked nucleic acid T nucleobase, and /5Acryd/ is a 5′ acrydite modification available through IDT) in 2x SSC with 30% v/v formamide, 0.1% wt/v yeast tRNA (approximately, Life Technologies) and 10% v/v dextran sulfate (D8906, Sigma). We then incubated the sample at 37°C for 36-48 hours. The polyA-anchor probe was hybridized to polyA sequences on polyadenylated endogenous mRNAs, thereby anchoring these RNAs to the polyacrylamide gel. After hybridization, the samples were washed in the encoding-probe wash buffer for 30 minutes at 47°C for a total of two times to remove excess encoding probes and polyA-anchor probes. The samples were washed once with 2x SSC prior to poly-acrylamide gel-embedding.

Samples were embedded and tissue cleared similarly to section “Sample preparation for In-gel T7 amplification with fully-edited 4T1 cells” with the following modifications. Following tissue clearing, samples were washed in 2x SSC buffer supplemented with 0.2% (vol/vol) Rnase Inhibitor Murine (NEB) at room temperature for 2 hours. We changed the buffer every 20 minutes to ensure sufficient washing. Samples could be stored at 2x SSC supplemented with 0.2% Rnase Inhibitor Murine at 4°C for up to 2 weeks.

### MERFISH Imaging

MERFISH imaging was performed for each sample as described above in the section “Imaging of integration barcodes and lineage marks” with the following changes: two adaptors and two readout probes were used in each round of imaging; z-stack images consisting of 24 or 25 planes separated by 1.2 µm were collected for all channels; the number of imaging rounds was adjusted for each MERFISH library, specifically, for the 124-gene MERFISH library we used 16-bits (8 imaging rounds with 2 bits per round) and for the 175-gene MERFISH library we used 18-bits (9 imaging rounds with 2 bits per round). Four sections were imaged for mice 1 and 3, and 1 section was imaged for mouse 2. The number of fields of view and the set of tumors captured for each section is listed in **table S22.**

### MERFISH decoding

All MERFISH image analysis was performed using MERlin (available at https://github.com/zhengpuas47/MERlin) (*163*), as previously described (*155*). First, we identified the locations of fiducial beads in each FOV in each round of imaging and used these locations to determine the x–y drift in the stage position relative to the first round of imaging and to align images for each FOV across all imaging rounds. We then high-pass filtered the MERFISH image stacks for each FOV to remove background, performed ten rounds of Lucy-Richardson deconvolution to tighten RNA spots, and low-pass filtered them to account for small movements in the centroid of RNAs between imaging rounds. Individual RNA molecules imaged by MERFISH were identified by our previously-published pixel-based decoding algorithm using MERlin (*163*). After assigning barcodes to each pixel independently, we aggregated adjacent pixels that were assigned with the same barcodes into putative RNA molecules, and then filtered the list of putative RNA molecules to enrich for correctly identified transcripts as previously described for a gross barcode misidentification rate at 5% using MERlin.

### Nuclei segmentation for *in vivo* samples

Nuclei segmentation for *in vivo* samples was performed using Cellpose, following a similar approach to that used for *in vitro* nuclei segmentation (see “Nuclei segmentation for *in vitro* samples” section for reference) (*109*). However, modifications were necessary to accurately segment the small, densely packed nuclei observed *in vivo*. To enhance image sharpness and contrast of DAPI images before segmentation, we applied the Deconwolf deconvolution algorithm (*110*). Deconwolf was run for 100 iterations, with each field of view (FOV) divided into four tiles to facilitate computation on an Nvidia GeForce GTX 2080ti GPU. The resulting deconvolved images were segmented using Cellpose (v3.1.0) with nucleus diameter set to 17 pixels. Nuclei with a volume <100 μm³ were excluded from analysis. The number of nuclei detected per tumor section is listed in **table S22**.

### Assignment of MERFISH-decoded transcripts to cells

Using the 3-D coordinates from MERlin, transcripts were assigned to nuclei following the same procedure as for T7 amplicons (see “Assignment of T7 amplicons to cells”). However, rather than assigning cytoplasmic transcripts to the nearest nucleus, we used Proseg (v1.1.3) to infer the most likely cell boundaries based on the assumption that cytoplasmic gene expression mirrors that of the nucleus (*113*). Proseg was run separately for each tissue section with the following parameters: voxel-layers = 3, ncomponents = 10, nbglayers = 10, enforce-connectivity = True, max-transcript-nucleus-distance = 10, and nuclear-reassignment-prob = 0.2. The cells identified by Proseg were then mapped back to nuclear masks based on their assigned transcripts. Each Proseg cell was assigned to the nucleus with the highest Jaccard similarity, provided a similarity >0.4, which helped filter out spurious Proseg cell calls. Transcripts outside of nuclei were assigned using this mapping if the Proseg assignment probability was >0.5, while transcripts already within nuclei retained their original assignments. The total number of assigned transcripts and mean number of transcripts per cell for each tumor section is listed in **table S22**.

### MERFISH cell clustering and annotation

RNA count matrices were used for cell clustering and annotation with resolVI (*111*), a scvi-tools (*164*) (version 1.3.0) method for deconvolution and label transfer of spatial transcriptomics data. Separate models were trained for the 124-gene and 175-gene MERFISH libraries using sections M1 S3 and M3 S2, respectively. For model training, initial cell clustering was performed with Scanpy (v1.10.0) following standard preprocessing steps: cells with fewer than 5 genes or 20 transcripts were removed, data were normalized and log-transformed, total transcript counts were regressed out, and the data were scaled and reduced in dimensionality (20 PCs). Clusters were then identified using the Leiden algorithm (resolution = 0.6), and UMAP embeddings (min_dist = 0.25) were generated for visualization. Fine resolution clustering was achieved by re-clustering individual clusters with varying resolutions (0.1-0.4).

Labeled RNA count matrices were used as input for the resolVI model, which was trained for 100 epochs under a semi-supervised framework without downsampling counts. The trained model was used for reference mapping with the full dataset from all sections imaged using the same MERFISH library, with input data filtered, normalized, regressed, and scaled as described, and then projected. Cells were annotated by predicting labels and retaining those with a probability >0.5. Diffusion and background transcript proportions were calculated, and cells with >20% diffusion or background transcripts were removed. The latent representation from resolVI served as input for a final UMAP projection (min_dist = 0.2).

For cells profiled with the 175-gene MERFISH library cell cycle phase was classified using the “score_genes_cell_cycle” function implemented by Scanpy (v1.10.0). *Dscc1*, *Msh2*, *Rad51*, *Rpa2* and *Ung* were used to score S phase, while *Cdca2*, *Kif2c*, *Ncapd2*, *Nek2* were used to score G2/M.

### Comparison of MERFISH and scRNA-seq data

To assess MERFISH data quality, we compared correlation between MERFISH replicates as well as the correlation between MERFISH, bulk RNA-seq and scRNA-seq. For comparison between MERFISH replicates, we computed the average counts/cell for each transcript and calculated the Pearson correlation between two sections (M1 S1 and M1 S2). For comparison between MERFISH and bulk RNA-seq, we obtained normalized counts per million (CPM) data from published 4T1-derived lung metastasis bulk RNA-seq data (*114*) and calculated the Pearson correlation with average counts/cell obtained by MERFISH. To compare to scRNA-seq data described above (“Single cell RNA analysis of primary 4T1 tumors”), we computed the average normalized counts/cell per scRNA-seq or MERFISH cluster and calculated the pairwise Pearson correlation between cell types, including all cell types that were identified in both datasets.

### Cell type neighborhood enrichment and density analysis

Cell neighborhood enrichment analysis was performed using Squidpy (v1.6.2). First, spatial neighborhoods were computed by identifying the 6 nearest neighbors of each cell based on 2-D Euclidean distance, then neighborhood enrichment was assessed with the “nhood_enrichment” function.

Tumor and normal lung boundaries were delineated through a series of geometric operations and represented as polygons using geopandas (v1.0.1). To construct tumor boundaries, malignant cell centroids were dilated, and the outer edge of their combined area was determined. In contrast, normal lung boundaries were defined by subtracting the tumor boundary polygons from the union of dilated AT1/AT2 and Club cell centroids.

For each cell, distances to both tumor and lung boundaries were determined using the geopandas “sjoin_nearest” function. The distance to the lung/tumor interface was defined differently depending on cell location: for cells within the tumor boundary, it was the negative of the lung boundary distance; for cells outside the tumor boundary, it was the distance to the tumor boundary. These distances were scaled per tumor in each section by dividing by the maximum interface distance observed.

Cell type densities (cells per mm^2^) were determined by counting the total number of cells of each type within a 0.1 mm radius circle centered on each cell and dividing by the area of that circle. Rolling averages of these densities, computed along the gradient from normal lung to the outer tumor edge, were calculated using windows of 2,000 cells then scaled by dividing by the maximum average density for each cell type.

### FISH-based lineage readout after MERFISH for *in vivo* samples

After MERFISH imaging, samples were removed from FCS2 flow chamber, washed once with hybridization wash buffer (2x SSC and 30% v/v formamide), further digested, washed, and T7 *in situ* amplified as described in the“Sample preparation for In-gel T7 amplification” section. Samples were then imaged and analyzed as described above.

### Sample alignment between MERFISH and lineage imaging

To align samples on the home-built imaging platform for both MERFISH and lineage readouts, during MERFISH imaging we captured DAPI images using the 10x objective across entire tissue sections. During the setup of lineage imaging, we first adjusted the orientation of samples manually to roughly align their orientations with that used for MERFISH imaging. We then captured a new DAPI mosaic image for each complete tissue section using the 10x objective. Between the DAPI images captured for each sample during MERFISH and lineage imaging, we manually picked 10 positions as featured anchoring points and recorded their x-y coordinates. From these paired coordinates we performed a singular value decomposition (SVD) of the multiplicated position matrix to acquire a rotation and translation matrix by assuming samples only experienced affine transformation. Given the rotation and translation that we calculated, we applied the transformation to the center coordinates of the FOVs imaged during MERFISH for each sample and used these transformed coordinates as the new FOV coordinates for lineage imaging.

After image acquisition, we corrected residual alignment issues and sample distortions computationally. We generated large mosaic images by combining DAPI images from all fields of view acquired during the MERFISH and lineage readouts. These mosaics were downsampled by a factor of 4, smoothed with an unsharp mask (Gaussian sigma = 5 pixels), intensity-rescaled by clipping values >98th percentile, and rotated by an angle determined using anchoring points. Residual x-y drift was corrected by using phase cross-correlation to calculate the shift between the mosaics, and the lineage mosaic was adjusted accordingly. To address non-linear distortions, TV-

L1 optical flow (implemented in scikit-image v0.24.0) (*165*) was used to compute a vector field aligning the two mosaics. This vector field was divided into 100×100 pixel tiles, and for each tile the mean, intensity-weighted shift was calculated. The global x-y shift was then added to the local shift for each tile, and the tiles were saved. In experiments combining MERFISH and lineage imaging, nuclear masks were generated from the MERFISH images, and T7 amplicons were assigned to these masks after adjusting their 3-D coordinates based on the corresponding tile.

### Generation, preparation, and imaging of primary tumors with fully-edited 4T1 cells

Primary tumors were generated as above with fully-edited 4T1 cells used for *in vitro* validation experiments. 1,000,000 cells were subcutaneously injected into the mammary fat pad of female Balbc/J mice. Following injection, mice were monitored in accordance with IACUC guidelines prior to sacrifice at 2.5 weeks post-injection. Mice were sacrificed by CO_2_ asphyxiation and tumors collected.

Primary tumors embedded in OCT were prepared as in section “*In vivo* MERFISH sample preparation”; Specifically, the 20 μm sections were placed on treated coverslips and treated as MERFISH samples but skipping MERFISH imaging procedures. Samples were prepared for FISH-based lineage readout as described in the “Sample preparation for In-gel T7 amplification” section and then imaged and analyzed as described above. 146 FOVs were imaged, capturing the entirety of one primary tumor in one section.

### Processing of lineage data from *in vivo* tumor sections

T7 amplicons in lineage readout images were detected, quantified, and decoded as described above, with one key modification in the intBC decoding step. Instead of using a whitelist of all 2,171 possible intBCs, we limited it to the 36 intBCs found in the tumor-initiating clones. This smaller whitelist improved decoding under higher *in vivo* noise levels and allowed us to increase the maximum allowable distance between the codeword and the intensity profile to 2, thanks to the increased Hamming distances among the 36 intBCs.

LM decoding and quality control were carried out as described earlier, using a logistic regression classifier trained on fully-edited *in vitro* cells. For the primary tumor containing fully-edited cells, clones were identified based on their Jaccard similarity to intBCs and LMs in the fully-edited clone whitelist (see “Quantification of lineage tracing data accuracy in fully-edited cells” section for details). Cells with a Jaccard similarity <0.2 were not assigned to a clone, as they likely represent non-malignant cells lacking lineage tracing cassettes. LM decoding accuracy was then evaluated for malignant cells, serving as an out-of-sample validation of the classifier trained on *in vitro* data.

To account for imperfect nuclei segmentation and sample distortion between the MERFISH and lineage imaging, decoded intBCs were assigned to a whitelist of malignant cells derived from the MERFISH data. If the position of the decoded T7 amplicon was just outside of a malignant cell’s nuclear segmentation mask, it was assigned to that cell even if it was closer to a non-malignant cell, as long as the distance to the nuclear mask was less than 4 µm.

### Reconstruction of *in vivo* tumor phylogenies

Spatial data from all tissue sections for each mouse were manually aligned in the x-y plane using tissue morphology. Cells were then grouped into spatially distinct tumors using the DBSCAN clustering algorithm (ε = 0.3 mm), with abutting tumors separated manually. Clone calling, as described in the “Clone calling using intBCs” section, was performed for each tumor to determine its clonality and the corresponding set of intBCs. As with the colony data, spectral clustering was used for clone calling instead of NMF, due to the lower proportion of doublets.

Phylogenies for malignant cells were reconstructed using neighbor joining, as detailed in the “Distance-based tree reconstruction” section, incorporating data from all sections for each tumor, except for one mouse 3 section which was excluded because of low intBC detection. For tumor 2 in mouse 2 (M2 T2), separate phylogenies were built for the two contributing clones. To account for partial nuclei generated by tissue sectioning, only nuclei with centroids located more than 5 µm from the top or bottom of the section were used. Among these intact nuclei, the intBC detection rate was calculated, and only cells with a detection rate >60% were used for tree reconstruction. To improve tree reconstruction with missing data, edit sites with LMs shared by all cells were excluded, and select edit sites with LMs defining early splits were manually upweighted by a factor of 2 before calculating the Hamming distance matrix. Statistics for the 12 reconstructed tumor phylogenies are reported in **table S23**.

Further analysis was performed on phylogenies with edit fractions >50% (M1 T1, M1 T2, M1 T4, M2 T5, and M1 T1). This included inference of ancestral LM states and branch lengths as described in the “Inference of branch lengths” section and comparison of phylogenetic, character, and spatial distance as described in the “Comparison of pairwise phylogenetic, character, and spatial distance” section. To compute the LM installation efficiency and account for edits that occur at different depths of the tree, we computed the overall transition probability from the unedited state to a given lineage mark based on transitions along each branch in the tree. The resulting LM installation efficiency is normalized by tumor and edit site to account for differences in LM installation rate.

### Reconstruction of tumor growth dynamics

To reconstruct the tumor growth in 3-D we estimated the z-position of sections for mouse 1 tumor 1 (M1 T1) based on section number and thickness, and then inferred the position of each phylogenetic clade as the mean 3-D position of cells within that clade. The inferred tumor origin was determined by averaging the 3-D positions of all cells in the phylogeny, and the positions of ancestral nodes between the origin and each clade’s common ancestor were inferred by averaging the positions of their descendants with the inferred tumor origin.

To estimate the number of extant cells in each clade during the growth of M1 T1, we divided the timeline from day 0 to 39 into 20 discrete timepoints. At each timepoint, we cut the phylogenetic tree at the corresponding depth and counted the number of branches for each clade. These branches represent ancestral cells that were alive at that time point.

### Calculation of local LM diversity

We define local character diversity as the average pairwise LM distance between all cells within a 100 μm radius of a given cell. To compute this metric, we first identify spatial neighbors based on x-y coordinates and then extract the corresponding *M × M* subset from the *N × N* weighted Hamming distance matrix given the *M* nearest neighbors. The mean pairwise distance within this subset represents the local character diversity. Cells in phylogenetically diverse regions have higher local character diversity scores, as neighboring cells are less likely to share similar LMs.

### Estimation of cell fitness

Phylogenetic fitness was estimated using the infer_fitness function from the Jungle package (https://github.com/felixhorns/jungle) which implements the probabilistic fitness estimation algorithm described by Neher et al. 2014 (*166*). Conceptually, this algorithm leverages two key assumptions:

**1.** <u>Fitter cells divide more rapidly</u>, producing more offspring and leading to shorter branch lengths.
**2.** <u>Fitness changes gradually along lineages</u>, meaning a cell’s fitness is expected to be similar to that of its ancestors and relatives.

Phylogenetic fitness was calculated using the inferred branch lengths generated by ConvexML, as described in the “Branch Length Inference” section.

As an alternative tree-independent fitness metric, we calculated the mean neighbor LM distance for each cell in the phylogeny. Using the *N-by-N* weighted Hamming distance matrix, computed as described in the “Distance-Based Tree Reconstruction” section, we identified the 20 nearest neighbors of each cell and calculated the mean LM distance among them. This metric is based on the expectation that fitter cells will have closely related neighbors with similar LMs, as there has been less time for additional LMs to accumulate compared to less fit cells.

### Fitness correlation analysis

To identify gene expression patterns, spatial positions, and cell type neighborhoods associated with increased phylogenetic fitness, we computed the Pearson correlation between fitness and each feature for every tumor with fitness estimates. For gene expression, raw counts were used; for spatial positioning, we employed distance to the tumor boundary and the lung/tumor interface; and for cell type neighborhoods, we used local cell type density (see “Cell type neighborhood enrichment and density analysis” for details on boundary distance and density calculations).

Additionally, we calculated the Pearson correlation between these features and the distance to the tumor boundary to separate the effects of cellular features from those related to tumor position. P-values were derived using a two-sided Student’s t-test and adjusted for multiple comparisons with the Benjamini-Hochberg procedure. Results from this analysis are reported in **table S24**.

### Comparison of MERFISH libraries

To compare gene expression data generated by our 124- and 175-gene MERFISH libraries, we computed the average normalized counts/cell per annotated cluster across all samples profiled using each library and calculated the pairwise Pearson correlation between cell types, including all cell types that were identified using both libraries.

### Hotspot module analysis

Hotspot (v1.1.1) was used to call spatially coherent gene expression modules within the mouse 3 malignant cell population (*120*). Modules were identified using sample M3 S2. Spatially variable genes were selected based on Geary’s C autocorrelation (calculated with 100 neighbors), retaining genes with Geary’s C >0.05 and an average expression >0.1 counts per cell. Hotspot was then run with a minimum_gene_threshold=4 and fdr_threshold=0.01. For each M3 section, scores for the four modules identified in S2 were calculated using the same parameters. Each cell was assigned to the module with the highest score.

To identify cell types associated with each module, we calculated the mean density (see “Cell type neighborhood enrichment and density analysis” for reference) of each cell type across cells in each module, and then z-scored these values. For genes selected by Hotspot to distinguish between modules, we calculated their mean expression by aggregating normalized counts across cells in each module, and then z-scored these values.

Transition probabilities between Hotspot modules were calculated by first identifying the set of phylogenetic neighbors for each cell–those with a last common ancestor within 10 days. Then, for each module, we computed the fraction of these neighbors assigned to every other module, yielding the transition probabilities between modules.

### Moran’s *I* statistic for heritability

We use Moran’s *I* autocorrelation statistic to quantify the heritability of gene expression values and Hotspot module scores within a phylogeny. While Moran’s *I* is typically applied to spatial data to measure the clustering or dispersion of a variable, it can also be used in phylogenetics by replacing the spatial neighbor graph with a phylogenetic neighbor graph (*167*). Moran’s *I* is defined as:

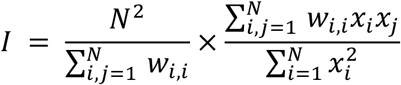

where *x* is the z-scored variable of interest, *w_i,j_* represents the connectivity between cells *i* and *j* (*w_i,jj_* = 1 for neighbors and 0 otherwise), and *N* is the number of cells. To construct the phylogenetic neighbor graph, each cell is connected to all other cells that share a LCA within the last 10 days. Moran’s *I* is then computed for z-scored MERFISH counts and Hotspot module scores. P-values are calculated with a permutation test (1,000 permutations) and adjusted for multiple comparisons with the Benjamini-Hochberg procedure (*168*). Results from this analysis are reported in **table S26**.

## Acknowledgments

We thank members of the Weissman, Yosef, and Zhuang labs for helpful discussions. We thank Jeffrey Moffitt for helpful discussions and advice on MERFISH sample preparation. We also thank the Whitehead Flow Cytometry Core, the Whitehead Genome Technology Core, the Koch Institute Hope Babette Tang Histology Facility, and the Broad Institute Klarman Cell Observatory.

## Author contributions

Conceptualization: LWK, KEY, JSW; Methodology: LWK, KEY, PZ, WNC, DY, RAS, WEA; Investigation: LWK, KEY, PZ, AK, JS, AS, DS; Software: WNC, PZ, MGJ, CE; Formal analysis: WNC, PZ, KEY, LWK; Visualization: WNC, LWK, KEY, PZ; Funding acquisition: JSW; Supervision: JSW, XZ, NY; Writing – original draft: LWK, KEY, WNC, PZ, JSW; Writing – review & editing: LWK, KEY, WNC, PZ, JSW, MGJ, DY, AK, JS, AS, DS, CE, RAS, XZ, WEA, NY

## Data and materials availability

Plasmids generated in this study will be available on Addgene. All scRNA sequencing data generated in this study are available under GEO accession GSE290975. All other sequencing data is available on SRA under BioProject accession PRJNA1231108. Additional processed data including MERFISH data and lineage reconstructions are available on Figshare (https://figshare.com/articles/dataset/PEtracer_Data/28473866, DOI 10.6084/m9.figshare.28473866). Custom code used in this study is available at https://github.com/jweissmanlab/PEtracer-2025.

